# Neural G0: a quiescent-like state found in neuroepithelial-derived cells and glioma

**DOI:** 10.1101/446344

**Authors:** Heather M. Feldman, Chad M. Toledo, Sonali Arora, Pia Hoellerbauer, Philip Corrin, Lucas Carter, Megan Kufeld, Hamid Bolouri, Ryan Basom, Jeffrey Delrow, José L. McFaline-Figueroa, Cole Trapnell, Steven M. Pollard, Anoop Patel, Christopher L. Plaisier, Patrick J. Paddison

## Abstract

Single cell RNA-seq has emerged as a powerful tool for resolving cellular states associated with normal and maligned developmental processes. Here, we used scRNA-seq to examine the cell cycle states of expanding human neural stem cells (hNSCs). From this data, we created a cell cycle classifier, which, in addition to traditional cell cycle phases, also identifies a putative quiescent-like state in neuroepithelial-derived cell types during mammalian neurogenesis and in gliomas. This state, Neural G0, is enriched for expression of quiescent NSC genes and other neurodevelopmental markers found in non-dividing neural progenitors. For gliomas, Neural G0 cell populations and gene expression is significantly associated with less aggressive tumors and extended patient survival. Genetic screens to identify modulators of Neural G0 revealed that knockout of genes associated with the Hippo/Yap and p53 pathways diminished Neural G0 *in vitro*, resulting in faster G1 transit, down regulation of quiescence-associated markers, and loss of Neural G0 gene expression. Thus, Neural G0 represents a dynamic quiescent-like state found in neuro-epithelial derived cells and gliomas.

## INTRODUCTION

Most developing and adult tissues are hierarchically organized such that tissue growth and maintenance is driven by the production of lineage-committed cells from populations of tissue-resident stem and progenitor cells (Reya et al, 2001). In adult tissues, stem cells are typically found in a quiescent or reversible G0 state and must re-enter the cell cycle and divide to promote lineage commitment (Doetsch, 2003; Obernier et al, 2018). Their progeny, e.g., amplifying progenitors, further balance lineage commitment with proliferation to produce adequate numbers of lineage committed and terminally differentiated cells to keep pace with demand (Lin, 2008). While much is known about specific regulatory events governing organismal development and tissue homeostasis, we lack a detailed picture of how cells enter, maintain, and exit quiescent-like states.

However, data from recent studies using single cell analysis of specific developmental compartments has begun to unravel some of the mysteries around G0-like states, including: hematopoiesis (Cabezas-Wallscheid et al, 2017; Hay et al, 2018), adult and fetal neurogenesis (Artegiani et al, 2017; Llorens-Bobadilla et al, 2015; Nowakowski et al, 2017), skeletal muscle regeneration (Scott et al, 2019), colon homeostasis (Grun et al, 2015), and a variety of other tissue types. The picture emerging from these studies indicates that in any given tissue, there is a continuum of highly regulated G0-like states in stem and progenitor cells and their progeny, which cause cells to enter long- or short-term states of quiescence (distinguishable from terminal differentiation/ maturation states). For example, during adult mammalian neurogenesis scRNA-seq analysis has led to a model where “dormant” quiescent neural stem cell (NSC) populations (e.g., in the subventricular zone or hippocampus) enter a “primed” state before entering the cell cycle and differentiating (Llorens-Bobadilla et al, 2015).

Use of scRNA-seq has also provided critical insight into intratumoral heterogeneity and developmental gene expression patterns for primary gliomas (Darmanis et al, 2017; Filbin et al, 2018; Neftel et al, 2019; Patel et al, 2014; Tirosh et al, 2016; Venteicher et al, 2017). One key conclusion from these studies is that each tumor represents a complex, yet maligned, neuro-developmental ecosystem, harboring diverse cell types, which presumably contribute to tumor growth and homeostasis in specific ways (e.g., vascular mimicry, immune evasion, recreating NSC niches, neural injury responses, etc.). However, these data sets have failed to produce models for transitions in and out of G0-likes states. In contrast to NSC scRNA-seq studies, where established cell-based markers are used to enrich for NSCs (e.g., GLAST+/Prom1+)(Llorens-Bobadilla et al, 2015), for GBM tumor cells there are no pre-existing universal markers that can neatly resolve subpopulations into quiescent, “primed”, G1, or differentiated cellular states (Lathia et al, 2015). As a result, these studies generally lump cells with “low cell cycle index” or low expression of genes primary expressed S/G2/M into a single G1 category.

Another underlying reason is that scRNA-seq cell cycle classifiers are not trained to identify G0-like populations. For example, the cell cycle classifier from the Seurat scRNA-seq analysis pipeline (ccSeurat), a gold standard, by design only allows classification of cells into G1, S, and G2/M phases (Butler et al, 2018). ccSeurat was trained on a mESC scRNA-seq data set, where mESCs were Hoescht stained and sorted into G1, S, and G2/M populations and then subjected to scRNA-seq (Buettner et al, 2015b; Scialdone et al, 2015). Since the training forced only these three states as the outcome (and mESCs do not transition into a natural state of quiescence), this classifier cannot identify G0-like states in somatic cells.

Here, we performed scRNA-seq on *in vitro* grown human neural stem cells (hNSCs) derived from the developing mammalian telencephalon (Davis & Temple, 1994; Johe et al, 1996), which can recapitulate the expansion, specification, and maturation of each of the major cell types in the mammalian central nervous system (Pollard et al, 2006; Sun et al, 2008). We have previously used hNSCs as non-transformed, tissue-appropriate controls for functional genomic screens in patient derived glioblastoma stem-like cells (GSCs) (Danovi et al, 2013; Ding et al, 2017; Ding et al, 2013; Hubert et al, 2013; Toledo et al, 2015; Toledo et al, 2014). We have observed that NCSs have longer doubling times of 40-50hrs compared to 30-40hrs for GSCs isolates, due to longer G0/G1 transit times (see below). NSC scRNA-seq analysis led to discovery of a transient Neural G0 subpopulation, which is enriched for genes expressed in quiescent NSCs and also a broader set of neurodevelopment markers expressed in other neural progenitors and cell types poised for cell division.

We then constructed a classifier, which we apply to neurodevelopment and glioma patient data to determine the functional impact of this cell subpopulation in real-world data. Finally, we identify genes that when perturbed diminish this G0-like state. Thus, our results reveal Neural G0 as a cellular state associated with quiescence in neuro-epithelial derived cell types.

## RESULTS

### Identification of cell cycle phases and candidate G0 and G1 subpopulations in human NSCs

We profiled 5,973 actively actively dividing U5-hNSCs (Bressan et al, 2017) using scRNA-seq to identify all the cellular gene expression states corresponding to cell cycle phases and specifically G0/G1 subpopulations (Methods). We then used unbiased cluster analysis to identify eight prominent clusters (Figs. 1A & B). One small cluster had cells with significantly lower RNA levels and has been included only as an outgroup for classifier construction (i.e., “other”). Meaning was attributed to the remaining seven clusters based on the set of marker genes significantly over-expressed within each cluster (avg log fold-change ≥ 0.3; adjusted p-value ≤ 0.05; Supplemental Fig. S1A; Supplemental Table S1). Based on these comparisions we labeled the clusters as follows (partly based on the Whitfield et al., 2002 convention): Neural G0 (17.3% of cells), G1 (36.7%), Late G1 (6.4%), S (7.2%), S/G2 (10.9%), G2/M (10.6%), and M/Early G1 (8.4%) (Fig. 1B).

**Fig. 1:**
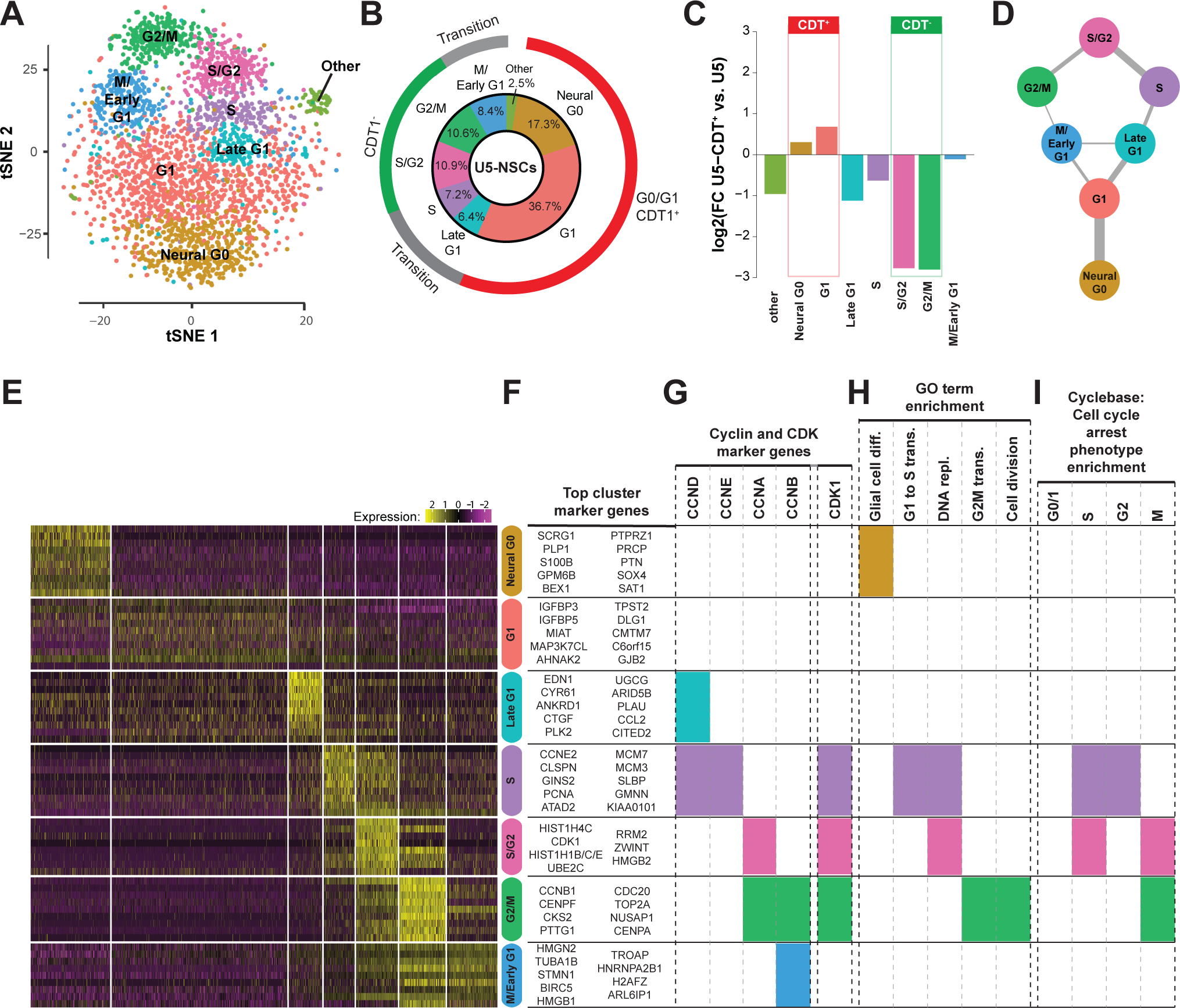
Gene Expression Map of Cell Cycle and Candidate G0 and G1 Subpopulations using Single Cell RNA-seq in hNSCs. **A,** Transcriptional clusters of unsorted U5-NSCs derived through an unbiased shared nearest neighbor (SNN) modularity and visualized through a t-Distributed Stochastic Neighbor Embedding (tSNE) comparison. **B,** Proportion of cells found in each cluster for the WT U5-NSCs. **C,** Fold-change between U5-hNSCs sorted for CDT+ compared to unsorted U5-hNSC cells on the log base 2 scale. Positive values indicate an increase in a given cell cycle subpopulation in CDT+ sorted relative to unsorted, and negative values indicate reduced cell subpopulations. The expected CDT^+^ cell subpopulations are found below the red bar, and the CDT^-^ cell subpopulations below the green bar. The others are expected to be transition subpopulations **D,** Network derived from Canberra distance between cluster medioids describes most likely connections between the cell cycle clusters and recapitulates the canonical cell cycle. **E,** Heat map of the relative expression (row-wise z-score) for the top 10 non-redundant genes for each prominent cluster in WT U5-NSCs and gene ontology analysis of the up-regulated genes defining each cluster. **F,** Top cluster marker genes. **G,** Cyclin and CDK marker genes found for each cluster. **H,** Function GO term enrichment for key cell cycle related and “glial cell differentiation” terms. Full cluster-defining gene list is in Supplemental Table S1 and full gene ontology **I,** Enrichment of knock-down cell cycle arrest phenotype genes in clusters.

As experimental validation we used the Fucci system (Sakaue-Sawano et al, 2008) to sort out CDT^+^ U5-hNSC subpopulation using a stably expressed mCherry gene fused to the ubiquitylation domains of human Cdt1. Importantly, the Late G1, S/G2, and G2/M were all significantly depleted (log2(FC) ≤ -1), whereas the Neural G0 and G1 subpopulations were enriched in the Cdt+ scRNA-seq populations (log2(FC) > 0; Fig 1C; Suppl. Fig. S1A). This validates that we have correctly defined the cell cycle cluster labels using an established system for detecting cell cycle state. It also demonstrates that the Neural G0 population is enriched when sorting out CDT+ cells, which validates that this subpopulation is a is a part of the G1/G0 pool of cells.

Network analysis of mean cluster gene expression resolved the trajectories of cells through the seven clusters into a pattern that fits well with cell cycle progression and predicted transit through G0/G1 (Figs. 1D & S2). Cells from the candidate G0 population were linked solely to the G1 cluster, which is consistent with G0 as a cell cycle exit from G1. The linkages between the clusters are not directed and thus the flow cells may pass in either direction. However, the model is consistent with results below in which indicate that cultured hNSCs enter G0-like state of variable length between M and S-phase. Importantly, this model of cell cycle progression was further validated by unique molecular identify (UMI) counts across clusters, where the counts start low in Neural G0 and peak in G2/M (Suppl. Fig. S1D). UMI counts can be viewed as an approximation of total mRNA expression in scRNA-seq data. Total mRNA expression during the cell cycle exactly follows this pattern, peaking with expression of Cyclin B and other mitotic genes (Shapiro, 1981).

There were four definable G0/G1 clusters: G1, M/Early G1, Late G1 and Neural G0. Despite being the largest cluster, the “G1” cluster had the smallest number of enriched genes, which included IGFR1 signaling genes (e.g., *IGFBP3* and *IGFBP5*), and significant reductions of genes expressed in S, S/G2, and G2/M clusters (Figs. 1E & S1C). The M/early G1 cluster showed low but significant residual expression of M phase genes and enrichment for splicing factor genes, which could represent residual mRNA from G2/M (Figs. 1E, S1C, & S2). The Late G1 cluster was defined by genes important in G1 cell cycle progression, including *CCND1* and *MYC*, and enriched for cholesterol biosynthesis, cell adhesion genes, and the subset of YAP target genes, such as *CTGF* and *SERPINE1* (Figs. 1E, S1B, S1C, & S2).

The Neural G0 cluster also showed significant repression of 246 genes peaking in other phases of cell cycle, including suppression of *CCND1* expression, which is an indicator of cell cycle exit (Sherr, 1995) and other cell cycle regulated genes such as *AURKB*, *CCNB1/2*, *CDC20*, *CDK1*, and *MKI67* (Figs. 1E & S1C). Moreover, the 158 up regulated genes defining this cluster were key genes with roles in neural development, including glial cell differentiation, neurogenesis, neuron differentiation, and oligodendrocyte differentiation (Figs. 1E, S1C, & S2; Supplementary Table SS). These genes included transcription factors with known roles in balancing stem cell identity and differentiation, including *BEX1, HEY1*, *HOPX*, *OLIG2, SOX2, SOX4*, and *SOX9* (Bergsland et al, 2006; Sakamoto et al, 2003; Scott et al, 2010) (Figs. 1E & S2).

The marker genes for each cluster was compared these marker genes for each cluster were analyzed for cyclin and CDK expression (Fig 1G), GO term functional enrichment (Fig 1H), and enrichment of genes that associated with specific cell cycle phases (Fig 1I)(Santos et al, 2015). Cyclin expression patterns are consistent with prior knowledge where CCND1 is a marker gene for the Late G1 and S phase clusters, CCNE2 is a marker gene for S phase cluster only, CCNA2 is a marker for the S/G2 and G2/M phase clusters, CCNA1 and CCNB1 for G2/M phase cluster only, and CCNB2 for G2/M and M/Early G1 phase clusters. In addition the CDK1 gene is a marker gene for the S, S/G2 and G2/M phase clusters. The cyclin and CDK1 expression pattern in the clusters is highly consistent with the expected cell cycle expression pattern (Darzynkiewicz et al, 1996). Functional enrichment analysis of each clusters marker genes linked Neural G0 with “glial cell differentiation”, S phase with “G1 to S transition”, S and S/G2 with “DNA replication”, and G2/M with “G2M transition” and “cell division” (Fig 1H). Gene knock-downs that arrest cells in S and G2 cell cycle phases were enriched in the S phase marker genes, arrest in S and M phase enriched in S/G2 phase marker genes, and arrest in M phase enriched in M phase marker genes (Fig 1I; (Santos et al, 2015)).

### Neural G0 is enriched in neuroepithelial-derived stem and progenitor cell populations

Comparison of our scRNA-seq cell clusters to gene expression profiles derived from *in vivo* neurogenesis samples supported our definition of the Neural G0 cluster. In two independent scRNA-seq analyses of adult rodent neurogenesis (Artegiani et al, 2017; Llorens-Bobadilla et al, 2015), the Neural G0 cluster showed most significant enrichment for genes defining quiescent neural stem cells and oligodendrocyte progenitor cells (Figs. 2A-D). These genes include, among others: *CLU*, *HOPX*, *ID3*, *OLIG2, PTN*, *SYT11*, *S100B*, *SOX9*, *PTPRZ1*, and *TTYH1* (Fig. 2B). Interestingly, for our S, S/G2, G2/M, and M/early G1 cluster genes, we found significant overlap with the activated NSCs of Llorens-Bobadilla et al. (2015) and the NPCs of Artegiani et al. (2017), which are no longer quiescent (Figs. 2C and 2D).

**Fig. 2:**
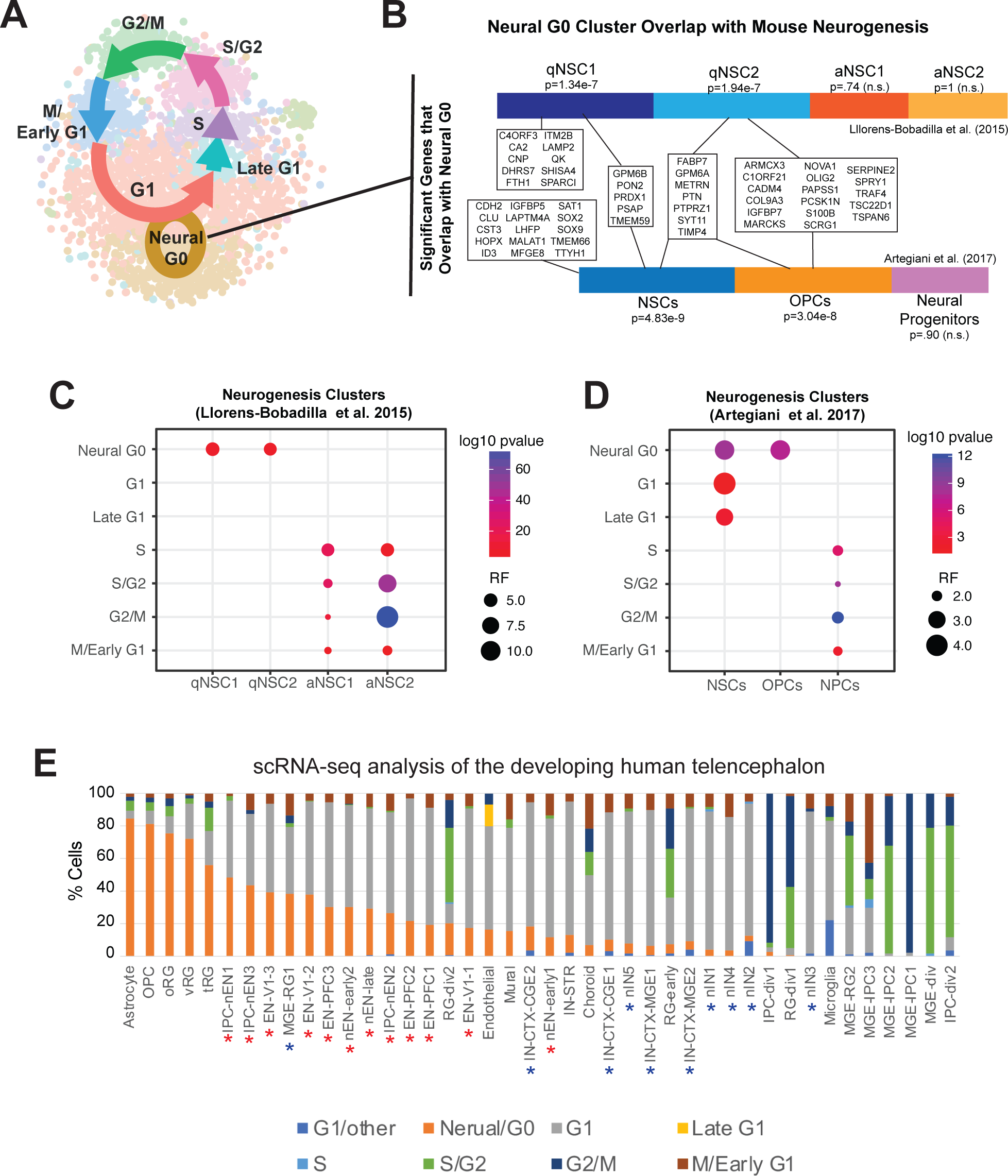
Comparison of hNSC cell cycle classifier with other neuroepithelial-derived cell populations. **A,** Model of the cell cycle of cultured hNSCs based on single cell transcriptomes. **B,** Overlap of the Neural G0 cluster with single cell transcriptomic profiles of quiescent neural stem cells (qNSCs) and activated (aNSC) NSCs/neural progenitors from adult rodent hippocampal niche (Llorens-Bobadilla et al. 2015; Artegiani et al. 2017). Significance assessed using hypergeometric analysis. OPC = oligodendrocyte progenitor. **C, D,** Significance of overlap of the neurogenesis cell subpopulation-defining genes in the early neurogenic lineage from two murine single cell RNA-sequencing studies compared to single cell cluster definitions (up-regulated genes) from unsorted U5-hNSCs grown in culture. Clusters presented in order of increasing activation with quiescent neural stem cells on the left and proliferating progenitors on the right. Significance assessed though hypergeometric analysis. RF = representation factor. **E**, Application of the hNSCs cell cycle classifier to scRNA-seq data from the developing human telencephalon (from (Nowakowski et al. 2017)). RG = radial glia, div = dividing. All cell type abbreviations are available in Table S3. Red asterisks indicate cells derived from the excitatory cortical neuron lineage, which originate from radial glial cells. Blue asterisks indicate cells from inhibitory cortical interneuron lineage.

To further investigate how Neural G0 might arise during mammalian development, we created a hNSC cell cycle classifier from our scRNA-seq data (dubbed ccAF, for <u>c</u>ell <u>c</u>ycle <u>A</u>SU/<u>F</u>red Hutch) (Methods) and applied into the developing human telencephalon. We analyzed scRNA-seq data from microdissected developing human cerebral cortex samples (PCW 5.85-19), which was previously used to analyze the spatial and temporal developmental trajectories for 11 cell types: astrocytes, oligodendrocyte precursor cells (OPC), microglia, radial glia (RG), intermediate progenitor cells, excitatory cortical neurons, ventral medial ganglionic eminence progenitors, inhibitory cortical interneurons, choroid plexus cells, mural cells, and endothelial cells (Nowakowski et al, 2017). We classified each single cell using our cell cycle categories and cross tabulated with the 11 cell types (Fig. 2E).

We found that the Neural G0 category was significantly enriched in non-dividing astrocytes, OPCs, and RGs (ventral, outer, and truncated), which had a Neural G0 population ranging from 85-72% (Fig. 2E; Supplemental Table S3). The signature diminishes in differentiating cells where G1 becomes the dominant category classification (Fig. 2E): excitatory cortical neuron lineage which originates from RGs, and the inhibitory cortical interneuron lineage which originate from MGE-RGs. We also observe a small but significant M/Early G1 subpopulation among differentiating cells, suggesting that it likely captures lineage committed cells that have just completed mitosis. Further, populations characterized as dividing (i.e., “div”, “div1”, or “div2”) are highly enriched with S/G2 and/or G2/M classified cells, and Neural G0 and G1 are absent or greatly diminished. Further, microglia, which arise from the embryonic mesoderm rather than neuroectoderm (Ginhoux & Garel, 2018), do not classify as harboring Neural G0 cells, but instead are classified as G1 and other.

Moreover, analysis of scRNA-seq of mouse embryonic stem cells (mESCs), representing blastocyst-stage pluripotent cells (i.e., pre-neuroepithelial cell), lacked cells from the Neural G0 subpopulation. For this analysis, we used scRNA-seq data from mESCs that were live sorted for DNA content via Hoechst staining into G1, S-phase, and G2/M populations (Buettner et al, 2015a). We found that our G1 category captured 83% of their Hoechst G1 cells, our G2/M category captured 89% of their G2/M, and their S-phase cells were split between G1, S, and G2/M, which is consistent with their Hoechst S-phase gate overlapping portions of these populations (Supplemental Fig. S3). However, the mESCs failed to classify into our Neural G0, Late G1, or M/early G1 categories. This is consistent with the shorter G1 of ESCs compared to somatic cells (Coronado et al, 2013).

These results suggest that the ccAF-defined Neural G0 classification identifies quiescent neuroepithelial-derived stem and progenitors as well also astrocytes, which may have progenitor-like properties, during fetal and adult neurogenesis.

### Validation of the ccAF classifier with other cell cycle data sets

To further examine the ccAF classifier we compared its performance with the current gold-standard cell cycle classifier from the Seurat scRNA-seq analysis pipeline (ccSeurat) (Butler et al, 2018) (https://satijalab.org/seurat/v3.1/cell_cycle_vignette.html).

The current ccSeurat classification system consists of only of G1, S, and G2/M phases. When applied to our hNSC scRNA-seq data, ccSeurat bins most of the cells called as Neural G0, G1, and M/Early G1 by ccAF into as G1, masking the ccAF subpopulations (Fig 1A & 3A). Overall there is good agreement between ccAF and ccSeurat when considering only the cells called as G1, S, or G2/M by ccAF (accuracy = 90%; Fig. 3B). Closer examination of cyclin expression across classified cell cycle phases reveals that the subdivision of the G1 ccSeurat category into Neural G0, G1, Late G1, and M/Early G1 by the ccAF is meaningful (Fig 3D-E). For example, ccAF identifies the Neural G0 subpopulation with low CCND1 gene expression (Fig 3D), a hallmark of quiescence (Sherr, 1995), whereas Seurat lumps together hi, medium, and low CCND1 expressing cells (Fig. 3E). In addition, ccAF better stratifies CCNA2 and CCNB1 expression across S, S/G2, G2/M, and M/Early G1, further suggesting that these phases are distinct (Fig. 3E). In support of this notion, examination of scRNA-seq data of proliferating HEK293 cells shows that ccAF classifies cells such that the expected cyclical pattern is observed, while ccSeurat misclassifies cells situated between S and G2/M as G1 (which ccAF classifies as S/G2) (Suppl. Fig. S4 A-B). And again, the cyclin and, also, CDK expression pattern better resolves the cell cycle with ccAF (Suppl. Fig. S4 C). Of note, ccAF does not classify any Neural G0 cells in the HEK293 kidney cells, which makes sense beacuase they of mesodermal, not neuroepithelial, origin.

**Fig. 3:**
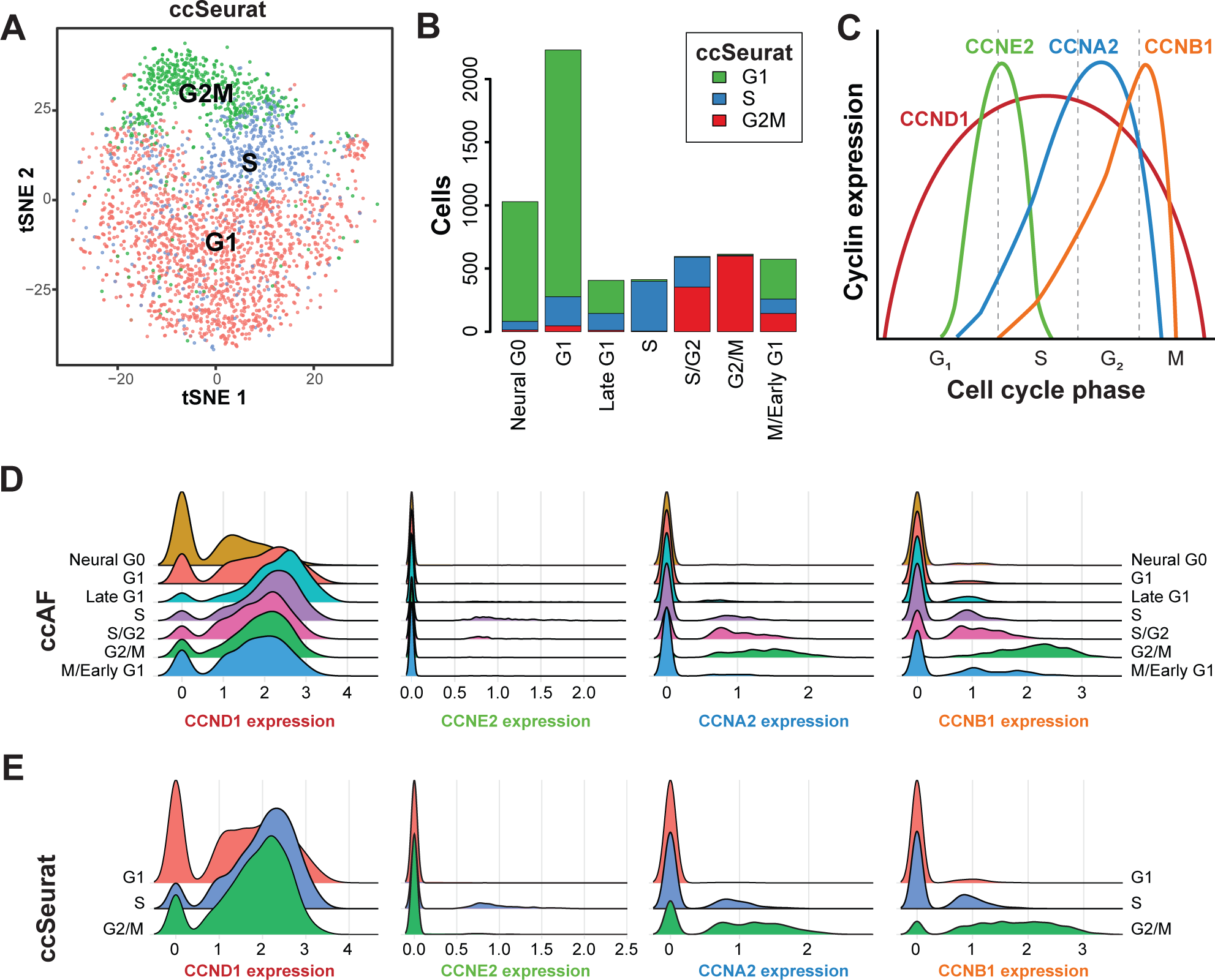
Comparison of ccAF and Seurat cell cycle classifiers for hNSC scRNA-seq data. **A,** tSNE plot displaying Seurat classified cell cycle phases overlaid on unsorted U5 cells. **B,** Number of cells for each ccAF cluster colorized by the proportion of cells that were called G1 (green), S (blue), and G2M (red) by Seurat’s built-in cell cycle classification approach. **C,** Expected pattern of cyclin expression during the mammalian cell cycle. **E,** Ridge graph comparisons of cyclin expression in ccAF-classified cell cycle phases. **F,** Ridge graph comparisons of cyclin expression in Seurat-classified cell cycle phases.

Taken together, these results illustrate that the ccAF classifier can be utilized as a cell cycle classifier for scRNA-seq data for neural and non-neural subtypes.

### Neural G0 is a prominent subpopulation in human glioma cells

Gliomas are tumors of the central nervous system which have a neuroepithelial cell of origin (Chen et al, 2012; Zong et al, 2015). They contain subpopulations of cells with stem cells-like characteristics that include expression of markers associated with NSCs, OPCs, and astrocytes, which may that may contribute to progression, therapy resistance, and tumor recurrence (Dirks, 2008; Zong et al, 2015). Recently, scRNA-seq has been applied to human gliomas of different grades and subtypes to reveal intratumoral cellular heterogeneity (Darmanis et al, 2017; Filbin et al, 2018; Neftel et al, 2019; Patel et al, 2014; Tirosh et al, 2016; Venteicher et al, 2017). To address whether Neural G0 also exists in gliomas, we used the ccAF classifier to analyze the scRNA-seq data available for 60 gliomas from these studies (Table 1; Fig. 4; Supplemental Table S3).

**Fig 4:**
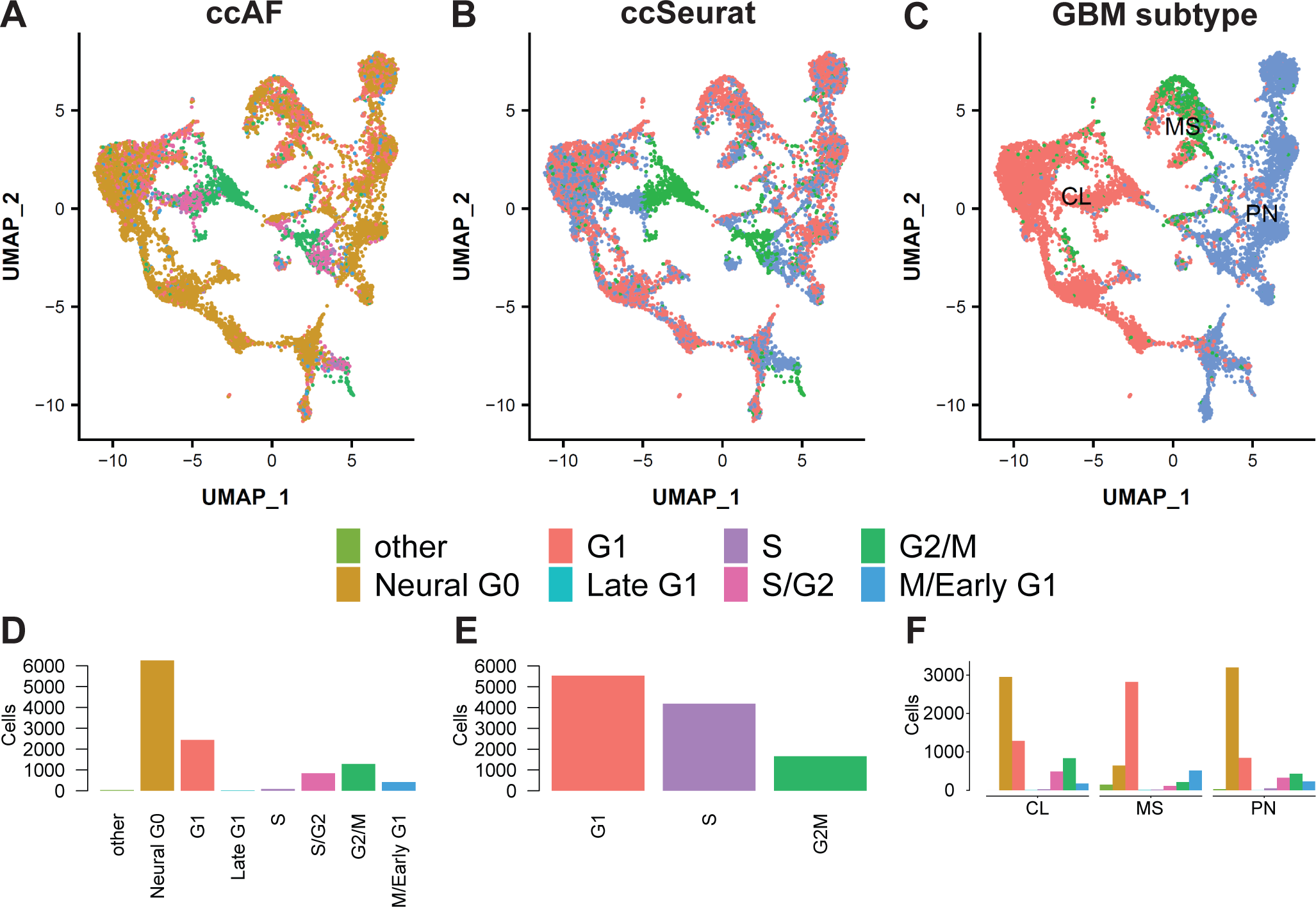
Comparison of ccAF and Seurat cell cycle classifiers for GBM scRNA-seq data from 22 tumors in Wang et al., 2019. **A,** UMAP plots of ccAF cell cycle classifications. **B,** UMAP plots of ccSeurat cell cycle classifications. **C,** UMAP plots of GBM subtype classifications. **D,** Cell count for ccAF cell type classifications. **C,** Cell count for ccSeurat cell type classifications. **D,** Cell count for each ccAF cell cycle phase broken down by GBM subtype.

**Fig. 5:**
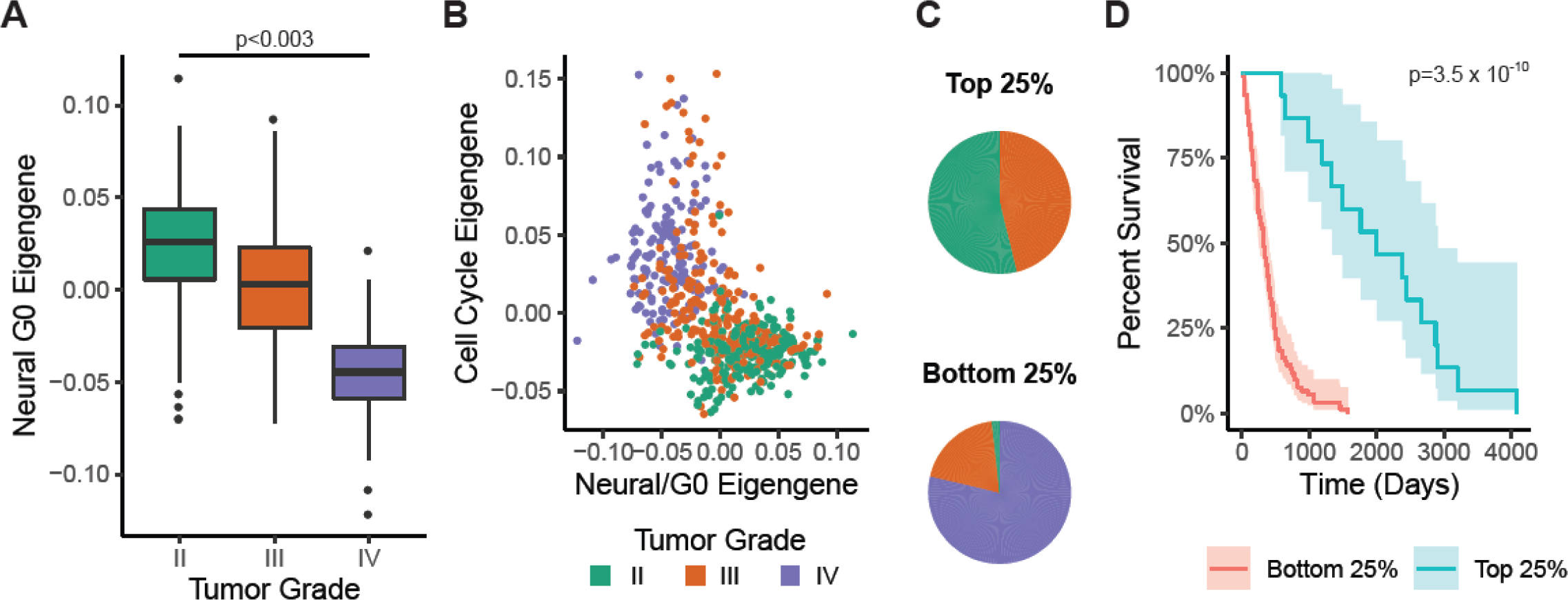
Neural G0 gene expression in 641 human gliomas. **A,** Relative Neural G0 eigengene expression between grade II, III, and IV tumors (TCGA; LGG and GBM). An eigengene represents the common variation across each patient tumor for the Neural G0 genes, i.e. first principal component corrected for direction if necessary. All pairwise Student’s t-tests comparisons had p-values <0.003. **B,** Comparison of cell cycle and Neural G0 eigengene expression in each glioma. Each tumor is colored by its grade (green = II, red = III, and purple = IV). **C,** Differences in the distribution of tumor grade between tumors with top 25% and bottom 25% of Neural G0 eigengene expression. **D,** Kaplan Meier survival plot of tumors with top 25% and bottom 25% of Neural G0 eigengene expression of Neural G0 genes. A Fleming-Harrington survival p-value was used to determine significance.

These tumors represent a broad range of gliomas, including: grades II, III, and IV, IDH1wt and mutant tumors, as well as glioma developmental subclasses (i.e., classical, mesenchymal, and proneural) and tumor types (i.e., astrocytoma, oligodendroglioma, GBM, and pediatric diffuse midline gliomas). Our analysis revealed that Neural G0 and G1 are the two most prominent tumor subpopulations regardless of stage (Table 1; Table S3). The Neural G0 and G1 represent 95.5% and 2.6%, respectively, of stage II oligodendrogliomas, 76% and 16.4% of stage III astrocytomas, 31-39% and 31-56% of stage IV GBMs, and 73.4% and 16.5% of diffuse midline gliomas (Table 1; Fig 4D). GBM subtype analysis (Wang et al, 2017) of each tumor cell further revealed that Neural G0 subpopulations showed strong bias against appearing in mesenchymal cell subpopulations in stage III and IV cancers (Fig 4C & F). Overall the prevalence of the Neural G0 state diminished as stage increased regardless of subtype (Table 1; Fig. 4C & F; Table S3).

Examining non-tumor brain cells types associated with stromal tissue available from Darminis et al., showed that Neural G0 populations could only be found in neuro-epithelial derived cells such as astrocytes, OPCs, and oligodendrocytes, whereas CD45+ cells and endothelial cells were negative. This was further evidenced by analysis of scRNA-seq data from 21 primary and metastatic head and neck cancers (Puram et al, 2017), where we observe that 80.3% of these tumor cells appeared in G1 but none contain a Neural G0 classified cell (Table 1).

Comparing ccAF to ccSeurat performance on these datasets revealed that ccAF-Neural G0 acounts for ∼46% of cells called by ccSeurat as G1 and resulting in a identification of G0-like states in across GBM patentient samples and cell subtypes (Fig. 4A-E).

Examination of scRNA-seq data for specific Neural G0 genes expressed in glioma revealed that 121 Neural G0 genes were significantly enriched in at least one data set (Suppl. Table S4). 12 genes, in particular, showed the strongest intersection between data sets (Suppl. Table S4; Suppl. Fig S5), including included EDNRB, FABP7, GPM6A, GMP6B, HEY1, PRDX1, PTPRZ1, SCD5, and TTYH1. Interestingly, these genes are preferentially expressed in GBM and LGGs compared to other cancers (Suppl. Fig. S6). Many have known or proposed roles in maintaining NSC/GSC “stemness” (EDNRB ((Liu et al, 2011)), PTPRZ1 ((Fujikawa et al, 2017)), TTYH1 (Kim et al, 2018; Wu et al, 2019)), slow cycling GBM cells (FABP7)(Hoang-Minh et al, 2018), possible neurogenic niche functions (e.g., GMP6B (Choi et al, 2013)), and neuroinflammation (PRDX1 (Kim et al, 2013) and PTN (Fernandez-Calle et al, 2017)).

We next determined whether the Neural G0 gene expression would be associated with bulk gene expression, genetic drivers, and survival data from 681 gliomas available in The Cancer Genome Atlas (TCGA; including both GBM and LGG). First, we calculated eigengenes for Neural G0 genes and a cell cycle genes (GO BP term Mitotic Cell Cycle = GO:0000278) that could be associated with the genetic drivers and patient survival. An eigengene represents the common variation across each patient tumor, i.e. first principal component corrected for direction if necessary. Figures 3A and 3B show that the Neural G0 eigengene is significantly down regulated as tumor grade increases. Neural G0 eigengene expression is significantly anti-correlated (R = - 0.58, p-value < 2.2 x 10^-16^) with cell cycle eigengene expression. Moreover, the Neural G0 and cell cycle eigengenes cell distribution demonstrate there is a lack of cells with high expression of both eigengenes, suggesting that the states are mutually exclusive (Fig. 3B).

To examine survival differences, we compared survival of patients with tumors exhibiting higher (top 25%) or lower (bottom 25%) Neural G0 gene expression (Figs. 3C and 3D). This analysis revealed a highly significant trend that tumors with higher Neural G0 expression survive on average 4.6 years longer than low Neural G0 expressing tumors (Fig. 3D). This difference likely driven by grade enrichment, where high Neural G0 tumors are exclusively grade II and III in the TCGA data set, while low tumors are mainly grade IV (Fig 3C), which have much worse survival (Claus et al, 2015; Stupp et al, 2005). Consistent with this notion, Neural G0 signature is also significantly associated with IDH1/2 mutation (Suppl. Fig. S7), which are primarily found in lower grade glioma (Claus et al, 2015; Yan et al, 2009). However, in a multivariate survival model the Neural G0 eigengene remains a significant predictor of overall survival even with the inclusion of the covariates (tumor grade and IDH1/2 mutation status), suggesting that the Neural G0 cell state is associated with patient survival variance independently from the common glioma survival associated covariates (tumor grade, IDH1/2).

Taken together, these results demonstrate that Neural G0 cells represent significant subpopulations in gliomas, which diminish by grade and are associated with better clinical outcomes. Thus, the results are consistent with a model whereby higher steady-state Neural G0 populations removes cells from the pool of cycling cells leading to slower tumor growth.

### CRISPR-Cas9 gene knockout screens identify regulators of Neural G0 *in vitro*

We next wished to investigate whether the Neural G0 state causes slower cell cycles. We reasoned that if Neural G0 ingress/egress is rate limiting for NSC cell cycles, diminishing Neural G0 would cause NSCs to cycle faster. If true, a simple pooled LV-CRISPR-Cas9 sgRNA library outgrowth screen in normal culture conditions should reveal overrepresented sgRNAs that cause diminished Neural G0 (Suppl. Fig. S8A).

We performed four separate CRISPR-Cas9 outgrowth screens, using three separate libraries, two different time points (10 days versus ∼3 weeks), and two different human NSC isolates, CB660 and U5 (Bressan et al, 2017; Pollard et al, 2006) (Suppl. Figs. S8 & S9; Suppl. Table S5). These screens revealed dozens of candidate screen hits significantly enriched at the end of outgrowth period (Suppl. Fig. S9A). These sgRNAs targeted genes found mutated across 35 different cancer (Suppl. Fig. S9C) and validated tumor suppressor genes (Futreal et al, 2004) (Suppl. Fig. S9D). Examining the intersection of all of our screen data revealed five reproducible and robust proliferation-enhancing screen hits: *CREBBP*, *NF2*, *PTPN14*, *TAOK1*, and *TP53* (Suppl. Fig. S8 & S9A & B), which we chose to validate further.

KO of *CREBBP, NF2, PTPN14, TAOK1,* and *TP53* in hNSCs caused a significant proliferative advantage over control cells in a 23-day outgrowth competition assay, while KO of the essential gene *KIF11* showed the opposite result (Suppl. Fig. S10A). However, the competitive advantage did not appear to be based on differences in survival since no changes in Annexin-V staining were observed following normal culturing or in co-cultures, where apoptosis remained <2% regardless of the experimental condition (data not shown).

Using cell proliferation assays (Suppl. Fig. S10B-D), we found that each KO significantly increased cell accumulation in 48-96 hour outgrowth assays. Importantly, this effect was independent of cell density, as KO cells showed increased proliferation at both low and high densities (Suppl. Fig. S10B). Further, the doubling time significantly decreased for each KO, shortening from ∼50 hours to 30-40 hours (Suppl. Fig S10E), similar to two GSC isolates used in the same assay.

### Neural G0 is reduced after KO of *CREBBP*, *NF2*, *PTPN14*, *TAOK1*, or *TP53* in NSCs

In order to further investigate changes in cell cycle dynamics, we utilized the fluorescent ubiquitination cell cycle indicator (FUCCI) system (Sakaue-Sawano et al, 2008). In normal culture conditions, ∼63% of U5-NSCs cells are in G0/G1, ∼15% are in S/G2/M, and the remainder are transitioning between these phases (Fig. 6A). KO of *CREBBP*, *NF2*, *PTPN14*, *TAOK1*, or *TP53*, however, caused a dramatic loss of the G0/G1 populations (reducing the frequency to 47-38%) and significantly lowered the ratio of G0/G1 to S/G2/M cells (∼2-4 fold lower) (Fig. 6B,C).

**Fig. 6:**
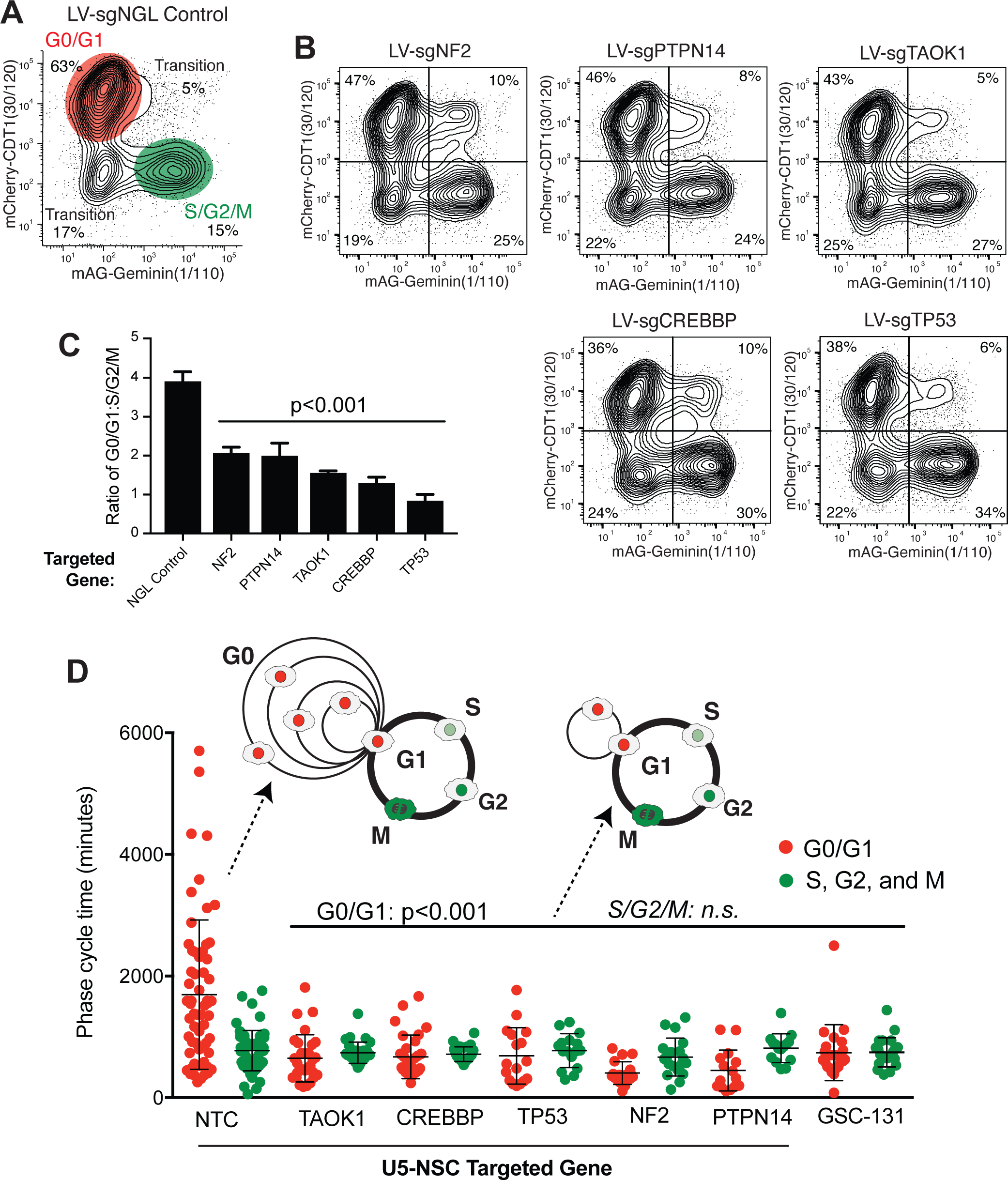
Reduction of G0/G1 Transit Time in NSCs after KO of *CREBBP*, *NF2*, *PTPN14*, *TAOK1*, or *TP53*. **A,** Representative contour plot of flow cytometry for Fucci (Sakaue-Sawano et al. 2008) in U5-NSCs after targeting of a non-growth limiting (NGL) control gene, *GNAS1*. Values are similar to wild-type and NTC U5-NSCs under similar culture conditions. The system relies on cell-cycle dependent degradation of fluorophores using the degrons from CDT1 (amino acids (aa) 30-120) (present in G0 and G1; mCherry) and geminin (aa1-110) (present in S, G2, and M; monomeric Azami-Green (mAG)). **B,** Representative contour maps of flow cytometry for Fucci following loss of *NF2*, *PTPN14*, *TAOK1*, *CREBBP*, and *TP53*. **C,** Ratio of G0/G1 (mCherry-CDT1+) to S/G2/M (mAG-Geminin+) from (A) and (B). Values are mean from 4 individually-tested LV guides per gene at 21 days post-selection. **D,** G0/G1 and S/G2/M transit times using time-lapse microscopy and Fucci. Differences in G0/G1 are statistically significant with p<0.0001 for targeted U5-NSCs and p=0.0006 for GSC-131 compared to NTC. The data are presented as the mean ± SD. Significance was assessed using a two-tailed student’s t-test (**C**) or Mann-Whitney test (**D**).

We also measured transit time through G0/G1 and S/G2/M in individual NSCs using time-lapse microscopy (Figs. 6D & S11). For G0/G1 transit times, we found that our control hNSCs exhibit variable G1 transit times and a wide distribution of G0/G1 transit times in control hNPCs, from fast (4.3 hrs), medium, and extremely slow (95 hrs) (averaging 32.5 hrs) (Fig. 6D). By contrast, S/G2/M transit times were much more uniform (∼12.4 hrs) (Fig. 6D). KO of *CREBBP*, *NF2*, *PTPN14*, *TAOK1*, or *TP53* dramatically collapsed the distributed G0/G1 transit times leading to a highly significant, faster transit of <11.7 hrs in KOs (p<0.0001) (Figs. 6D & S11). However, S/G2/M transit times were not significantly affected. GSCs also exhibit collapsed and faster G0/G1 transit times, similar to the KO hNSCs (Fig. 6D).

To further examine possible changes in G0/G1 dynamics, we examined molecular features associated with G0, G1, and late G1 (Suppl. Fig. S12A), including Rb phosphorylation, CDK2 activity, and p27 accumulation. In mammals, cell cycle ingress is governed by progressive phosphorylation of Rb by CDK4/6 and CDK2 as cells pass through the restriction point in late G1, causing de-repression of E2F transcription factors (Sherr & McCormick, 2002; Weinberg, 1995; Yao et al, 2008; Zetterberg et al, 1995). We observed that KO of *CREBBP*, *NF2*, *PTPN14*, *TAOK1*, or *TP53* in U5-NPCs results in a pronounced increase in the intensity of phosphorylated Rb during G1, consistent with an enrichment for a late G1 state (Suppl. Fig. S12B).

CDK2 activity correlates with cell cycle progression; if CDK2 activity levels are low during G1, cells enter G0 (Spencer et al, 2013). If CDK2 activity is intermediate (relative to its peak during G2/M), they progress past the restriction point and into S-phase (Spencer et al, 2013). Using the steady-state cytoplasmic to nuclear ratios of a DNA helicase B (DHB)-mVenus reporter as a readout of CDK2 activity (Hahn et al, 2009; Spencer et al, 2013), we observed significant increases in CDK2 activity in each KO in G0/G1 cells (Suppl. Figs. S12C,D). This was true either by total intensity or the proportion of cells with a reporter ratio greater than 1, a ratio which corresponds with the entrance to S-phase observed in mammary epithelium (Spencer et al, 2013). Control cells averaged ∼8% of G1 cells with >1 cytoplasmic:nuclear reporter ratios CDK2 activity, while KOs were 20-27% (Suppl. Fig. S12D).

Another hallmark of G0/quiescence is the stabilization of p27, a G1 cyclin-dependent kinase (CDK) inhibitor required for maintaining G0 (Coats et al, 1996; Susaki et al, 2007). Consistent with loss of transient G0 cells, we observed that KO of *CREBBP*, *NF2*, *PTPN14*, *TAOK1*, or *TP53* resulted in significant reduction of p27 levels in proliferating NSCs (Suppl. Fig. S12E,F).

Collectively, the above data demonstrate that KO of proliferation-limiting genes in U5-NSCs causes a cell autonomous decrease in cell cycle length with less distributed and faster G0/G1 transit times, an increase in the molecular features associated with late G1, and a reduction in the molecular features associated with G0 (Suppl. Fig. S12G). These data are consistent with KOs either blocking entry of cells into a transient G0 state or causing failure to maintain cells in G0. Therefore, we call these G0-skip genes.

### G0-skip mutants reprogram G0/G1, diminishing Neural G0 gene expression

To further characterize G0-skip genes, we performed gene expression analysis of KO cells specifically in G0/G1 phase. To this end, RNA-seq was performed on mCherry-CDT1+ sorted NSCs after KO, which captures both G0 and G1 subpopulations (Fig. 7A; Supplemental Table S6). In control NSCs, as expected, comparing G0/G1 sorted cells to unsorted populations revealed down-regulation of genes involved in cell cycle regulation, DNA replication, and mitosis (Fig. 7A; Suppl. Table S7). Overall comparisons between the KOs and NTC U5-NSCs showed that KO of *NF2* and *PTPN14* were most similar by unsupervised clustering as well as having the most overall gene changes, while *TAOK1* KO was most similar to the controls (Fig. 7B).

**Fig. 7:**
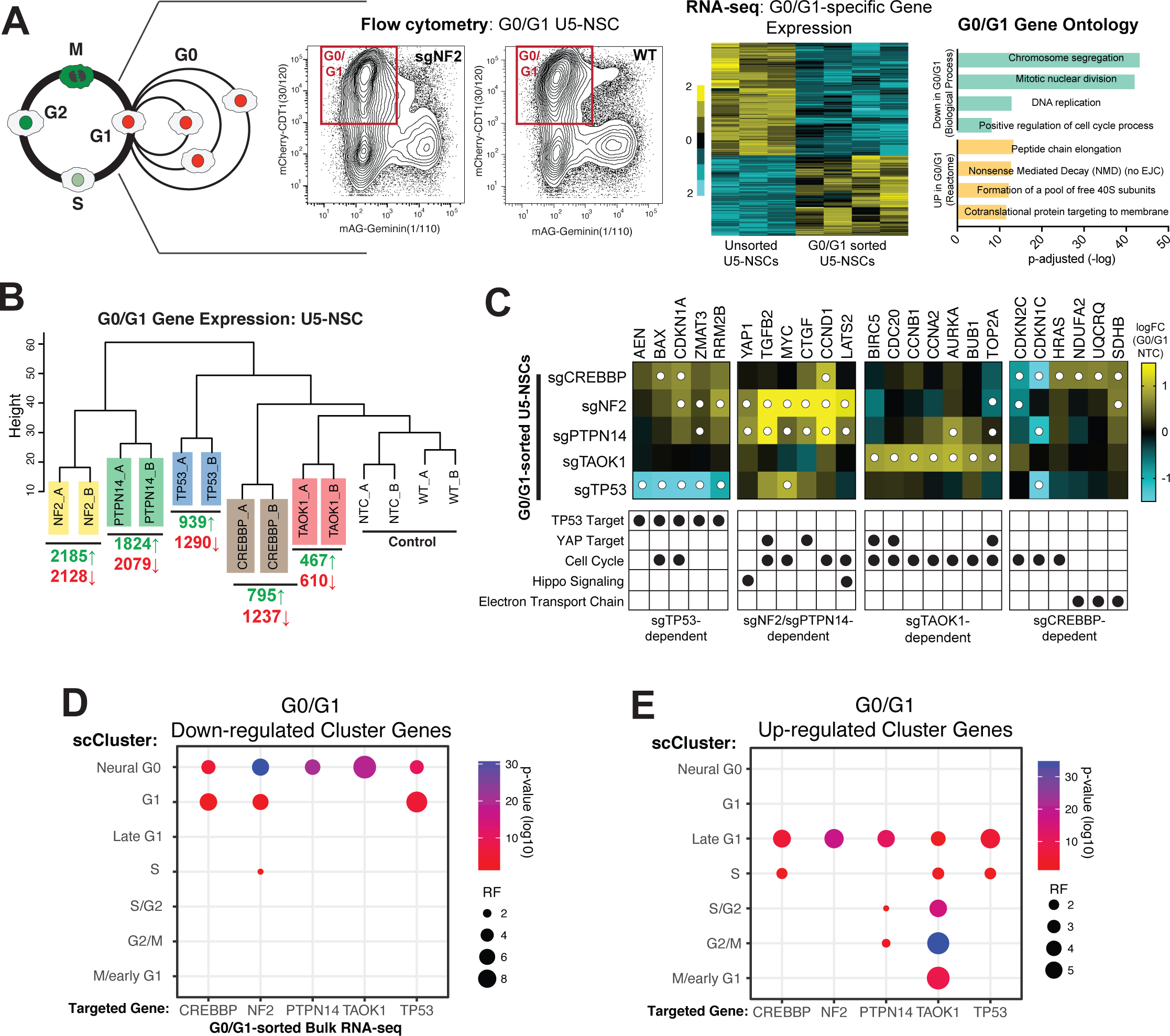
Transcriptional Reprogramming of G0/G1 Following Loss of G0-skip Genes. **A,** Schematic of G0/G1 sorting for gene expression analysis: mCherry-CDT1+ U5-NSCs (red box), heat maps of the significantly altered genes (FDR<0.05) between WT unsorted U5-NSCs and NTC and WT G0/G1 U5-NSCs, and gene ontology analysis (Young et al. 2010) of some of the top biological processes down-regulated and reactome groups (Yu and He 2016) up-regulated in the G0/G1 sorted cells. Full list in Supplementary Tables S10 & S11. **B,** Dendrogram of unbiased hierarchical clustering of gene expression from G0/G1-sorted U5-NSCs with the number genes up (green) and down (red) regulated (FDR<0.05) in each KO compared to NTC. Complete results in Supplementary Table S10. **C,** Heat map of log2FC compared to NTC for key genes changed in G0/G1 in following loss of *TP53, NF2/PTPN14*, *TAOK1*, and/or *CREBBP*, including genes from TP53 targets, YAP targets, the cell cycle, Hippo signaling, and electron transport genes. White dots indicate FDR<0.05. **D-E,** Significance of overlap of the down (D) and up (E) regulated genes from bulk RNA-sequencing of G0/G1 sorted cells with the single cell cluster definitions (up-regulated genes). Significance assessed though hypergeometric analysis. RF = representation factor.

However, comparison of the overlapping up- or down-regulated genes showed that *TAOK1* KO up-regulated genes were more similar to *NF2* and *PTPN14* KO than the other KOs (Suppl. Fig. S13A).

We next evaluated whether KO of the G0 skip genes were consistent with previously published and suggested roles in p53 pathway (for *TP53* and *CREBBP*) (Fischer, 2017; Ito et al, 2001) or the Hippo-YAP pathway signaling (for *NF2*, *PTPN14*, and *TAOK1*) (Lin et al, 2013; Plouffe et al, 2016; Wilson et al, 2014; Zhang et al, 2010). Evaluating p53 target genes, we found that only *TP53* KO significantly down-regulated the expression of high confidence p53 targets including: *BAX*, *CDKN1A/p21, RRM2B,* and *ZMAT3* (Fischer, 2017) (Figs. 7C & S13B). However, none of the other KOs showed inhibition of p53 targets or p53 itself, strongly suggesting that the other G0-skip genes are not acting through p53-dependent transcriptional activity.

Evaluation of 55 conserved HIPPO-YAP pathway transcriptional targets (Cordenonsi et al, 2011) revealed that each KO, except for *CREBBP*, showed significant enrichment for YAP targets with *NF2* KO having increased expression of the largest subset (Figs. 7C, S13C-E). Interestingly, *NF2* KO activated one subset of YAP targets important in the biological process of extracellular matrix (ECM) organization, while *TAOK1* KO activated a different subset of YAP targets important in nuclear chromosome segregation, such as during mitosis (Suppl. Fig. S13C-E). *NF2* and *PTPN14* KO shared the most overlap in YAP target activation, including targets considered universal Hippo-YAP targets (e.g., *CTGF*, *CYR61*, and *SERPINE1*).

We next used our ccAF classifier to determine whether genes associated with each phase change in G0/G1 populations in after KO of *CREBBP*, *NF2*, *PTPN14*, *TAOK1*, or *TP53*. We observed that Neural G0 were significantly down regulated in each KO (Fig. 7D & S14A), which included those expressed in quiescent NSCs and others cited above with key roles in neural development (e.g., *CLU*, *HOPX*, *ID3*, *PTN*, *PTPRZ1*, *SOX2*, and *SOX4*) (Suppl. Fig. S14B,C). By contrast, genes from late G1 cluster, including, for example, *CCND1* and *MYC*, were significantly up regulated in each KO, with *TAOK1* KO cells additionally showing increase in cell cycle phases as well (Fig. 7E & S15A-C). Examination of G0/G1 sorted populations from two GSC isolates (0131-mesenchymal and 0827-proneural) showed similar trends, with suppression of Neural G0 and G1 signature and higher expression of S and G2/M genes (Suppl. Fig. S16).

For NSC KOs, we also performed a more in-depth analysis of transcriptional changes of cell cycle genes and novel gene sets (Suppl. Fig. S17). These included cell cycle genes that could be causal for reprograming G0/G1 dynamics, such as up-regulation of G1 cyclins, E2F1/2 or down-regulation of CDKN1A/p21 and CDKN1B/p27 (Suppl. Fig. S17A). We also noted that for both *NF2* and *PTPN14* KO there was up-regulation of various Hippo-YAP pathway members, including *LATS2*, *TEAD1,* and *YAP1*, suggesting a possible feedback regulation of the pathway unique to *NF2* and *PTPN14* (Suppl. Fig. S17B) TAOK1 KO, in contrast to other KOs, strongly up-regulated >40 key regulators of mitosis (e.g., *AURKA*, *BUB1*, *CCNB1/2*, *CDK1*, *KIF11*, *etc.*), suggesting it may act to inhibit their precocious activation in G0/G1 or expression after completion of mitosis (Supplemental Fig. S14C).

*CREBBP* KO, uniquely among KOs, caused up-regulation of key nuclear-encoded mitochondrial genes, including members of the NADH dehydrogenase complex, the succinate dehydrogenase complex, and mitochondrial DNA polymerase (Suppl. Fig. S17D), which are direct transcriptional regulatory targets of nuclear respiratory factors 1 and 2 (NRF1 and NRF2) (Kelly & Scarpulla, 2004).

Finally, to more directly confirm reprograming of G0/G1 population in a G0-skip mutant, we performed scRNA-seq on G0/G1-sorted hNSCs with KO of *TAOK1* (Suppl, Fig. S18). The steady-state percentage of Neural G0 and, to a lesser degree, G1 cells in *TAOK1* KO cells is significantly diminished from 21.3% to 10.3% and 58.9% to 53.3%, respectively (Suppl. Fig. S18B,C). However, the late G1 population is increased (from 3.0% to 9.8%) as are cells in the M/early G1 (from 7.8% to 15.3%) and G2/M phase (from 1.5% to 4.4%). The expansion of the M/early G1 in *TAOK1* KO cells could explain the increase in mitotic genes observed in the bulk G0/G1 RNA-seq data in *TAOK1* KO cells (Suppl. Fig. S18C), suggesting that TAOK1 helps attenuate expression of mitotic genes from the previous cell cycle.

These results strongly suggest that NSC G0-skip mutants lose a significant fraction of Neural G0 subpopulation and reprogram G1 transcription networks to promote entry into G1-S.

## DISCUSSION

Using scRNA-seq from hNSCs, we identified clusters that represented cell cycle states G1, late G1, S, S/G2, G2/M, M/early G, and a new state, Neural G0. We built a classifier that accurately classifies new cells into these cell cycle states, and demonstrated that this classifier is accurate by applying it to synchronized bulk population data and scRNA-seq from HEK293T. Our classifier improves on the existing ccSeurat classifer, and can identify the new Nerual G0 that is a G0-like cellular state in hNSCs and other neuroepithelial-derived cell types. The identify of Neural G0 was determined by upregulation of genes associated with adult quiescent NSCs and neurodevelopment. We found that Neural G0 is a prominent subpopulation among non-dividing stem and progenitors, including OPCs and radial glial cells, which was diminished and replaced by G1 cells during differentiation. Analyzing scRNA-seq from human gliomas also revealed that Neural G0 is a significant non-dividing cell population, which is diminished as tumors become more aggressive and replaced by G1 cells.

Neural G0 appears to be restricted to neuroepithelial-derived cells, as we failed to find evidence for Neural G0 subpopulations in numerous non-neuroeptithelial cell populations (e.g., CD45+ cells). Finally, we observed in NSCs that Neural G0 can be ablated *in vitro* through genetic manipulation of at least 5 genes (*CREBBP*, *NF2*, *PTPN14*, *TAOK1*, or *TP53*), which causes dramatically faster G0/G1 transit times and loss of Neural G0-associated gene expression. Taken together, these results demonstrate that Neural G0 is a bona fide quiescent-like state for neuroepithelial cells *in vitro* and *in vivo* and that it has real-world implications for glioma patient survival.

The Neural G0 state appears exclusive to the neuroepithelial lineage (i.e., astrocytes, OPCs, RGs, and glioma cells). However, Neural G0 is not a singular state. Instead, each Neural G0 cell is enriched for a portion, but not all, of the 158 genes present in the hNSCs’ Neural G0, which helps distinguish it from G1 and other cell cycle phases. Thus, Neural G0 represents a mixed state that incorporates elements of qNSC and other neural progenitors, which likely results from the multipotency of fetal hNSCs combined with the effects of their ex vivo culture environment. G0-like states for non-neuroectoderm cells might be identified using an alternative set of developmental markers (e.g., Mesoderm G0).

With regard to the function, one possibility is that Neural G0 provides a compartment for maintenance of neurodevelopmental potential. That is, it could allow time for reinforcing transcriptional and epigenetic programs associated with neurodevelopment gene expression. Consistent with this possibility, Neural G0 genes are up regulated in quiescent NSCs *in vivo* and diminished during neural differentiation programs during corticogenesis or by KO of G0-skip genes in CDT+ NSCs. Moreover, multiple Neural G0 genes significantly enriched in NSCs and glioma Neural G0 cells are known to help maintain “stemness”. For example, HEY1 and TTYH1, are both are key players in Notch signaling pathway in NSCs and help maintain the NSC identity *in vivo* (Kim et al, 2018; Than-Trong et al, 2018). PTN and its target PTPRZ1 also may help promote stemness, signaling, and proliferation of neural progenitors and glioma tumor cells (Fujikawa et al, 2016; Fujikawa et al, 2017; Zhang et al, 2016b). Moreover, FABP7 expression and activity have been associated with lipid metabolism in slow-cycling GBM tumor cells (Hoang-Minh et al, 2018), consistent with Neural G0 state. Other functions for Neural G0 could include: time for repair of DNA lesions that persist from the previous cell cycle (Arora et al, 2017; Barr et al, 2017), oxidative stress/mitochondrial maintenance (Mohrin & Chen, 2016), or regulation of structural RNAs (e.g., rRNAs, tRNAs) (Roche et al, 2017). Future studies will be required to address these and other possibilities.

Our results have important implications for glioma biology. First, our classifier provides a method for identifying G0-like subpopulations in glioma tumor cells. While gliomas were among the first tumors dissected by scRNA-seq (Patel et al, 2014) and also for in depth genomic analysis (TCGA, 2008), computational analysis of scRNA-seq data and pathological examination of tumor samples has been up till now unable to distinguish G0 from G1 cells. However, our analysis suggests that these populations can be readily identified.

Second, we show that the proportion of Neural G0 cells in tumors correlates well with grade, patient survival, and proliferative state of gliomas. Outside of providing an important companion diagnostic to existing methods of grading gliomas, this analysis raises questions about the cellular composition of gliomas and the root causes of progression and responses to therapy. For example, our analysis of lower grade gliomas (LGG) suggests that they are could be “trapped” in Neural G0, where >93% of grade II cells categorized in Neural G0 (Table I). This would be consistent with Neural G0 acting as a barrier to progression in low grade gliomas, which is overcome in secondary gliomas.

**Table I:**
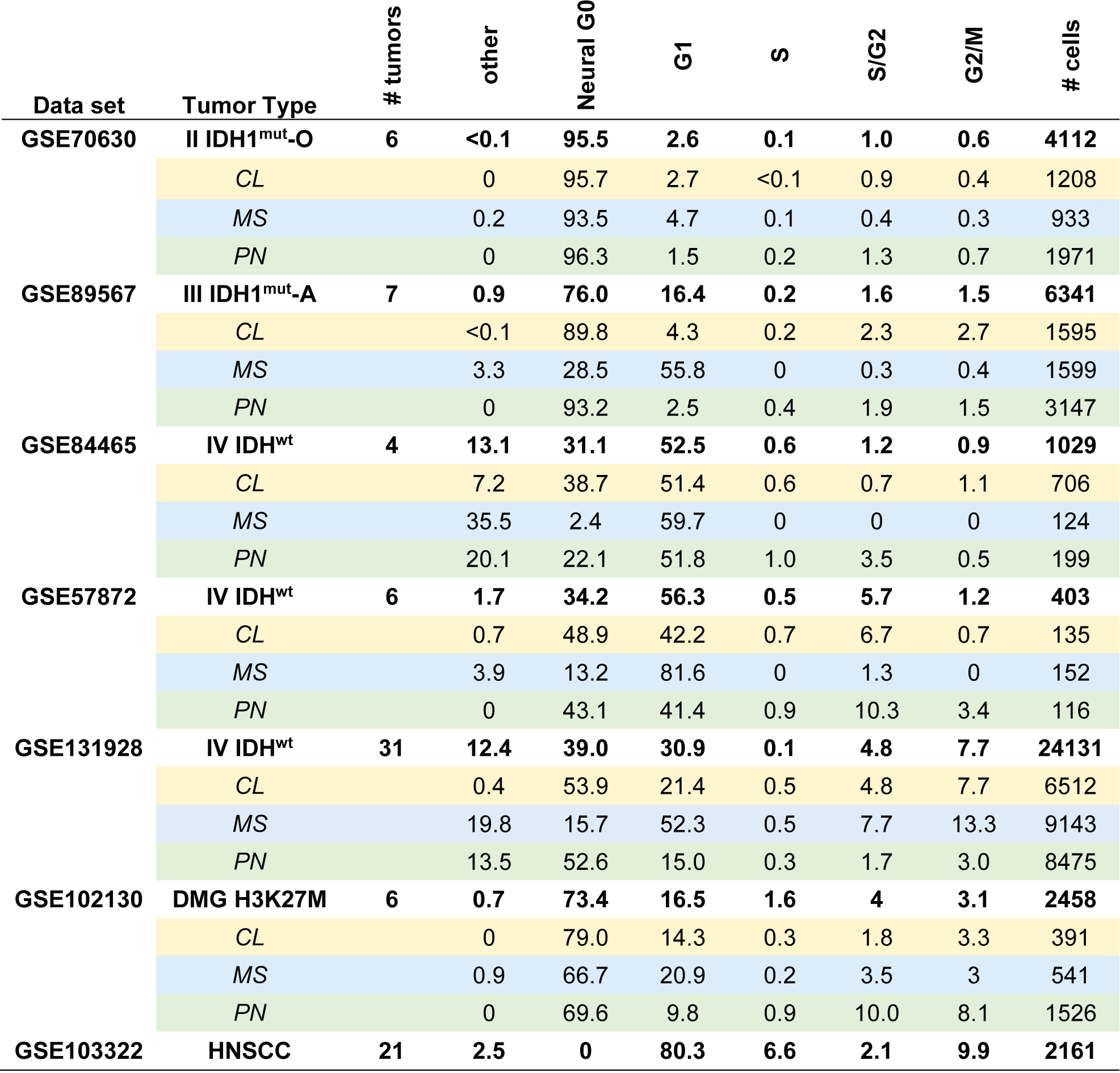
Percentage of glioma tumor cells defined by hNSCs cell cycle classifier. O = oligodendroglioma; A = astrocytoma; DMG = diffuse midline glioma; HNSCC = head and neck squamous cell carcinoma; CL = classical subtype; MS = mesenchymal subtype; PN = proneural subtype.

| Data set | Tumor Type | # tumors | other | Neural G0 | G1 | S | S/G2 | G2/M | # cells |
| --- | --- | --- | --- | --- | --- | --- | --- | --- | --- |
| GSE70630 | II IDH1<sup>mut</sup>-O | 6 | <0.1 | 95.5 | 2.6 | 0.1 | 1.0 | 0.6 | 4112 |
|  | CL |  | 0 | 95.7 | 2.7 | <0.1 | 0.9 | 0.4 | 1208 |
|  | MS |  | 0.2 | 93.5 | 4.7 | 0.1 | 0.4 | 0.3 | 933 |
|  | PN |  | 0 | 96.3 | 1.5 | 0.2 | 1.3 | 0.7 | 1971 |
| GSE89567 | III IDH1<sup>mut</sup>-A | 7 | 0.9 | 76.0 | 16.4 | 0.2 | 1.6 | 1.5 | 6341 |
|  | CL |  | <0.1 | 89.8 | 4.3 | 0.2 | 2.3 | 2.7 | 1595 |
|  | MS |  | 3.3 | 28.5 | 55.8 | 0 | 0.3 | 0.4 | 1599 |
|  | PN |  | 0 | 93.2 | 2.5 | 0.4 | 1.9 | 1.5 | 3147 |
| GSE84465 | IV IDH<sup>wt</sup> | 4 | 13.1 | 31.1 | 52.5 | 0.6 | 1.2 | 0.9 | 1029 |
|  | CL |  | 7.2 | 38.7 | 51.4 | 0.6 | 0.7 | 1.1 | 706 |
|  | MS |  | 35.5 | 2.4 | 59.7 | 0 | 0 | 0 | 124 |
|  | PN |  | 20.1 | 22.1 | 51.8 | 1.0 | 3.5 | 0.5 | 199 |
| GSE57872 | IV IDH<sup>wt</sup> | 6 | 1.7 | 34.2 | 56.3 | 0.5 | 5.7 | 1.2 | 403 |
|  | CL |  | 0.7 | 48.9 | 42.2 | 0.7 | 6.7 | 0.7 | 135 |
|  | MS |  | 3.9 | 13.2 | 81.6 | 0 | 1.3 | 0 | 152 |
|  | PN |  | 0 | 43.1 | 41.4 | 0.9 | 10.3 | 3.4 | 116 |
| GSE131928 | IV IDH<sup>wt</sup> | 31 | 12.4 | 39.0 | 30.9 | 0.1 | 4.8 | 7.7 | 24131 |
|  | CL |  | 0.4 | 53.9 | 21.4 | 0.5 | 4.8 | 7.7 | 6512 |
|  | MS |  | 19.8 | 15.7 | 52.3 | 0.5 | 7.7 | 13.3 | 9143 |
|  | PN |  | 13.5 | 52.6 | 15.0 | 0.3 | 1.7 | 3.0 | 8475 |
| GSE102130 | DMG H3K27M | 6 | 0.7 | 73.4 | 16.5 | 1.6 | 4 | 3.1 | 2458 |
|  | CL |  | 0 | 79.0 | 14.3 | 0.3 | 1.8 | 3.3 | 391 |
|  | MS |  | 0.9 | 66.7 | 20.9 | 0.2 | 3.5 | 3 | 541 |
|  | PN |  | 0 | 69.6 | 9.8 | 0.9 | 10.0 | 8.1 | 1526 |
| GSE103322 | HNSCC | 21 | 2.5 | 0 | 80.3 | 6.6 | 2.1 | 9.9 | 2161 |

Lastly, we found that KO of five genes, *CREBBP*, *NF2*, *PTPN14*, *TAOK1*, or *TP53* diminish Neural G0 *in vitro* in hNSCs. Gene expression changes in G0/G1 populations of KOs confirmed reduction of Neural G0 genes and characteristic gene expression changes associated with the p53 transcriptional network, Hippo-YAP targets, cell cycle gene regulation, and many novel targets and pathways, including those downstream of CREBBP and TAOK1. Interestingly, in glioma, Hippo-Yap pathway activity has been shown to significantly increases with grade and is associated with poorer patient survival (Orr et al, 2011; Zhang et al, 2016a). Moreover, proneural tumors exhibit the lower Hippo-Yap pathway activity while mesenchymal tumors, the highest (Guichet et al, 2018; Orr et al, 2011). These data fit well with our results that Neural G0 populations are more prominent in lower grade gliomas and decreased in mesenchymal subpopulations in more aggressive GBM cells. However, it less clear whether p53 wouldl have a similar role in promoting G0-like states in tumors. *TP53* is among the most frequently altered genes in lower grade gliomas (26-74%) and in GBM (∼30%) tumors (TCGA data; cbioportal) and there are many examples of p53 independent pathways that regulate G0 ingress/egress in tumor contexts, e.g., (Brown et al, 2017; Chen et al, 2012). Consistent with this possibility, p27, but not p53-inducible p21, expression is significantly associated with longer term survival in gliomas (Kirla et al, 2003). Thus, *in vitro* in hNSCs, low level cellular stresses or DNA damage may trigger partial p53 activation and a transient p21-dependent G0-like state, as has been reported for other cell types (Spencer et al, 2013). Future studies will be required to address how these genes and pathways might effect G0-like states in NSCs and tumors.

Collectively, our data reveals Neural G0 is cellular state shared by multiple neural epithelial-derived stem and progenitor cell types, which likely plays key roles in neurogenesis and glioma tumor development and recurrence.

## METHODS

### Cell culture

NSC and GSC lines were grown in NeuroCult NS-A basal medium (StemCell Technologies) supplemented with B27 (Thermo Fisher), N2 (2x stock in Advanced DMEM/F-12 (Fisher) with 25 μg/mL insulin (Sigma), 100 μg/mL apo-Transferrin (Sigma), 6 ng/mL progesterone (Sigma), 16 μg/mL putrescine (Sigma), 30 nM sodium selenite (Sigma), and 50 μg/mL bovine serum albumin (Sigma), and EGF and FGF-2 (20ng/mL each) (Peprotech) on laminin (Sigma or Trevigen) coated polystyrene plates and passaged according to previously published protocols(Pollard et al, 2009). Cells were detached from their plates using Accutase (Thermo Fisher). 293T (ATCC) cells were grown in 10% FBS/DMEM (Invitrogen).

### CRISPR-Cas9 screening

For large-scale transduction, NSC cells were plated into T225 flasks at an appropriate density such that each replicate had 250-500-fold representation, using the two previously published CRISPR-Cas9 libraries (Doench et al, 2016; Shalem et al, 2014) (Addgene) or a custom synthesized sgRNA library (Twist Biosciences) targeting 1377 genes derived from(Toledo et al, 2015). NPCs and GSCs were infected at MOI <1 for all cell lines. Cells were infected for 48 hours followed by selection with 1-2 μg/mL (depending on the target cell type) of puromycin for 3 days. Post-selection, a portion of cells were harvested as Day 0 time point. The remaining cells were then passaged in T225 flasks maintaining 250-500-fold representation and cultured for an additional 21-23 days (∼10-15 cell doublings) or 10 days. Genomic DNA was extracted using QiaAmp Blood Purification Mini or Midi kit (Qiagen). A two-step PCR procedure was performed to amplify sgRNA sequence. For the first PCR, DNA was extracted from the number of cells equivalent to 250-500-fold representation (screen-dependent) for each replicate (2-4 replicates) and the entire sample was amplified for the guide region. For each sample, ∼100 separate PCR reactions (library and representation dependent) were performed with 1 μg genomic DNA in each reaction using Herculase II Fusion DNA Polymerase (Agilent) or Phusion High-Fidelity DNA Polymerase (Thermo Fisher). Afterwards, a set of second PCRs was performed to add on Illumina adaptors and to barcode samples, using 10-20ul of the product from the first PCR. Primer sequences are in Supplementary Table 8. We used a primer set to include both a variable 1-6 bp sequence to increase library complexity and 6 bp Illumina barcodes for multiplexing of different biological samples. The whole amplification was carried out with 12 cycles for the first PCR and 18 cycles for the second PCR to maintain linear amplification.

Resulting amplicons from the second PCR were column purified using Monarch PCR & DNA Cleanup Kit (New England Biolabs; NEB) to remove genomic DNA and first round PCR product. Purified products were quantified (Qubit 2.0 Fluorometer; Fisher), mixed, and sequenced using HiSeq 2500 (Illumina). Bowtie was used to align the sequenced reads to the guides(Langmead et al, 2009). The R/Bioconductor package edgeR was used to assess changes across various groups(Robinson et al, 2010). For the tiling library, only guides that mapped once to the genome and are within the gene’s coding region were considered for further analysis.

Raw and mapped data files are available at the Gene Expression Omnibus database (GSE117004).

### Individual lentiviral-sgRNA assembly for validation

For retests, individual or pooled sgRNA were cloned into lentiCRISPR v2 plasmid. Briefly, DNA oligonucleotides were synthesized with sgRNA sequence flanked by the following:

5’: tatatcttGTGGAAAGGACGAAACACCg

3’: gttttagagctaGAAAtagcaagttaa

PCR was then performed with the ArrayF and ArrayR primers (Supplementary Table 8). The PCR product was gel purified using the ZymoClean Gel DNA recovery kit (Zymo Research). Gibson Assembly Master Mix (NEB) was used to clone the PCR product into lentiCRISPR v2 plasmid(Sanjana et al, 2014). The ligated plasmid was then transformed into Stellar Competent cells (Clontech), and streaked onto LB agar plates. The resulting clones were grown up and sequence verified (GeneWiz).

### Lentiviral production

For virus production, lentiCRISPR v2 plasmids(Sanjana et al, 2014) were transfected using polyethylenimine (Polysciences) into 293T cells along with psPAX and pMD2.G packaging plasmids (Addgene) to produce lentivirus. To produce lentivirus for the whole-genome CRISPR-Cas9 libraries, 25x150mm plates of 293T cells were seeded at ∼15 million cells per plate. Fresh media was added 24 hours later and viral supernatant harvested 24 and 48 hours after that. For screening, virus was concentrated 1000x following ultracentrifugation at 6800*xg* for 20 hours. For validation, lentivirus was used unconcentrated at an MOI<1.

### Viability and Proliferation Assays

Cells were infected with lentiviral gene pools containing 3-4 sgRNAs per gene or with lentivirus containing a single sgRNA to the respective gene (Supplementary Table 8). Initial cell density was carefully controlled for in each experiment by counting cells using a Nucleocounter NC-100 (Eppendorf) and cells were always grown in subconfluent conditions. For viability assays, following selection, cells were outgrown for 7-10 days, then harvested, counted, and plated in triplicate onto 96-well plates coated with laminin in dilution format starting at 1,000 cells to 3,750 cells per well (cell density depended on cell isolate and duration of assay). Cells were fed with fresh medium every 3-4 days.

After 7-12 days under standard growth conditions, cell proliferative rates were measured using Alamar blue reagent according to manufacturer’s instructions (Invitrogen). For analysis, sgRNA-containing samples were normalized to their respective nontargeting control (NTC) samples. For doubling time assays, cells infected with individual sgRNAs or NTC were routinely cultured (split every 3-5 days), and counted at each split (Nucleocounter NC-100; Eppendorf). The overall growth of each well containing an individual sgRNA was calculated and compared to the NTC well.

Comparisons between multiple experiments were normalized.

### Competition experiment

NSCs were infected with lentiviral gene pools containing 3-4 sgRNAs per gene, puromycin selected, and mixed with NSCs infected with lentiviruses containing turboGFP at an approximate 1:9 ratio, respectively. Cultures were outgrown for 23 to 31 days and flow analysis (FACS Canto; Becton Dickinson) was conducted every 7-8 days for GFP expression. Flow analysis data was analyzed using FlowJo software. For each sample, the GFP-population for each time point was normalized to its respective Day 0 GFP-population and the NTC (competition index).

### Time-lapse microscopy

NPCs were infected with lentiviral gene pools containing 3-4 sgRNAs per gene or with individual sgRNAs, puromycin selected, outgrown for >13 days, and plated onto 96-well plates or 24-well plates. Plates were then inserted into the IncuCyte ZOOM (Essen BioScience), which was in an incubator set to normal culture conditions (37° and 5% CO2), and analyzed with its software. For the cell confluency experiment, phase images were taken every hour for 72 hours. For the FUCCI cell cycle experiment, images were taken every 10-15 minutes for 72-120 hours. Cell cycle transit time for G0/G1 (mCherry-CDT1(aa30-120)+) and S/G2/M (mAG-Geminin(aa1-110)+) was manually scored by three different observers in actively dividing cells (those that could be followed from mitosis to mitosis). Each KO was scored by at least 2 independent observers and consistency between scorers was checked through shared analysis of a standard.

### Western blotting

Cells were harvested, washed with PBS, and either immediately lysed or snap-frozen and stored at -80°C until lysis. Cells were lysed with modified RIPA buffer (150mM NaCl, 50mM Tris, pH 7.5, 2mM MgCl2, 0.1% SDS, 2mM DDT, 0.4% deoxycholate, 0.4% Triton X-100, 1X complete protease inhibitor cocktail (complete Mini EDTA-free, Roche) and 1U/μL benzonase nuclease (Novagen) at room temperature for 15 minutes. Cell lysates were quantified using Pierce 660nm protein assay reagent and proteins were loaded onto SDS-PAGE for western blot. The Trans-Blot Turbo transfer system (Bio-Rad) was used according to the manufacturer’s instructions. See Supplementary Table 8 for antibodies and dilutions. An Odyssey infrared imaging system was used to visualize blots (LI-COR) following the manufacturer’s instructions.

### Flow Cytometry

FUCCI constructs (RIKEN, gift from Dr. Atsushi Miyawaki) were transduced into wild-type U5-NPCs and sorted sequentially for the presence of mCherry-CDT1(aa30-120) and S/G2/M mAG-Geminin(aa1-110) on an FACSAria II (BD). Normal growth was verified post-sorting and then the FUCCI U5-NPCs were transduced with individual sgRNA-Cas9 (4 independent guides per gene) and selected with 1 μg/mL puromycin. Cells were grown out for 21 days with splitting every 3-4 days and maintaining equivalent densities. Cells were counted (Nucleocounter NC-100; Eppendorf) and plated 3 days before analysis on an LSR II (BD). Controls cultured in the same conditions included cells transduced with guides against 3 non-growth limiting genes, including *GNAS1*, and showed equivalent FUCCI ratios. Results were analyzed using FlowJo software.

### Immunofluorescence and CDK2 Activity

U5-NSCs were plated on acid-washed glass coverslips (phosphorylated Rb and CDK2 activity) or 96-well imaging plates (differentiation; Corning). They were fixed overnight in 2% paraformaldehyde (USB) at 4°C, washed with DPBS (with calcium and magnesium) (Fisher), and blocked and permeabilized with 5% goat serum (Millipore), 1% bovine serum albumin (Sigma), and 0.1% triton X-100 (Fisher) in DPBS for 45 minutes at room temperature. Samples were stained with primary antibody diluted in 5% goat serum in DPBS overnight at 4°C, washed with DPBS, and stained with secondary antibody (diluted 1:200 in 5% goat serum in DPBS) at 37°C for 45 minutes. See Supplementary Table 8 for antibodies and dilutions. Samples were washed with DPBS, dyed with 100 ng/mL 4’,6-diamidino-2-phenylindole (DAPI) diluted in DPBS for 20 minutes at room temperature, and washed with DPBS. Coverslips were preserved using ProLong Gold Antifade Mountant (Fisher) and inverted on glass slides. For differentiation, images were acquired on Nikon Eclipse Ti using NIS-Elements software (Nikon).

### Phosphorylated Rb and CDK2 Activity Image Analysis

Cells were transduced with mVenus-DNA helicase B (DHB) (amino acids 994– 1087)(Hahn et al, 2009) (gift from Dr. Sabrina Spencer) and the mCherry-*CDT1* FUCCI and sorted on a FACSAria II flow cytometer (BD). Cells were outgrown to ensure normal growth and then transduced with individual sgRNA-Cas9. After >10 days outgrowth, cells were counted and plated, grown for 2 days, and stained for phosphorylated Rb and imaged on a TISSUEFAXS microscope (TissueGnostics), 54 fields per KO or NTC.

Cells were analyzed using CellProfiler(Kamentsky et al, 2011). G0/G1 nuclei were identified by the presence of the *CDT1* FUCCI reporter (25-120 pixel diameter, Global/Otsu thresholding, and distinguishing clumped objects by shape). CDK2 activity was defined by the cytoplasmic to nuclear ratio of the mVenus-DHB reporter, with the cytoplasmic intensity of the DHB reporter defined as the upper quartile intensity of a 2-pixel ring around the CDT1-defined nucleus due to the irregular shape of the U5-NPCs.

### p27 reporter

The p27 reporter was constructed after (Oki et al., 2014), using a p27 allele that harbors two amino acid substitutions (F62A and F64A) that block binding to Cyclin/CDK complexes but do not interfere with its cell cycle-dependent proteolysis. This p27K^-^ allele was fused to mVenus to create p27K^-^-mVenus. To this end, the p27 allele and mVenus were synthesized as gBlocks (IDT) and cloned via Gibson assembly (NEB) into a modified pGIPz lentiviral expression vector (Open Biosystems). Lentivirally transduced cells were puromycin selected and validated using mCherry-CDT1 FUCCI and HDAC inhibitor treatment (48 hours of 5 μM apicidin (Cayman)) to induce G0/G1 arrest using FACS (LSR II from Becton Dickinson and FlowJo software).

### Bulk RNA sequencing expression analysis

For G0/G1 NSC, cells singly positive for mCherry-*CDT1* FUCCI were sorted on a FACSAria II (BD) directly into TRIzol reagent (Life Technologies). For differentiating cells, cells were sparsely plated and cultured with growth medium without EGF or FGF-2 for 7 days before being lysed with TRIzol reagent. For both, 2 replicates per condition were harvested. RNA was extracted using Direct-zol RNA MiniPrep Plus (Zymo Research). Total RNA integrity was checked and quantified using a 2200 TapeStation (Agilent). RNA-seq libraries were prepared using the KAPA Stranded mRNA-seq Kit with mRNA capture beads (KAPA Biosystems) according to the manufacturer’s guidelines. Library size distributions were validated using a 2200 TapeStation (Agilent). Additional library QC, blending of pooled indexed libraries, and cluster optimization was performed using the Qubit 2.0 Fluorometer (Fisher). RNA-seq libraries were pooled and sequencing was performed using an Illumina HiSeq 2500 in Rapid Run mode employing a paired-end, 50 base read length (PE50) sequencing strategy.

### Bulk RNA sequencing data analysis

RNA-seq reads were aligned to the UCSC mm10 assembly using Tophat2 (Trapnell et al, 2012) and counted for gene associations against the UCSC genes database with HTSeq (Anders et al, 2015). Differential expression analysis was performed using R/Bioconductor package edgeR (Robinson et al, 2010). Samples for G0/G1 bulk RNA-seq were collected in two batches, so batch-dependent genes were removed before analysis (inter-batch p-value<0.01 by Wilcoxon-Mann-Whitney). To ensure that no genes were eliminated that may be regulated specific to a particular knockout, genes with a CPM variability greater than 2-fold compared to the internal batch control and an expression greater than 1 CPM in at least one sample were retained. Differentially expressed genes (DEG) at the transcription level were found using a statistical cutoff of FDR < 0.05 and visualized using R/Bioconductor package pheatmap. Kolmogorov-Smirnov test were conducted in R using the function ks.test from stats package. Raw sequencing data and read count per gene data can be accessed at the NCBI Gene Expression Omnibus (GSE117004).

### Gene ontology analysis

Gene Ontology (GO)-based enrichment tests were implemented using GOseq (v 1.23.0)(Young et al, 2010), which corrects for gene length bias. Gene lists were also analyzed for pathways using the R/Bioconductor package ReactomePA (v 1.15.4)(Yu & He, 2016). Analysis used all genes either up or down-regulated with a FDR<0.05 compared to NTC. GO terms with adjusted P-values<0.05 were considered significantly enriched. Venn diagrams were generated on http://bioinformatics.psb.ugent.be/webtools/Venn/.

### Single cell RNA-sequencing Sample Preparation

Single cell RNA-sequencing was performed using 10x Genomics’ reagents, instruments, and protocols. Single cell RNA-Seq libraries were prepared using GemCode Single Cell 3’ Gel Bead and Library Kit. FUCCI U5-NSCs (both with and without lentiviral TAOK1 KO, >14 days outgrowth) were harvested and half the cells were sorted using the FACSAria II (BD) for cells singly positive for mCherry-CDT1 FUCCI. Sorted cells were kept on ice before suspensions were loaded on a GemCode Single Cell Instrument to generate single cell gel beads in emulsion (GEMs) (target recovery: 2500 cells). GEM-reverse transcription (RT) was performed in a C1000 Touch Thermal cycler (Bio-Rad) and after RT, GEMs were broken and the single strand cDNA cleaned up with DynaBeads (Fisher) and SPRIselect Reagent Kit (Beckman Coulter). cDNA was amplified, cleaned up and sheared to ∼200bp using a Covaris M220 system (Covaris).

Indexed sequencing libraries were constructed using the reagents in the GemCode Single Cell 3’ Library Kit, following these steps: 1) end repair and A-tailing; 2) adapter ligation; 3) post-ligation cleanup with SPRIselect; and 4) sample index PCR and cleanup. Library size distributions were validated for quality control using a 2200 TapeStation (Agilent). The barcoded sequencing libraries were quantified by a Qubit 2.0 Fluorometer (Fisher) and sequenced using HiSeq 2500 (Illumina) with the following read lengths: 98bp Read1, 14bp I7 Index, 8bp I5 Index and 10bp Read2. Sequencing data can be accessed at the NCBI Gene Expression Omnibus (GSE117004).

### scRNA-seq Analysis

CellRanger (10x Genomics) was used to align, quantify, and provide basic quality control metrics for the scRNA-seq data. Using Seurat version 2.3.0, the scRNA-seq data from wild-type U5 cells and sgTAOK1 knock-out cells were merged and analyzed. Both scRNA-seq data were loaded as counts, normalized, and then scaled while taking into account both percent of mitochondria and the number of UMIs per cell as covariates. The union of the top 1,000 most variant genes from each dataset were used in canonical correlation analysis (CCA) to merge the two datasets via alignment of their subspace. We then identified clusters of cells using a shared nearest neighbor (SNN) modularity optimization-based clustering algorithm. Marker genes for each cluster were identified as differentially expressed genes, and the determination of 8 clusters was based on the discovery of strong markers for 6 of the eight clusters (both the G1 and low RNA clusters did not have significantly upregulated marker genes). Identity of clusters was determined primarily through the expression of cyclins and cyclin-dependent kinases, and secondarily through the function of other marker genes. A tSNE visualization was generated with a perplexity setting of 23.

Network analysis was used to determine the trajectories of cells through the cell cycle. First, the cluster centroids (mean expression for each gene across all the cells from a cluster) were used to compute the Canberra distance measure. In a cycle like a cell cycle, it is expected that on average there would be 2 edges between each cell cycle state. A distance cutoff of 240 led to 2.28 connections per cluster was used to turn the distance matrix into a network (Futreal et al, 2004).

Network analysis of the clusters was performed using the STRING database (Szklarczyk et al, 2017) and visualized using Cytoscape software. Transcription factors were identified according to TFcheckpoint (Chawla et al, 2013).

### Training cell cycle ASU/Fred Hutch (ccAF) classifier

The top 257 most variant genes from the integrated U5 dataset were used to train a random forest classifier for all eight cell cycle states using Seurat’s function BuildRFClassifier which relies upon the ranger package. The ccAF classifier was then applied to properly normalized Seurat objects using the ClassifyCells function or to non-Seurat data using the predict.ranger function in R. Confusion matrices and classifier accuracy were calculated using the confusionMatrix function in the caret package in R.

### Hypergeometric Analysis and Representation Factor Calculations

Hypergeometric tests (Johnson et al, 2005) were carried out in R using function phyper (stat.ethz.ch/R-manual/R-devel/library/stats/html/Hypergeometric.html). Gene lists were pre-filtered for the shared genes in each analysis to get the total gene population size, (i.e., 2739 genes for single cell analysis that had greater than 3 counts per cell in at least 10 cells and removing batch-effected genes for G0/G1 bulk RNA-sequencing).

Representation factors were calculated according to (Kim et al, 2001). The representation factor shows whether genes from one list (list A) are enriched in another list (list B), assuming that genes behave independently.

### Statistics and Reproducibility

Data are presented as the mean or median ± standard deviation (SD) or standard error of the mean (SEM), as specified in the figure legends. Statistics were performed using GraphPad Prism 7.0 or analysis-specific functions in R. All statistical tests are specified in figure legends. The number of independent experiments is indicated in the figures, figure legends, or Methods.

## Supporting information

Supplmental Data Inventory

## Acknowledgements

We thank Jon Cooper, Eric Holland, and members of the Paddison lab for helpful discussions, and Atsushi Miyawaki and Sabrina Spencer for providing reagents. This work was supported by the following grants: Interdisciplinary Training in Cancer Fellowship NCI T32CA080416 (PH), NCI/NIH (R01CA190957; R21CA170722; P30CA15704) (PP), DoD Translational New Investigator Award CA100735 (PP), and the Pew Biomedical Scholars Program (PP).

## Author contributions

Project conception and design was carried out by P.J.P., C.L.P., H.M.F., and C.M.T. CRISPR-Cas9 screening was performed by H.M.F., C.M.T., and P.H.; hit validation was performed by H.M.F, C.M.T., P.H., and M.K.; critical reagents were generated by P.C. and L.C.; screen and RNA-seq data analysis and statistics was performed by S.A. with input from A.P.; scRNA-seq was performed by H.M.F. under supervision of J.L.M.-F. and C.T., and analyzed by C.L.P.; H.B. performed cancer mutation analysis; J.M. designed the tiling library; S.M.P. provided and validated the hNSCs; and P.J.P., H.M.F., A.P., and C.L.P. wrote the manuscript with input from all authors.

## REFERENCES

1. Anders S, Pyl PT, Huber W (2015) HTSeq--a Python framework to work with high-throughput sequencing data. Bioinformatics 31: 166–169

2. Arora M, Moser J, Phadke H, Basha AA, Spencer SL (2017) Endogenous Replication Stress in Mother Cells Leads to Quiescence of Daughter Cells. Cell Rep 19: 1351–1364

3. Artegiani B, Lyubimova A, Muraro M, van Es JH, van Oudenaarden A, Clevers H (2017) A Single-Cell RNA Sequencing Study Reveals Cellular and Molecular Dynamics of the Hippocampal Neurogenic Niche. Cell Rep 21: 3271–3284

4. Barr AR, Cooper S, Heldt FS, Butera F, Stoy H, Mansfeld J, Novak B, Bakal C (2017) DNA damage during S-phase mediates the proliferation-quiescence decision in the subsequent G1 via p21 expression. Nature communications 8: 14728

5. Bergsland M, Werme M, Malewicz M, Perlmann T, Muhr J (2006) The establishment of neuronal properties is controlled by Sox4 and Sox11. Genes Dev 20: 3475–3486

6. Bressan RB, Dewari PS, Kalantzaki M, Gangoso E, Matjusaitis M, Garcia-Diaz C, Blin C, Grant V, Bulstrode H, Gogolok S, Skarnes WC, Pollard SM (2017) Efficient CRISPR/Cas9-assisted gene targeting enables rapid and precise genetic manipulation of mammalian neural stem cells. Development 144: 635–648

7. Brown JA, Yonekubo Y, Hanson N, Sastre-Perona A, Basin A, Rytlewski JA, Dolgalev I, Meehan S, Tsirigos A, Beronja S, Schober M (2017) TGF-beta-Induced Quiescence Mediates Chemoresistance of Tumor-Propagating Cells in Squamous Cell Carcinoma. Cell Stem Cell 21: 650–664 e658

8. Buettner F, Natarajan KN, Casale FP, Proserpio V, Scialdone A, Theis FJ, Teichmann SA, Marioni JC, Stegle O (2015a) Computational analysis of cell-to-cell heterogeneity in single-cell RNA-sequencing data reveals hidden subpopulations of cells. Nat Biotechnol 33: 155–160

9. Buettner F, Natarajan KN, Casale FP, Proserpio V, Scialdone A, Theis FJ, Teichmann SA, Marioni JC, Stegle O (2015b) Computational analysis of cell-to-cell heterogeneity in single-cell RNA-sequencing data reveals hidden subpopulations of cells. Nat Biotechnol 33: 155–160

10. Butler A, Hoffman P, Smibert P, Papalexi E, Satija R (2018) Integrating single-cell transcriptomic data across different conditions, technologies, and species. Nat Biotechnol 36: 411–420

11. Cabezas-Wallscheid N, Buettner F, Sommerkamp P, Klimmeck D, Ladel L, Thalheimer FB, Pastor-Flores D, Roma LP, Renders S, Zeisberger P, Przybylla A, Schonberger K, Scognamiglio R, Altamura S, Florian CM, Fawaz M, Vonficht D, Tesio M, Collier P, Pavlinic D et al (2017) Vitamin A-Retinoic Acid Signaling Regulates Hematopoietic Stem Cell Dormancy. Cell 169: 807–823 e819

12. Chawla K, Tripathi S, Thommesen L, Laegreid A, Kuiper M (2013) TFcheckpoint: a curated compendium of specific DNA-binding RNA polymerase II transcription factors. Bioinformatics 29: 2519–2520

13. Chen J, Li Y, Yu TS, McKay RM, Burns DK, Kernie SG, Parada LF (2012) A restricted cell population propagates glioblastoma growth after chemotherapy. Nature 488: 522–526

14. Choi KM, Kim JY, Kim Y (2013) Distribution of the Immunoreactivity for Glycoprotein M6B in the Neurogenic Niche and Reactive Glia in the Injury Penumbra Following Traumatic Brain Injury in Mice. Exp Neurobiol 22: 277–282

15. Claus EB, Walsh KM, Wiencke JK, Molinaro AM, Wiemels JL, Schildkraut JM, Bondy ML, Berger M, Jenkins R, Wrensch M (2015) Survival and low-grade glioma: the emergence of genetic information. Neurosurg Focus 38: E6

16. Coats S, Flanagan WM, Nourse J, Roberts JM (1996) Requirement of p27Kip1 for restriction point control of the fibroblast cell cycle. Science 272: 877–880

17. Cordenonsi M, Zanconato F, Azzolin L, Forcato M, Rosato A, Frasson C, Inui M, Montagner M, Parenti AR, Poletti A, Daidone MG, Dupont S, Basso G, Bicciato S, Piccolo S (2011) The Hippo transducer TAZ confers cancer stem cell-related traits on breast cancer cells. Cell 147: 759–772

18. Coronado D, Godet M, Bourillot PY, Tapponnier Y, Bernat A, Petit M, Afanassieff M, Markossian S, Malashicheva A, Iacone R, Anastassiadis K, Savatier P (2013) A short G1 phase is an intrinsic determinant of naive embryonic stem cell pluripotency. Stem cell research 10: 118–131

19. Danovi D, Folarin A, Gogolok S, Ender C, Elbatsh AM, Engstrom PG, Stricker SH, Gagrica S, Georgian A, Yu D, U KP, Harvey KJ, Ferretti P, Paddison PJ, Preston JE, Abbott NJ, Bertone P, Smith A, Pollard SM (2013) A high-content small molecule screen identifies sensitivity of glioblastoma stem cells to inhibition of polo-like kinase 1. PLoS One 8: e77053

20. Darmanis S, Sloan SA, Croote D, Mignardi M, Chernikova S, Samghababi P, Zhang Y, Neff N, Kowarsky M, Caneda C, Li G, Chang SD, Connolly ID, Li Y, Barres BA, Gephart MH, Quake SR (2017) Single-Cell RNA-Seq Analysis of Infiltrating Neoplastic Cells at the Migrating Front of Human Glioblastoma. Cell Rep 21: 1399–1410

21. Darzynkiewicz Z, Gong J, Juan G, Ardelt B, Traganos F (1996) Cytometry of cyclin proteins. Cytometry 25: 1–13

22. Davis AA, Temple S (1994) A self-renewing multipotential stem cell in embryonic rat cerebral cortex. Nature 372: 263–266

23. Ding Y, Herman JA, Toledo CM, Lang JM, Corrin P, Girard EJ, Basom R, Delrow JJ, Olson JM, Paddison PJ (2017) ZNF131 suppresses centrosome fragmentation in glioblastoma stem-like cells through regulation of HAUS5. Oncotarget

24. Ding Y, Hubert CG, Herman J, Corrin P, Toledo CM, Skutt-Kakaria K, Vazquez J, Basom R, Zhang B, Risler JK, Pollard SM, Nam DH, Delrow JJ, Zhu J, Lee J, DeLuca J, Olson JM, Paddison PJ (2013) Cancer-Specific requirement for BUB1B/BUBR1 in human brain tumor isolates and genetically transformed cells. Cancer discovery 3: 198–211

25. Dirks PB (2008) Brain tumor stem cells: bringing order to the chaos of brain cancer. J Clin Oncol 26: 2916–2924

26. Doench JG, Fusi N, Sullender M, Hegde M, Vaimberg EW, Donovan KF, Smith I, Tothova Z, Wilen C, Orchard R, Virgin HW, Listgarten J, Root DE (2016) Optimized sgRNA design to maximize activity and minimize off-target effects of CRISPR-Cas9. Nat Biotechnol 34: 184–191

27. Doetsch F (2003) A niche for adult neural stem cells. Curr Opin Genet Dev 13: 543–550

28. Fernandez-Calle R, Vicente-Rodriguez M, Gramage E, Pita J, Perez-Garcia C, Ferrer-Alcon M, Uribarri M, Ramos MP, Herradon G (2017) Pleiotrophin regulates microglia-mediated neuroinflammation. J Neuroinflammation 14: 46

29. Filbin MG, Tirosh I, Hovestadt V, Shaw ML, Escalante LE, Mathewson ND, Neftel C, Frank N, Pelton K, Hebert CM, Haberler C, Yizhak K, Gojo J, Egervari K, Mount C, van Galen P, Bonal DM, Nguyen QD, Beck A, Sinai C et al (2018) Developmental and oncogenic programs in H3K27M gliomas dissected by single-cell RNA-seq. Science 360: 331–335

30. Fischer M (2017) Census and evaluation of p53 target genes. Oncogene 36: 3943–3956

31. Fujikawa A, Nagahira A, Sugawara H, Ishii K, Imajo S, Matsumoto M, Kuboyama K, Suzuki R, Tanga N, Noda M, Uchiyama S, Tomoo T, Ogata A, Masumura M, Noda M (2016) Small-molecule inhibition of PTPRZ reduces tumor growth in a rat model of glioblastoma. Scientific reports 6: 20473

32. Fujikawa A, Sugawara H, Tanaka T, Matsumoto M, Kuboyama K, Suzuki R, Tanga N, Ogata A, Masumura M, Noda M (2017) Targeting PTPRZ inhibits stem cell-like properties and tumorigenicity in glioblastoma cells. Scientific reports 7: 5609

33. Futreal PA, Coin L, Marshall M, Down T, Hubbard T, Wooster R, Rahman N, Stratton MR (2004) A census of human cancer genes. Nat Rev Cancer 4: 177–183

34. Ginhoux F, Garel S (2018) The mysterious origins of microglia. Nat Neurosci 21: 897–899

35. Grun D, Lyubimova A, Kester L, Wiebrands K, Basak O, Sasaki N, Clevers H, van Oudenaarden A (2015) Single-cell messenger RNA sequencing reveals rare intestinal cell types. Nature 525: 251–255

36. Guichet PO, Masliantsev K, Tachon G, Petropoulos C, Godet J, Larrieu D, Milin S, Wager M, Karayan-Tapon L (2018) Fatal correlation between YAP1 expression and glioma aggressiveness: clinical and molecular evidence. The Journal of pathology 246: 205–216

37. Hahn AT, Jones JT, Meyer T (2009) Quantitative analysis of cell cycle phase durations and PC12 differentiation using fluorescent biosensors. Cell cycle 8: 1044–1052

38. Hay SB, Ferchen K, Chetal K, Grimes HL, Salomonis N (2018) The Human Cell Atlas bone marrow single-cell interactive web portal. Exp Hematol 68: 51–61

39. Hoang-Minh LB, Siebzehnrubl FA, Yang C, Suzuki-Hatano S, Dajac K, Loche T, Andrews N, Schmoll Massari M, Patel J, Amin K, Vuong A, Jimenez-Pascual A, Kubilis P, Garrett TJ, Moneypenny C, Pacak CA, Huang J, Sayour EJ, Mitchell DA, Sarkisian MR et al (2018) Infiltrative and drug-resistant slow-cycling cells support metabolic heterogeneity in glioblastoma. EMBO J 37

40. Hubert CG, Bradley RK, Ding Y, Toledo CM, Herman J, Skutt-Kakaria K, Girard EJ, Davison J, Berndt J, Corrin P, Hardcastle J, Basom R, Delrow JJ, Webb T, Pollard SM, Lee J, Olson JM, Paddison PJ (2013) Genome-wide RNAi screens in human brain tumor isolates reveal a novel viability requirement for PHF5A. Genes Dev 27: 1032–1045

41. Ito A, Lai CH, Zhao X, Saito S, Hamilton MH, Appella E, Yao TP (2001) p300/CBP-mediated p53 acetylation is commonly induced by p53-activating agents and inhibited by MDM2. EMBO J 20: 1331–1340

42. Johe KK, Hazel TG, Muller T, Dugich-Djordjevic MM, McKay RD (1996) Single factors direct the differentiation of stem cells from the fetal and adult central nervous system. Genes Dev 10: 3129–3140

43. Johnson NL, Kemp AW, Kotz S (2005) Univariate discrete distributions, 3rd edn. Hoboken, N.J.: Wiley.

44. Kamentsky L, Jones TR, Fraser A, Bray MA, Logan DJ, Madden KL, Ljosa V, Rueden C, Eliceiri KW, Carpenter AE (2011) Improved structure, function and compatibility for CellProfiler: modular high-throughput image analysis software. Bioinformatics 27: 1179–1180

45. Kelly DP, Scarpulla RC (2004) Transcriptional regulatory circuits controlling mitochondrial biogenesis and function. Genes Dev 18: 357–368

46. Kim J, Han D, Byun SH, Kwon M, Cho JY, Pleasure SJ, Yoon K (2018) Ttyh1 regulates embryonic neural stem cell properties by enhancing the Notch signaling pathway. EMBO Rep 19

47. Kim SK, Lund J, Kiraly M, Duke K, Jiang M, Stuart JM, Eizinger A, Wylie BN, Davidson GS (2001) A gene expression map for Caenorhabditis elegans. Science 293: 2087–2092

48. Kim SU, Park YH, Min JS, Sun HN, Han YH, Hua JM, Lee TH, Lee SR, Chang KT, Kang SW, Kim JM, Yu DY, Lee SH, Lee DS (2013) Peroxiredoxin I is a ROS/p38 MAPK-dependent inducible antioxidant that regulates NF-kappaB-mediated iNOS induction and microglial activation. J Neuroimmunol 259: 26–36

49. Kirla RM, Haapasalo HK, Kalimo H, Salminen EK (2003) Low expression of p27 indicates a poor prognosis in patients with high-grade astrocytomas. Cancer 97: 644–648

50. Langmead B, Trapnell C, Pop M, Salzberg SL (2009) Ultrafast and memory-efficient alignment of short DNA sequences to the human genome. Genome Biol 10: R25

51. Lathia JD, Mack SC, Mulkearns-Hubert EE, Valentim CL, Rich JN (2015) Cancer stem cells in glioblastoma. Genes Dev 29: 1203–1217

52. Lin H (2008) Cell biology of stem cells: an enigma of asymmetry and self-renewal. J Cell Biol 180: 257–260

53. Lin JI, Poon CL, Harvey KF (2013) The Hippo size control pathway--ever expanding. Sci Signal 6: pe4

54. Liu Y, Ye F, Yamada K, Tso JL, Zhang Y, Nguyen DH, Dong Q, Soto H, Choe J, Dembo A, Wheeler H, Eskin A, Schmid I, Yong WH, Mischel PS, Cloughesy TF, Kornblum HI, Nelson SF, Liau LM, Tso CL (2011) Autocrine endothelin-3/endothelin receptor B signaling maintains cellular and molecular properties of glioblastoma stem cells. Mol Cancer Res 9: 1668–1685

55. Llorens-Bobadilla E, Zhao S, Baser A, Saiz-Castro G, Zwadlo K, Martin-Villalba A (2015) Single-Cell Transcriptomics Reveals a Population of Dormant Neural Stem Cells that Become Activated upon Brain Injury. Cell Stem Cell 17: 329–340

56. Mohrin M, Chen D (2016) The mitochondrial metabolic checkpoint and aging of hematopoietic stem cells. Curr Opin Hematol 23: 318–324

57. Neftel C, Laffy J, Filbin MG, Hara T, Shore ME, Rahme GJ, Richman AR, Silverbush D, Shaw ML, Hebert CM, Dewitt J, Gritsch S, Perez EM, Gonzalez Castro LN, Lan X, Druck N, Rodman C, Dionne D, Kaplan A, Bertalan MS et al (2019) An Integrative Model of Cellular States, Plasticity, and Genetics for Glioblastoma. Cell 178: 835–849 e821

58. Nowakowski TJ, Bhaduri A, Pollen AA, Alvarado B, Mostajo-Radji MA, Di Lullo E, Haeussler M, Sandoval-Espinosa C, Liu SJ, Velmeshev D, Ounadjela JR, Shuga J, Wang X, Lim DA, West JA, Leyrat AA, Kent WJ, Kriegstein AR (2017) Spatiotemporal gene expression trajectories reveal developmental hierarchies of the human cortex. Science 358: 1318–1323

59. Obernier K, Cebrian-Silla A, Thomson M, Parraguez JI, Anderson R, Guinto C, Rodas Rodriguez J, Garcia-Verdugo JM, Alvarez-Buylla A (2018) Adult Neurogenesis Is Sustained by Symmetric Self-Renewal and Differentiation. Cell Stem Cell 22: 221–234 e228

60. Orr BA, Bai H, Odia Y, Jain D, Anders RA, Eberhart CG (2011) Yes-associated protein 1 is widely expressed in human brain tumors and promotes glioblastoma growth. J Neuropathol Exp Neurol 70: 568–577

61. Patel AP, Tirosh I, Trombetta JJ, Shalek AK, Gillespie SM, Wakimoto H, Cahill DP, Nahed BV, Curry WT, Martuza RL, Louis DN, Rozenblatt-Rosen O, Suva ML, Regev A, Bernstein BE (2014) Single-cell RNA-seq highlights intratumoral heterogeneity in primary glioblastoma. Science 344: 1396–1401

62. Plouffe SW, Meng Z, Lin KC, Lin B, Hong AW, Chun JV, Guan KL (2016) Characterization of Hippo Pathway Components by Gene Inactivation. Mol Cell 64: 993–1008

63. Pollard SM, Conti L, Sun Y, Goffredo D, Smith A (2006) Adherent neural stem (NS) cells from fetal and adult forebrain. Cereb Cortex 16 **Suppl 1**: i112–120

64. Pollard SM, Yoshikawa K, Clarke ID, Danovi D, Stricker S, Russell R, Bayani J, Head R, Lee M, Bernstein M, Squire JA, Smith A, Dirks P (2009) Glioma stem cell lines expanded in adherent culture have tumor-specific phenotypes and are suitable for chemical and genetic screens. Cell Stem Cell 4: 568–580

65. Puram SV, Tirosh I, Parikh AS, Patel AP, Yizhak K, Gillespie S, Rodman C, Luo CL, Mroz EA, Emerick KS, Deschler DG, Varvares MA, Mylvaganam R, Rozenblatt-Rosen O, Rocco JW, Faquin WC, Lin DT, Regev A, Bernstein BE (2017) Single-Cell Transcriptomic Analysis of Primary and Metastatic Tumor Ecosystems in Head and Neck Cancer. Cell 171: 1611–1624 e1624

66. Reya T, Morrison SJ, Clarke MF, Weissman IL (2001) Stem cells, cancer, and cancer stem cells. Nature 414: 105–111

67. Robinson MD, McCarthy DJ, Smyth GK (2010) edgeR: a Bioconductor package for differential expression analysis of digital gene expression data. Bioinformatics 26: 139–140

68. Roche B, Arcangioli B, Martienssen R (2017) Transcriptional reprogramming in cellular quiescence. RNA Biol 14: 843–853

69. Sakamoto M, Hirata H, Ohtsuka T, Bessho Y, Kageyama R (2003) The basic helix-loop-helix genes Hesr1/Hey1 and Hesr2/Hey2 regulate maintenance of neural precursor cells in the brain. J Biol Chem 278: 44808–44815

70. Sakaue-Sawano A, Kurokawa H, Morimura T, Hanyu A, Hama H, Osawa H, Kashiwagi S, Fukami K, Miyata T, Miyoshi H, Imamura T, Ogawa M, Masai H, Miyawaki A (2008) Visualizing spatiotemporal dynamics of multicellular cell-cycle progression. Cell 132: 487–498

71. Sanjana NE, Shalem O, Zhang F (2014) Improved vectors and genome-wide libraries for CRISPR screening. Nat Methods 11: 783–784

72. Santos A, Wernersson R, Jensen LJ (2015) Cyclebase 3.0: a multi-organism database on cell-cycle regulation and phenotypes. Nucleic Acids Res 43: D1140–1144

73. Scialdone A, Natarajan KN, Saraiva LR, Proserpio V, Teichmann SA, Stegle O, Marioni JC, Buettner F (2015) Computational assignment of cell-cycle stage from single-cell transcriptome data. Methods 85: 54–61

74. Scott CE, Wynn SL, Sesay A, Cruz C, Cheung M, Gomez Gaviro MV, Booth S, Gao B, Cheah KS, Lovell-Badge R, Briscoe J (2010) SOX9 induces and maintains neural stem cells. Nat Neurosci 13: 1181–1189

75. Scott RW, Arostegui M, Schweitzer R, Rossi FMV, Underhill TM (2019) Hic1 Defines Quiescent Mesenchymal Progenitor Subpopulations with Distinct Functions and Fates in Skeletal Muscle Regeneration. Cell Stem Cell 25: 797–813 e799

76. Shalem O, Sanjana NE, Hartenian E, Shi X, Scott DA, Mikkelsen TS, Heckl D, Ebert BL, Root DE, Doench JG, Zhang F (2014) Genome-scale CRISPR-Cas9 knockout screening in human cells. Science 343: 84–87

77. Shapiro HM (1981) Flow cytometric estimation of DNA and RNA content in intact cells stained with Hoechst 33342 and pyronin Y. Cytometry 2: 143–150

78. Sherr CJ (1995) D-type cyclins. Trends in biochemical sciences 20: 187–190

79. Sherr CJ, McCormick F (2002) The RB and p53 pathways in cancer. Cancer Cell 2: 103–112

80. Spencer SL, Cappell SD, Tsai FC, Overton KW, Wang CL, Meyer T (2013) The proliferation-quiescence decision is controlled by a bifurcation in CDK2 activity at mitotic exit. Cell 155: 369–383

81. Stupp R, Mason WP, van den Bent MJ, Weller M, Fisher B, Taphoorn MJ, Belanger K, Brandes AA, Marosi C, Bogdahn U, Curschmann J, Janzer RC, Ludwin SK, Gorlia T, Allgeier A, Lacombe D, Cairncross JG, Eisenhauer E, Mirimanoff RO, European Organisation for R et al (2005) Radiotherapy plus concomitant and adjuvant temozolomide for glioblastoma. N Engl J Med 352: 987–996

82. Sun Y, Pollard S, Conti L, Toselli M, Biella G, Parkin G, Willatt L, Falk A, Cattaneo E, Smith A (2008) Long-term tripotent differentiation capacity of human neural stem (NS) cells in adherent culture. Mol Cell Neurosci 38: 245–258

83. Susaki E, Nakayama K, Nakayama KI (2007) Cyclin D2 translocates p27 out of the nucleus and promotes its degradation at the G0-G1 transition. Mol Cell Biol 27: 4626–4640

84. Szklarczyk D, Morris JH, Cook H, Kuhn M, Wyder S, Simonovic M, Santos A, Doncheva NT, Roth A, Bork P, Jensen LJ, von Mering C (2017) The STRING database in 2017: quality-controlled protein-protein association networks, made broadly accessible. Nucleic Acids Res 45: D362–D368

85. TCGA (2008) Comprehensive genomic characterization defines human glioblastoma genes and core pathways. Nature 455: 1061–1068

86. Than-Trong E, Ortica-Gatti S, Mella S, Nepal C, Alunni A, Bally-Cuif L (2018) Neural stem cell quiescence and stemness are molecularly distinct outputs of the Notch3 signalling cascade in the vertebrate adult brain. Development 145

87. Tirosh I, Venteicher AS, Hebert C, Escalante LE, Patel AP, Yizhak K, Fisher JM, Rodman C, Mount C, Filbin MG, Neftel C, Desai N, Nyman J, Izar B, Luo CC, Francis JM, Patel AA, Onozato ML, Riggi N, Livak KJ et al (2016) Single-cell RNA-seq supports a developmental hierarchy in human oligodendroglioma. Nature 539: 309–313

88. Toledo CM, Ding Y, Hoellerbauer P, Davis RJ, Basom R, Girard EJ, Lee E, Corrin P, Hart T, Bolouri H, Davison J, Zhang Q, Hardcastle J, Aronow BJ, Plaisier CL, Baliga NS, Moffat J, Lin Q, Li XN, Nam DH et al (2015) Genome-wide CRISPR-Cas9 Screens Reveal Loss of Redundancy between PKMYT1 and WEE1 in Glioblastoma Stem-like Cells. Cell Rep 13: 2425–2439

89. Toledo CM, Herman JA, Olsen JB, Ding Y, Corrin P, Girard EJ, Olson JM, Emili A, DeLuca JG, Paddison PJ (2014) BuGZ is required for Bub3 stability, Bub1 kinetochore function, and chromosome alignment. Dev Cell 28: 282–294

90. Trapnell C, Roberts A, Goff L, Pertea G, Kim D, Kelley DR, Pimentel H, Salzberg SL, Rinn JL, Pachter L (2012) Differential gene and transcript expression analysis of RNA-seq experiments with TopHat and Cufflinks. Nat Protoc 7: 562–578

91. Venteicher AS, Tirosh I, Hebert C, Yizhak K, Neftel C, Filbin MG, Hovestadt V, Escalante LE, Shaw ML, Rodman C, Gillespie SM, Dionne D, Luo CC, Ravichandran H, Mylvaganam R, Mount C, Onozato ML, Nahed BV, Wakimoto H, Curry WT et al (2017) Decoupling genetics, lineages, and microenvironment in IDH-mutant gliomas by single-cell RNA-seq. Science 355: eaai8478

92. Wang Q, Hu B, Hu X, Kim H, Squatrito M, Scarpace L, deCarvalho AC, Lyu S, Li P, Li Y, Barthel F, Cho HJ, Lin YH, Satani N, Martinez-Ledesma E, Zheng S, Chang E, Sauve CG, Olar A, Lan ZD, et al (2017) Tumor Evolution of Glioma-Intrinsic Gene Expression Subtypes Associates with Immunological Changes in the Microenvironment. Cancer Cell 32: 42–56 e46

93. Weinberg RA (1995) The retinoblastoma protein and cell cycle control. Cell 81: 323–330

94. Wilson KE, Li YW, Yang N, Shen H, Orillion AR, Zhang J (2014) PTPN14 forms a complex with Kibra and LATS1 proteins and negatively regulates the YAP oncogenic function. J Biol Chem 289: 23693–23700

95. Wu HN, Cao XL, Fang Z, Zhang YF, Han WJ, Yue KY, Cao Y, Zheng MH, Wang LL, Han H (2019) Deficiency of Ttyh1 downstream to Notch signaling results in precocious differentiation of neural stem cells. Biochem Biophys Res Commun 514: 842–847

96. Yan H, Parsons DW, Jin G, McLendon R, Rasheed BA, Yuan W, Kos I, Batinic-Haberle I, Jones S, Riggins GJ, Friedman H, Friedman A, Reardon D, Herndon J, Kinzler KW, Velculescu VE, Vogelstein B, Bigner DD (2009) IDH1 and IDH2 mutations in gliomas. N Engl J Med 360: 765–773

97. Yao G, Lee TJ, Mori S, Nevins JR, You L (2008) A bistable Rb-E2F switch underlies the restriction point. Nat Cell Biol 10: 476–482

98. Young MD, Wakefield MJ, Smyth GK, Oshlack A (2010) Gene ontology analysis for RNA-seq: accounting for selection bias. Genome Biol 11: R14

99. Yu G, He QY (2016) ReactomePA: an R/Bioconductor package for reactome pathway analysis and visualization. Mol Biosyst 12: 477–479

100. Zetterberg A, Larsson O, Wiman KG (1995) What is the restriction point? Curr Opin Cell Biol 7: 835–842

101. Zhang H, Geng D, Gao J, Qi Y, Shi Y, Wang Y, Jiang Y, Zhang Y, Fu J, Dong Y, Gao S, Yu R, Zhou X (2016a) Expression and significance of Hippo/YAP signaling in glioma progression. Tumour Biol

102. Zhang L, Laaniste L, Jiang Y, Alafuzoff I, Uhrbom L, Dimberg A (2016b) Pleiotrophin enhances PDGFB-induced gliomagenesis through increased proliferation of neural progenitor cells. Oncotarget 7: 80382–80390

103. Zhang N, Bai H, David KK, Dong J, Zheng Y, Cai J, Giovannini M, Liu P, Anders RA, Pan D (2010) The Merlin/NF2 tumor suppressor functions through the YAP oncoprotein to regulate tissue homeostasis in mammals. Dev Cell 19: 27–38

104. Zong H, Parada LF, Baker SJ (2015) Cell of origin for malignant gliomas and its implication in therapeutic development. Cold Spring Harbor perspectives in biology 7

