## Supplementary material for "Neural G0: a quiescent-like state found in neuroepithelial-derived cells and glioma": Supplmental Data Inventory

**Supplemental Data Inventory**

**SUPPLEMENTARY FIGURES**

**Figure S1: Data in support of scRNA-seq data presented in Figure 1.**

**Figure S2: Protein-protein interaction networks based on the STRING database for the enriched genes that define cell cycle clusters after single-cell RNA-sequencing in unsorted U5-hNSCs.**

**Figure S3: scRNA-seq analysis of embryonic stem cells.**

**Figure S4: Comparison of ccAF and Seurat cell cycle classifiers with scRNA-seq data for HEK293 cells.**

**Figure S5: scRNA-seq analysis human gliomas.**

**Figure S6: Examination of select Neural G0 genes using genes expression data from TCGA pan cancer analysis.**

**Figure S7: Association of IDH1/2 mutations with Neural G0 gene expression and Cell cycle genes.**

**Figure S8: CRISPR-Cas9 gene knockout screens to identify candidate Neural G0 regulating genes in hNSCs.**

**Figure S9: Additional screens to identify Neural G0-skip mutants.**

**Figure S10: Further retests and validation of G0-skip genes.**

**Figure S11: Data supporting Figure 6D for time lapse analysis of NSC cell cycle using FUCCI factors.**

**Figure S12: Loss of G0-skip Genes Reduces G0 and Increases Late G1 Molecular Features.**

**Figure S13: Transcriptional Target and Cell Cycle Gene Expression Following Loss of G0-skip Genes.**

**Figure S14: Gene Ontology Analysis of Genes Down Regulated in G0/G1 Following Loss of G0-skip Genes in hNSCs.**

**Figure S15: Gene Ontology Analysis of Genes Up Regulated in G0/G1 Following Loss of G0-skip Genes in hNSCs.**

**Figure S16: hNSC cell cycle cluster genes altered in patient-derived GSCs.**

**Figure S17: Additional gene expression analysis of G0-skip mutants in hNSCs.**

**Figure S18: scRNA-seq of TAOK1 KO hNSCs.**

**SUPPLEMENTARY TABLES**

**Table S1: Enriched and Depleted Genes in Cell Cycle Clusters Identified by Single-cell RNA-seq in U5-hNSCs.**

**Table S2: Gene Ontology Analysis of Defining Genes in Transcriptional Cell Cycle Clusters in U5-NSCs.**

**Table S3: scRNA-seq analysis of developing human telencephalon and human gliomas (in support of Fig. 2E and Table 1).**

**Table S4: Top differentially expressed Neural G0 genes from human glioma scRNA-seq data sets.**

**Table S5: CRISPR-Cas9 Screens to Identify and Validate Candidate Neural G0-Skip Genes in hNSCs.**

**Table S6: Gene Expression of G0/G1-sorted Cells Compared to Unsorted U5-NSCs, after G0-skip KO, and in GSC Isolates.**

**Table S7: Biological Process Gene Ontology Analysis of G0/G1 Transcriptional Reprogramming in U5-NSCs Following Loss of G0-skip Genes.**

**Table S8: List of Key Resources Including Antibodies, Validation sgDNA Sequences, and Primers**

**PRIMARY SCREEN AND GENE EXPRESSION DATA**

**NCBI Gene Expression Omnibus GSE117004** (Token = wvijeqcivtcbdop**)**
