## Supplemental Figs and Legends for "Neural G0: a quiescent-like state found in neuroepithelial-derived cells and glioma"

#### Supplemental Figures and Legends

**Figure S1: Data in support of scRNA-seq data presented in Figure 1.**
**A,** Comparison of cluster size from scRNA-seq for the entire cell population of U5-NSCs com- pared to a subpopulation sorted for G0/G1 (mCherry-CDT1(aa30-120)-positive). The clusters with an increase in representation were characterized as G0/G1. Clusters that remained the same or moderately decreased in representation are considered transition states with some of the cells retaining or initiating CDT1 accumulation. Clusters that decreased in representation greater than 2-fold were considered S/G2/M and mCherry-CDT1-negative.

**B**, Overlap of conserved YAP target genes up-regulated following G0-skip knockout of *NF2* (yellow), *TAOK1* (red), or both (purple) with genes defining cell cycle transcriptional clusters. The NF2-dependent YAP targets significantly overlap with the Late G1 cluster, while the TAOK1-dependent genes overlap with the S/G2 and G2/M clusters. Both *NF2* and *TAOK1* knockouts are analyzed as G0-skip genes further in Figs. 6 and S10. Significance assessed using hypergeometric analysis.

**C**, Dot plot for some of the key markers for each cluster progressing through cell cycle. Color is according to average expression (Avg. Exp.) per cell, while size of the dots corresponds to the proportion of cells within the cluster in which the gene was sequenced (% Exp.). Red text indicates cyclin and cyclin-dependent kinase genes that are differentially expressed in the different clusters. The genes that are depleted in each cluster supports the G0/G1 classification of the cells. For example, CDK1 is significantly reduced in Neural G0, G1, Late G1, and early G1/M, but enriched in S, S/G2, and G2/M. Similar depletion of other cell cycle genes is seen, including MKI67 (depleted in Neural G0, late G1, and enriched in G2/M and S/G2, AURKA (depleted in Neural G0, G1, S, and M/early G1 and enriched in G2/M), CCNA2 (depleted in Neural G0, G1, late G1, and M/early G1 and enriched in S/G2 and G2/M), and TOP2A (depleted in Neural G0, G1, late G1, and M/early G1 and enriched in S/G2 and G2/M). In addition, Neural G0 showed the most significant repression of these genes peaking in other phases of cell cycle, including suppression of CCND1 expression.

**D**, Number of unique molecular identifiers (nUMI) per cell in each single cell RNA-seq cluster. The Low RNA cluster has dramatically less nUMI than any other cluster, making it difficult to distinguish actual cells belonging to this cluster from background. Therefore, this cluster was excluded from further analysis. The nUMI increases as the cell cycle progresses, with G2/M cells having the greatest nUMI. Significance assessed using a two-tailed student's t-test.

### Figure S1

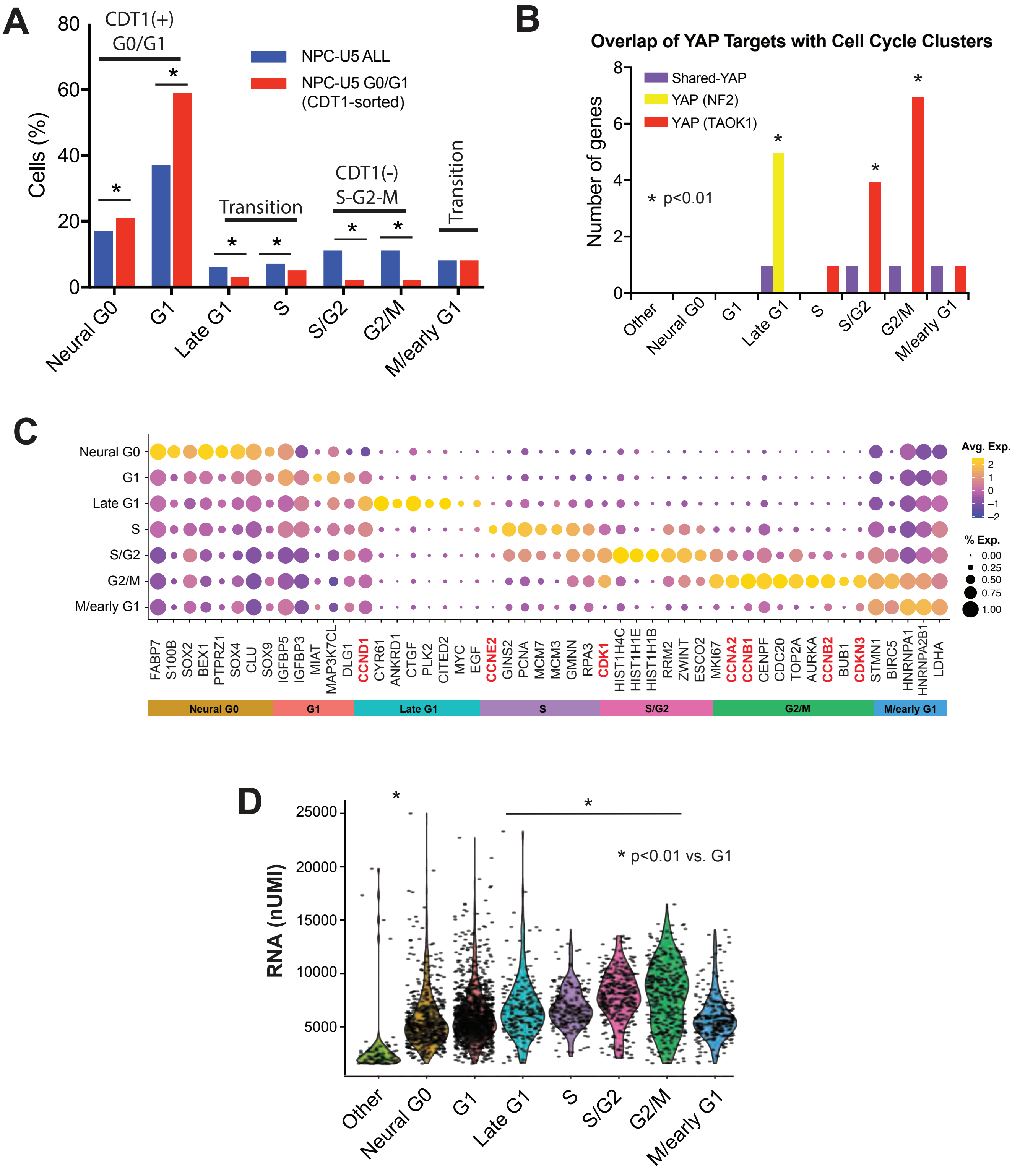
A
CDT1(+) G0/G1
NPC-U5 ALL
NPC-U5 G0/G1 (CDT1-sorted)
Cells (%)
Transition
CDT1(-) S-G2-M
Transition
Neural G0
G1
Late G1
S
S/G2
G2/M
M/early G1
B
Overlap of YAP Targets with Cell Cycle Clusters
Shared-YAP
YAP (NF2)
YAP (TAOK1)
Number of genes
* p<0.01
Other
Neural G0
G1
Late G1
S
S/G2
G2/M
M/early G1
C
Avg. Exp.
% Exp.
FABP7
S100B
SOX2
BEX1
PTPRZ1
SOX4
CLU
SOX9
IGFBP5
IGFBP3
MIAT
MAP3K7CL
DLG1
CCND1
CYR61
ANKRD1
CTGF
PLK2
CITED2
MYC
EGF
CCNE2
GINS2
PCNA
MCM7
MCM3
GMNN
RPA3
CDK1
HIST1H4C
HIST1H1E
HIST1H1B
RRM2
ZWINT
ESCO2
MKI67
CCNA2
CCNB1
CENPF
CDC20
TOP2A
AURKA
CCNB2
BUB1
CDKN3
STMN1
BIRC5
HNRNPA1
HNRNPA2B1
LDHA
D
RNA (nUMI)
* p<0.01 vs. G1
Other
Neural G0
G1
Late G1
S
S/G2
G2/M
M/early G1

**Figure S2: Protein-protein interaction networks based on the STRING database for the enriched genes that define cell cycle clusters after single-cell RNA-sequencing in unsorted U5-hNSCs.** The thickness of the connecting lines corresponds to the strength of the predicted interaction based on experimental data, curated databases, co-expression analysis, and gene ontology. The network visualization only includes genes that connect with the rest of the network.

**A**, enriched genes that define the G0/G1 clusters: Neural G0, Early G1/M and Late G1. The network visualization only includes genes that connect with the rest of the network. G1 only showed enriched interactions between IGFR1 signaling genes IGFBP3 and IGFBP5 (not visualized). Neural G0 has a more diffuse network with only a few interactions per gene and includes a number of transcription factors that may be important in mediating this cell cycle state (pink). The late G1 cluster has two gene groups: the cholesterol synthesis group (orange) and YAP-NF2 targets (yellow). M/early G1 retains residual expression or non-degradation of a subset of genes enriched in G2/M, including the mitosis-peaking genes CCNB2, CDKN3, and BIRC5 (red) as well as spliceosomal genes (green).

**B**, G2/M, S, and S/G2 cluster Genes from enriched gene ontology groups are indicated by colored nodes, including mitotic cell division (red), DNA replication (purple), DNA conformation change (yellow), and RNA-related genes from various gene ontology biological processes (green).

All genes from each scRNA-seq cluster are available in Table S1.

### Figure S2

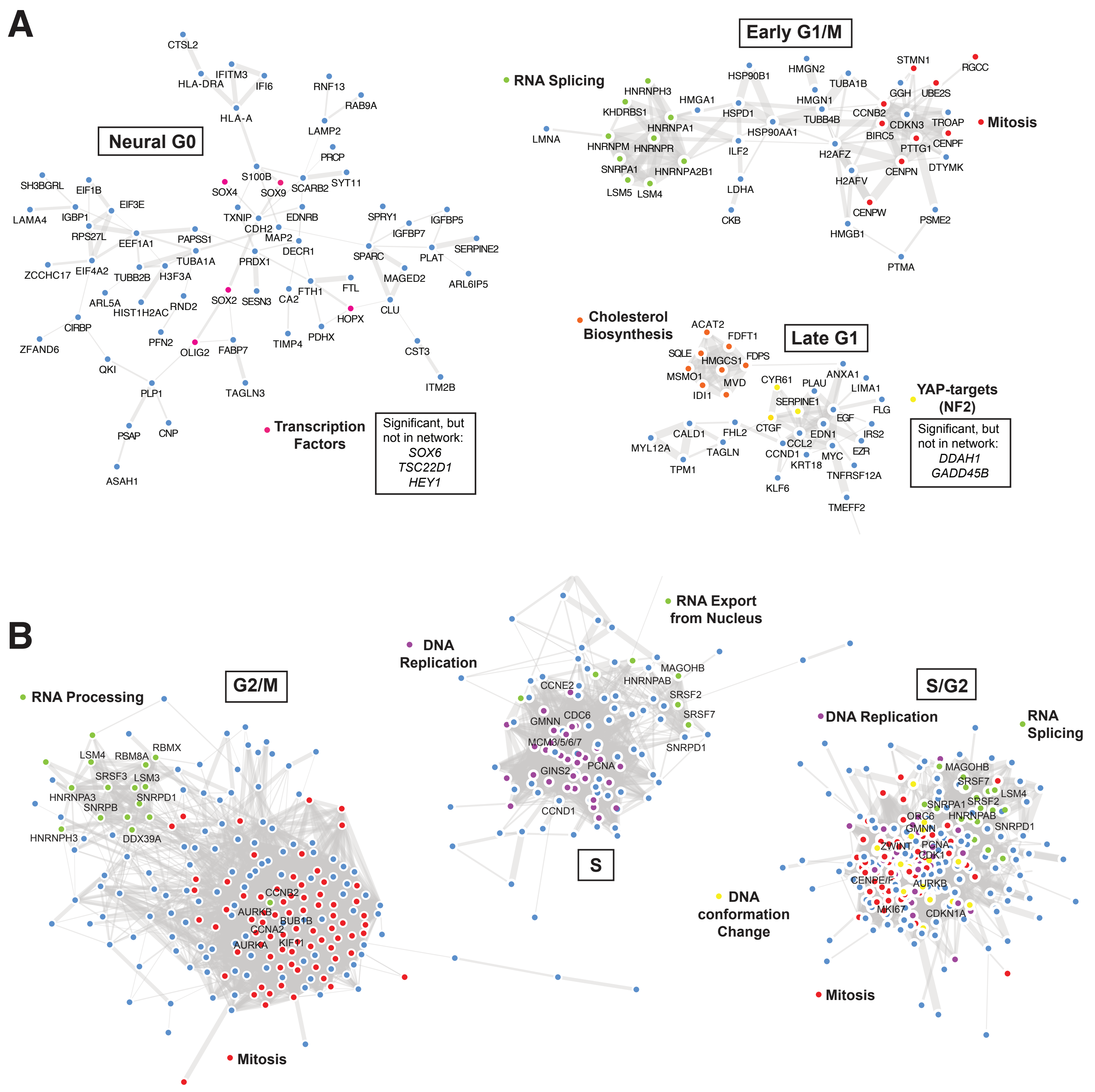
A
Neural G0
CTSL2
IFITM3
HLA-DRA
IFI6
RNF13
RAB9A
HLA-A
LAMP2
PRCP
S100B
SOX4
SOX9
SCARB2
SYT11
SH3BGRL
EIF1B
EIF3E
LAMA4
IGBP1
TXNIP
EDNRB
CDH2
SPRY1
IGFBP5
RPS27L
EEF1A1
PAPSS1
MAP2
IGFBP7
DECR1
SERPINE2
TUBA1A
PRDX1
SPARC
PLAT
ZCCHC17
EIF4A2
TUBB2B
H3F3A
MAGED2
ARL6IP5
FTH1
FTL
ARL5A
HIST1H2AC
RND2
SOX2
SESN3
CA2
CIRBP
HOPX
CLU
ZFAND6
PFN2
OLIG2
FABP7
TIMP4
PDHX
CST3
QKI
PLP1
TAGLN3
ITM2B
PSAP
CNP
ASAH1
Transcription Factors
Significant, but not in network:
SOX6
TSC22D1
HEY1
Early G1/M
RNA Splicing
HNRNPH3
HMGA1
KHDRBS1
HNRNPA1
HNRNPM
HNRNPR
SNRPA1
HNRNPA2B1
LSM5
LSM4
LMNA
HSP90B1
HSPD1
HSP90AA1
ILF2
LDHA
CKB
HMGN2
TUBA1B
HMGN1
TUBB4B
H2AFZ
H2AFV
HMGB1
CENPW
STMN1
RGCC
GGH
UBE2S
CCNB2
CDKN3
TROAP
BIRC5
CENPF
PTTG1
CENPN
DTYMK
PSME2
PTMA
Mitosis
Cholesterol Biosynthesis
ACAT2
FDFT1
SQLE
HMGCS1
FDPS
MSMO1
MVD
IDI1
Late G1
CYR61
PLAU
ANXA1
LIMA1
SERPINE1
FLG
EGF
CTGF
EDN1
IRS2
CALD1
FHL2
CCL2
MYL12A
TAGLN
CCND1
EZR
MYC
TPM1
KRT18
TNFRSF12A
KLF6
TMEFF2
YAP-targets (NF2)
Significant, but not in network:
DDAH1
GADD45B
B
RNA Processing
G2/M
LSM4
RBM8A
RBMX
SRSF3
LSM3
HNRNPA3
SNRPB
SNRPD1
HNRNPH3
DDX39A
CCNB2
AURKB
BUB1B
CCNA2
AURKA
KIF11
Mitosis
DNA Replication
RNA Export from Nucleus
S
CCNE2
MAGOHB
HNRNPAB
SRSF2
SRSF7
GMNN
CDC6
MCM3/5/6/7
SNRPD1
GINS2
PCNA
CCND1
S/G2
DNA Replication
RNA Splicing
MAGOHB
SRSF7
LSM4
SRSF2
SNRPA1
ORC6
HNRNPAB
SNRPD1
GMNN
ZWINT
PCNA
CDK1
CENPE/F
AURKB
MKI67
CDKN1A
DNA conformation Change
Mitosis

**Figure S3: scRNA-seq analysis of embryonic stem cells and human gliomas.** Proportion of mouse embryonic stem cells (mESCs) (Buettner et al., 2015) classified to each hNSC transcriptional cluster following sorting for cell cycle by Hoechst staining for amount of DNA.

### Figure S3

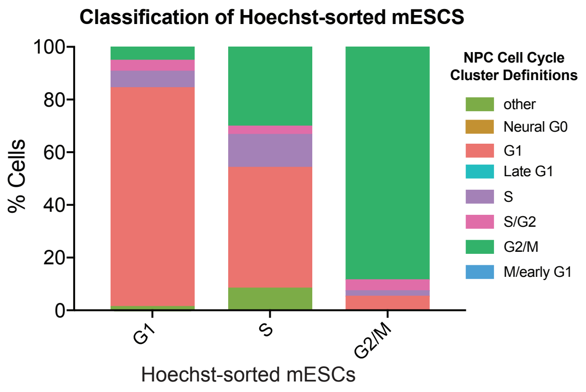

Classification of Hoechst-sorted mESCS
% Cells
100
80
60
40
20
0
G1
S
G2/M
Hoechst-sorted mESCs
NPC Cell Cycle Cluster Definitions
other
Neural G0
G1
Late G1
S
S/G2
G2/M
M/early G1

**Figure S4: Comparison of ccAF and Seurat cell cycle classifiers with scRNA-seq data for HEK293 cells.**
**A,** UMAP plots of ccAF and Seurat cell cycle classifiers of scRNA-seq HEK293 data.
**B,** A graph showing the composition of Seurat cell calls within ccAF called cell populations.
**C,** Ridge graph comparisons of cyclin and CDK1 expression in ccAF- or Seurat-classified cell cycle phases.

### Figure S4

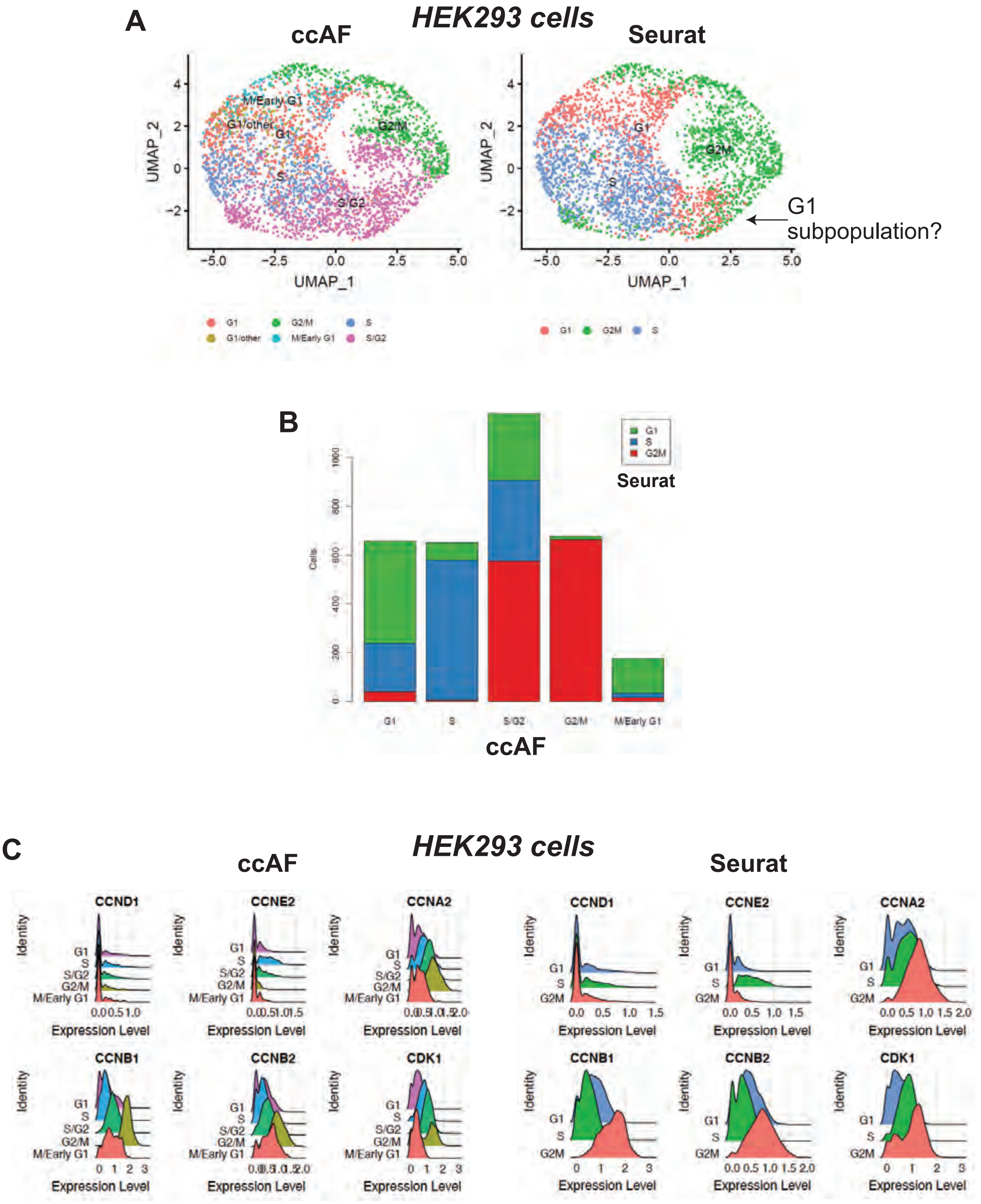
A
HEK293 cells
ccAF
Seurat
M/Early G1
G1/other
G1
G2/M
S
S/G2
G2M
UMAP_2
UMAP_1
G1
subpopulation?
G1
G2/M
S
G1/other
M/Early G1
S/G2
G1
G2M
S
B
G1
S
G2M
Seurat
Cells
G1
S
S/G2
G2/M
M/Early G1
ccAF
C
HEK293 cells
ccAF
Seurat
CCND1
CCNE2
CCNA2
CCNB1
CCNB2
CDK1
Identity
Expression Level
G1
S
S/G2
G2/M
M/Early G1
G2M

**Figure S5: scRNA-seq analysis human gliomas.** Gene expression of top 12 genes associated with Neural G0 in scRNA-seq of glioma tumor from GSE84465 for Table 1. G1 cells and tumor-associated CD45+ immune cells are also shown. A full list of Neural G0 glioma-associated genes is available in Table S3.

### Figure S5

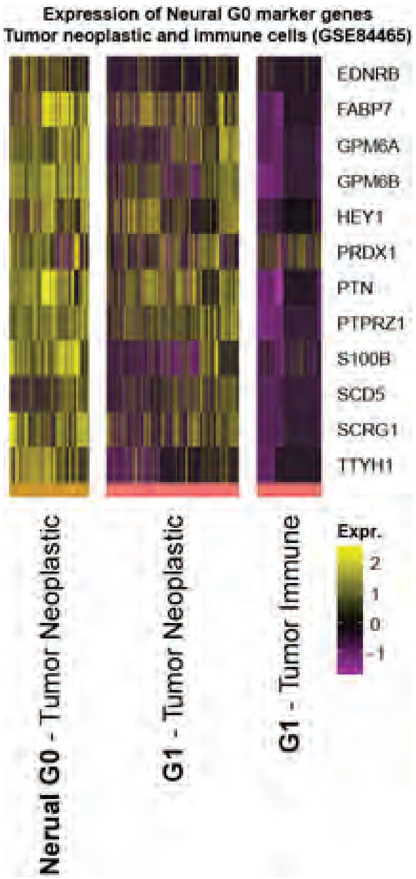
Expression of Neural G0 marker genes
Tumor neoplastic and immune cells (GSE84465)
EDNRB
FABP7
GPM6A
GPM6B
HEY1
PRDX1
PTN
PTPRZ1
S100B
SCD5
SCRG1
TTYH1
Nerual G0 - Tumor Neoplastic
G1 - Tumor Neoplastic
G1 - Tumor Immune
Expr.
2
1
0
-1

**Figure S6: Examination of select Neural G0 genes using genes expression data from TCGA pan cancer analysis.** All of the Neural G0 genes shown in Fig S3B were examined using cBioportal TCGA analysis pipeline (Gao et al. 2013). Each showed results similar to *GPM6B*, *S100B*, and *TTYH1*, which are displayed in the figure.

### Figure S6

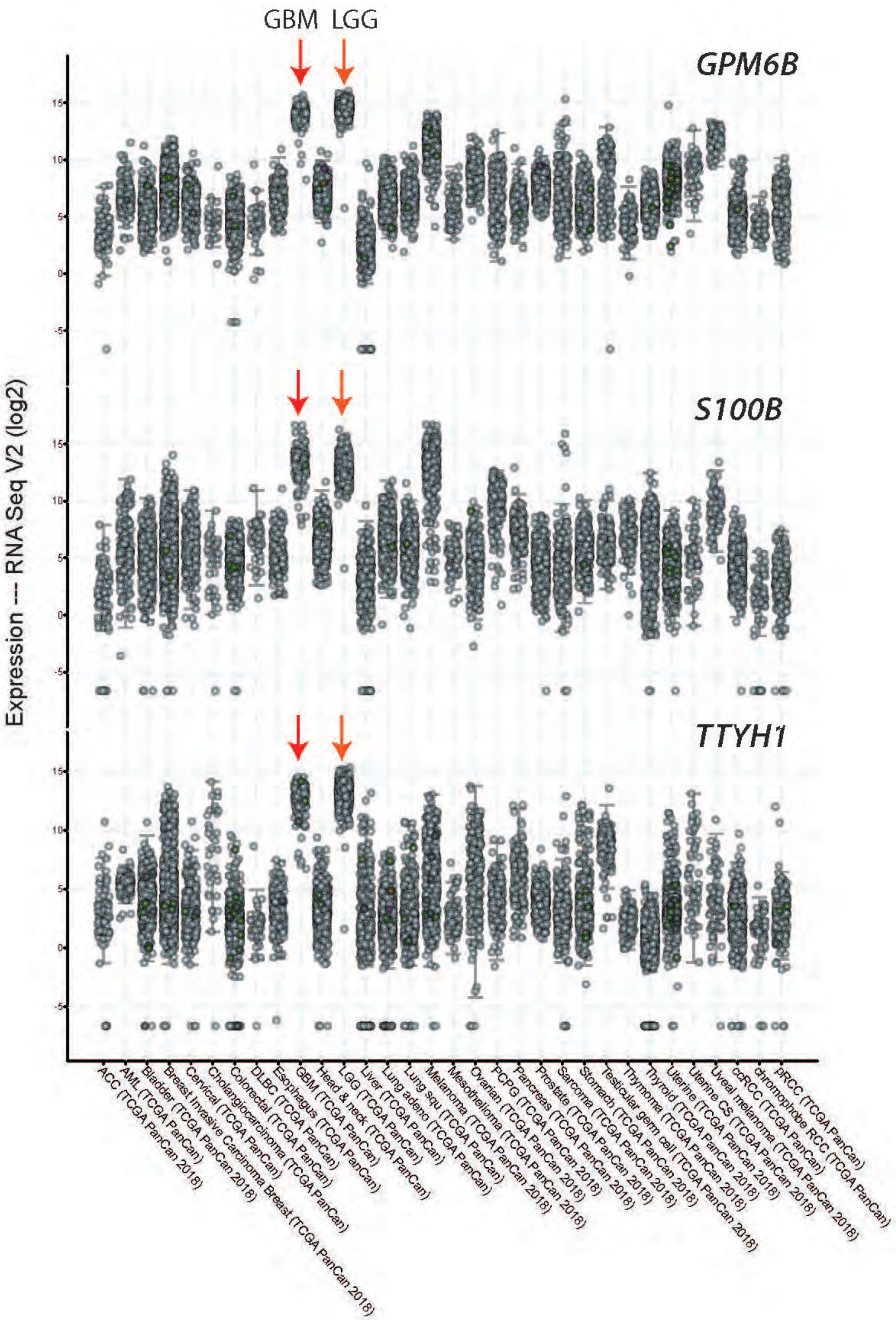
GBM
LGG
GPM6B
S100B
TTYH1
Expression --- RNA Seq V2 (log2)
15
10
5
0
-5
ACC (TCGA PanCan 2018)
AML (TCGA PanCan)
Bladder (TCGA PanCan 2018)
Breast Invasive Carcinoma Breast (TCGA PanCan 2018)
Cervical (TCGA PanCan)
Cholangiocarcinoma (TCGA PanCan)
Colorectal (TCGA PanCan)
DLBC (TCGA PanCan)
Esophagus (TCGA PanCan)
GBM (TCGA PanCan)
Head & neck (TCGA PanCan)
LGG (TCGA PanCan)
Liver (TCGA PanCan)
Lung adeno (TCGA PanCan)
Lung squ (TCGA PanCan)
Melanoma (TCGA PanCan 2018)
Mesothelioma (TCGA PanCan 2018)
Ovarian (TCGA PanCan 2018)
PCPG (TCGA PanCan 2018)
Pancreas (TCGA PanCan 2018)
Prostate (TCGA PanCan 2018)
Sarcoma (TCGA PanCan 2018)
Stomach (TCGA PanCan 2018)
Testicular germ cell (TCGA PanCan 2018)
Thymoma (TCGA PanCan 2018)
Thyroid (TCGA PanCan 2018)
Uterine (TCGA PanCan 2018)
Uterine CS (TCGA PanCan 2018)
Uveal melanoma (TCGA PanCan 2018)
ccRCC (TCGA PanCan)
chromophobe RCC (TCGA PanCan)
pRCC (TCGA PanCan)

**Figure S7: Association of IDH1/2 mutations with Neural G0 gene expression and Cell cycle genes.** An eigengene represents the common variation across each patient tumor for the Neural G0 genes or Cell Cycle genes, i.e., the first principal component corrected for direction if necessary. All pairwise Student's T-tests comparisons had p-values <0.001.

### Figure S7

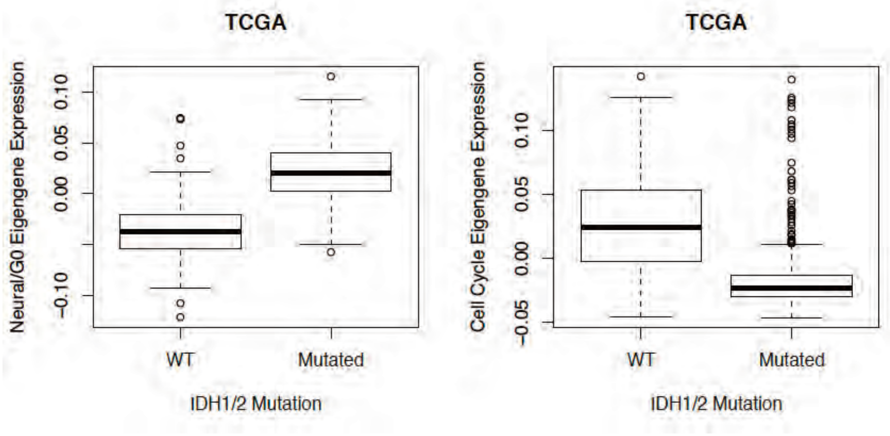
TCGA
Neural/G0 Eigengene Expression
0.10
0.05
0.00
−0.10
WT
Mutated
IDH1/2 Mutation
TCGA
Cell Cycle Eigengene Expression
0.10
0.05
0.00
−0.05
WT
Mutated
IDH1/2 Mutation

**Figure S8: CRISPR-Cas9 gene knockout screens to identify candidate Neural G0 regulating genes in hNSCs.**
**A,** Schematic of CRISPR-Cas9 knockout screen design.

**B,** Enrichment of guide DNA following 21 days outgrowth post-selection using GECKO library (n=2). Colored guides are enriched (z-score>2; green) or depleted (z-score<2; red) with false discovery rate (FDR<0.05). Statistical source data provided in Supplementary Table S6.

**C,** Enrichment of sgRNA using GECKO (n=2), Brunello(Doench et al. 2016) (n=3), or a custom-designed retest library (n=4), with U5-NSC or CB660-NSCs at 10 days or 3 weeks post-selection. Colored guides are enriched (z-score>2; green) or depleted (z-score<2; red) with false discovery rate (FDR)<0.05. Statistical source data provided in Supplementary Table S5.

### Figure S8

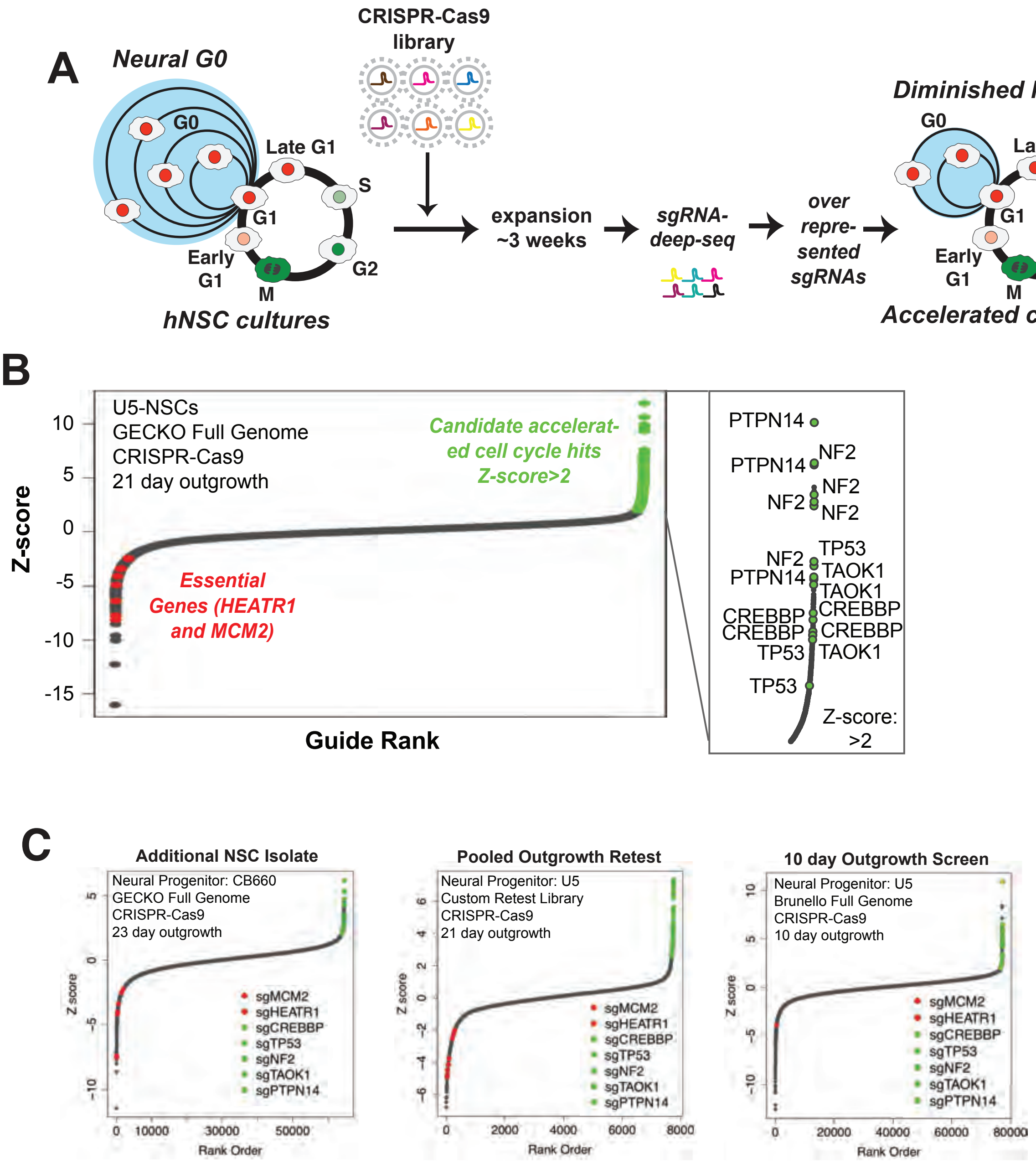
A
Neural G0
CRISPR-Cas9
library
G0
Late G1
S
G1
Early
G1
M
G2
hNSC cultures
expansion
~3 weeks
sgRNA-
deep-seq
over
repre-
sented
sgRNAs
Diminished Neural G0?
Accelerated cell cycle?
B
U5-NSCs
GECKO Full Genome
CRISPR-Cas9
21 day outgrowth
Candidate accelerat-
ed cell cycle hits
Z-score>2
Essential
Genes (HEATR1
and MCM2)
Z-score
Guide Rank
PTPN14
NF2
TP53
TAOK1
CREBBP
Z-score:
>2
C
Additional NSC Isolate
Neural Progenitor: CB660
GECKO Full Genome
CRISPR-Cas9
23 day outgrowth
Pooled Outgrowth Retest
Neural Progenitor: U5
Custom Retest Library
CRISPR-Cas9
21 day outgrowth
10 day Outgrowth Screen
Neural Progenitor: U5
Brunello Full Genome
CRISPR-Cas9
10 day outgrowth
sgMCM2
sgHEATR1
sgCREBBP
sgTP53
sgNF2
sgTAOK1
sgPTPN14
Z score
Rank Order

**Figure S9: Additional Screens to identify Neural G0-skip mutants.**
**A**, Overlap of screen hits between the four CRISPR-Cas9 screens. For all screens, hits were defined as genes having multiple guides with FDR<.01 and z-score>2. For Brunello library, there were additional requirements of at least one guide with z-score>3 and at least 20 reads at Day 0. *CREBBP*, *NF2*, *PTPN14*, *TAOK1*, and *TP53* were defined as hits in every screen.

**B,** Overlap of screen hits between the four CRISPR-Cas9 screens. For all screens, hits were defined as genes having multiple guides with FDR<.01 and z-score >2. For Brunello library, there were additional requirements of at least one guide with z-score >3 and at least 20 reads at Day 0. *CREBBP*, *NF2*, *PTPN14*, *TAOK1*, and *TP53* were defined as hits in every screen.

**C**, Significant overlap between candidate growth-limiting genes and top decile pan-cancer mutated genes (The Cancer Genome Atlas (TCGA); hypergeometric analysis).

**D**, Overlap of the candidate antiproliferative genes with validated tumor suppressors (Futreal et al., 2004) (Tier 1 and 2; Cancer Gene Census v84). Significance assessed using hypergeometric analysis.

### Figure S9

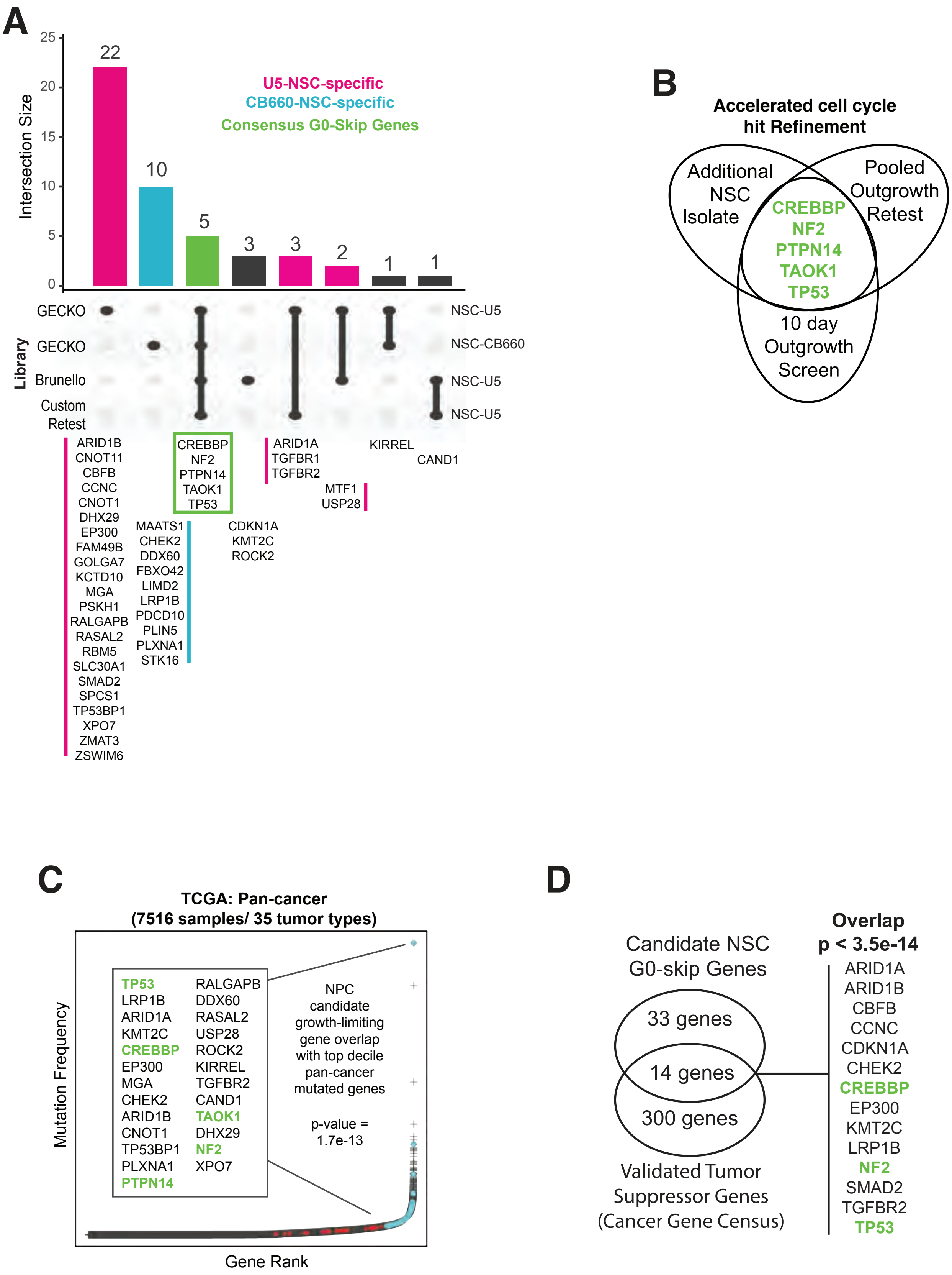
A
Intersection Size
25
20
15
10
5
0
22
10
5
3
3
2
1
1
U5-NSC-specific
CB660-NSC-specific
Consensus G0-Skip Genes
Library
GECKO
GECKO
Brunello
Custom Retest
NSC-U5
NSC-CB660
NSC-U5
NSC-U5
ARID1B
CNOT11
CBFB
CCNC
CNOT1
DHX29
EP300
FAM49B
GOLGA7
KCTD10
MGA
PSKH1
RALGAPB
RASAL2
RBM5
SLC30A1
SMAD2
SPCS1
TP53BP1
XPO7
ZMAT3
ZSWIM6
MAATS1
CHEK2
DDX60
FBXO42
LIMD2
LRP1B
PDCD10
PLIN5
PLXNA1
STK16
CREBBP
NF2
PTPN14
TAOK1
TP53
CDKN1A
KMT2C
ROCK2
ARID1A
TGFBR1
TGFBR2
MTF1
USP28
KIRREL
CAND1
B
Accelerated cell cycle hit Refinement
Additional NSC Isolate
Pooled Outgrowth Retest
10 day Outgrowth Screen
CREBBP
NF2
PTPN14
TAOK1
TP53
C
TCGA: Pan-cancer
(7516 samples/ 35 tumor types)
Mutation Frequency
Gene Rank
TP53
LRP1B
ARID1A
KMT2C
CREBBP
EP300
MGA
CHEK2
ARID1B
CNOT1
TP53BP1
PLXNA1
PTPN14
RALGAPB
DDX60
RASAL2
USP28
ROCK2
KIRREL
TGFBR2
CAND1
TAOK1
DHX29
NF2
XPO7
NPC candidate growth-limiting gene overlap with top decile pan-cancer mutated genes
p-value = 1.7e-13
D
Candidate NSC G0-skip Genes
33 genes
14 genes
300 genes
Validated Tumor Suppressor Genes (Cancer Gene Census)
Overlap
p < 3.5e-14
ARID1A
ARID1B
CBFB
CCNC
CDKN1A
CHEK2
CREBBP
EP300
KMT2C
LRP1B
NF2
SMAD2
TGFBR2
TP53

**Figure S10: Further retests and validation of G0-skip genes.**
**A**, Flow analysis of U5-NSC:GFP with LV-sgRNA:Cas9 retest pools competing with wild-type (WT) U5-NSC over a 23 day outgrowth with ~10% initial proportion (n=3). Competition index refers to the relative increase in %GFP+ compared with initial proportion and mean sgNTC.

**B,** Relative confluency of pooled sgCREBBP and sgNF2, and NTC in U5-NSCs compared to initial confluency over 72 hours at subconfluent density using time lapse microscopy (n=4 sampling regions x 3 biological replicates).

**C,** Representative images of cell confluency at time points 0 and 72h in LV-sgCREBBP and sgNF2 treated cells. Scale bars = 150 μm and 50 μm.

**D,** Confluency of cells relative to time=0 in comparison to NTC after 48 hours using time-lapse microscopy (n=4 sampling regions x 3 biological replicates).

**E**, Doubling time measurements (>14 days post-selection) in U5-NSCs or GSCs after 3-5 days outgrowth. n≥3, as noted in bars for each guide. The data are presented as the mean ± standard deviation (SD). Significance was assessed using a two-tailed student's t-test.

**F**, Western blot confirmation of CREBBP and NF2 protein depletion after gene targeting using lentiviral (LV) pools (4 guides/gene) sgRNA:Cas9 after >21 days outgrowth.

### Figure S10

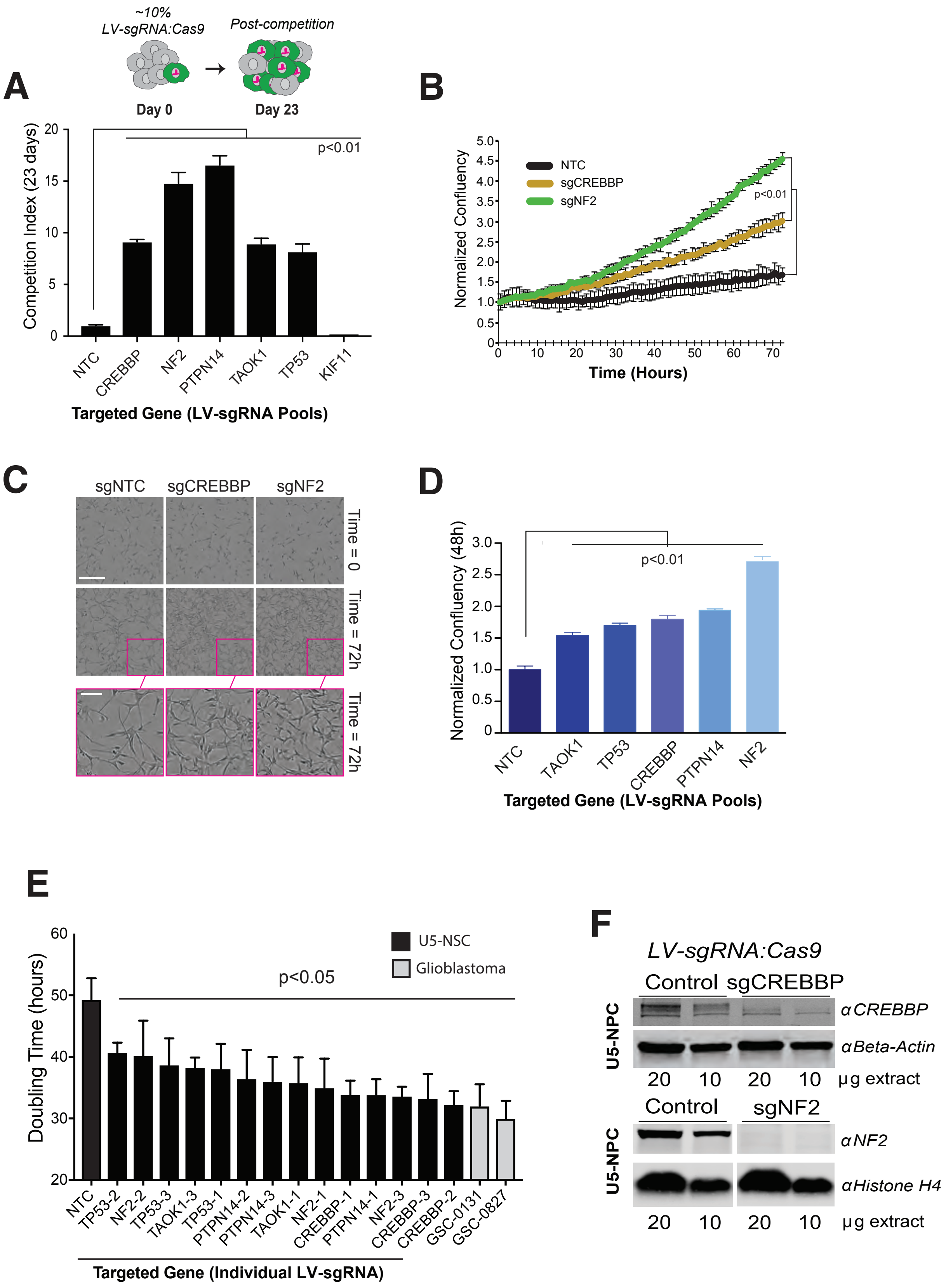
A
~10%
LV-sgRNA:Cas9
Post-competition
Day 0
Day 23
Competition Index (23 days)
p<0.01
NTC
CREBBP
NF2
PTPN14
TAOK1
TP53
KIF11
Targeted Gene (LV-sgRNA Pools)
B
Normalized Confluency
NTC
sgCREBBP
sgNF2
p<0.01
Time (Hours)
C
sgNTC
sgCREBBP
sgNF2
Time = 0
Time = 72h
Time = 72h
D
Normalized Confluency (48h)
p<0.01
NTC
TAOK1
TP53
CREBBP
PTPN14
NF2
Targeted Gene (LV-sgRNA Pools)
E
Doubling Time (hours)
U5-NSC
Glioblastoma
p<0.05
NTC
TP53-2
NF2-2
TP53-3
TAOK1-3
TP53-1
PTPN14-2
PTPN14-3
TAOK1-1
NF2-1
CREBBP-1
PTPN14-1
NF2-3
CREBBP-3
CREBBP-2
GSC-0131
GSC-0827
Targeted Gene (Individual LV-sgRNA)
F
LV-sgRNA:Cas9
Control
sgCREBBP
U5-NPC
αCREBBP
αBeta-Actin
20
10
20
10
μg extract
Control
sgNF2
U5-NPC
αNF2
αHistone H4
20
10
20
10
μg extract

**Figure S11: Data supporting Figure 6D for time lapse analysis of NSC cell cycle using Fucci factors.**

**A-D**, The complete cell cycle of individual U5-NSCs expressing Fucci factors that combine cell cycle-dependent degrons with fluorophores. mCherry-CDT1(aa30-120) (red) is present in G0/G1 and mAG-geminin(aa1-110) (green) is present in S/G2/M. The double negative cells correspond to either a transition state between M and G1 or G1 and S phase or cells that have silenced one or both reporters. Double positive cells which express both CDT1 and Geminin are an alternative G1-S transition state but are rare in the U5-NSC isolate. We tracked individual cells from mitosis to mitosis using timelapse microscopy, every 10-15 minutes, and representative cells from long-cycling NTC (A), short-cycling NTC (B), sgCREBBP (4 guide lentiviral pool) (C), and sgTAOK1 (4 guide lentiviral pool) (D) are presented here.

### Figure S11

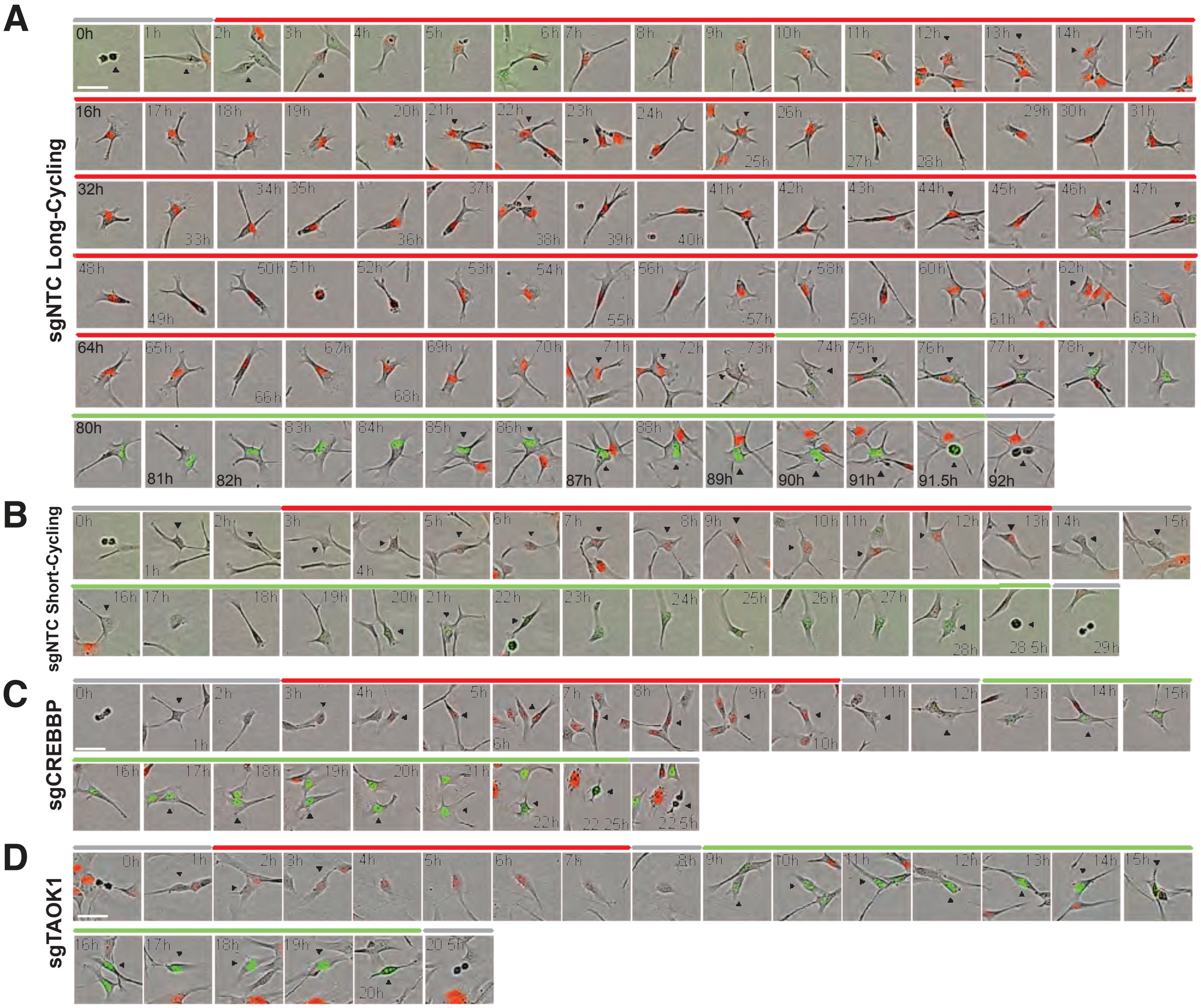
A
sgNTC Long-Cycling
B
sgNTC Short-Cycling
C
sgCREBBP
D
sgTAOK1

**Figure S12: Loss of G0-skip Genes Reduces G0 and Increases Late G1 Molecular Features.**
**A,** Diagram of key molecular features for G0/G1 subpopulations.

**B,** Intensity of immunofluorescence staining for phosphorylated Rb (Ser807/811) in G0/G1 U5-NSCs (mCherry-CDT1+) (Median ± SD; n=2456-8048 cells).

**C,** CDK2 activity in G0/G1 hNSCs after KO measured through the relative ratio of a DNA helicase B (DHB) reporter (Hahn et al. 2009) using fluorescence microscopy. G0/G1 nuclei were identified by the presence of mCherry-CDT1 and due to the irregular shape of the NSCs, the cytoplasmic intensity of the DHB reporter was defined as the upper quartile intensity of a 2-pixel ring around the CDT1-defined nucleus (n=960-8048 cells). *NF2* KO was not significantly different from NTC (p=0.51).

**D,** Proportion of G0/G1 cells with cytoplasmic:nuclear ratio >1 for CDK2 activity reporter (n=54 fields).

**E,** Representative distribution of the entire population of U5-NSCs for $p27K^{-}$-mVenus assayed using flow cytometry after KO along with WT and NTC. Threshold to define $p27^{high}$ set at the top ~10% of the NTC for each independent replicate.

**F,** Proportion of cells $p27^{high}$ following KO, assayed using flow cytometry (n=3).

**G,** Model for the changes in G0/G1 molecular features following loss of *CREBBP*, *NF2*, *PTPN14*, *TAOK1*, or *TP53*.

The data are presented as the mean ± SD. Significance was assessed using a two-tailed student's t test.

### Figure S12

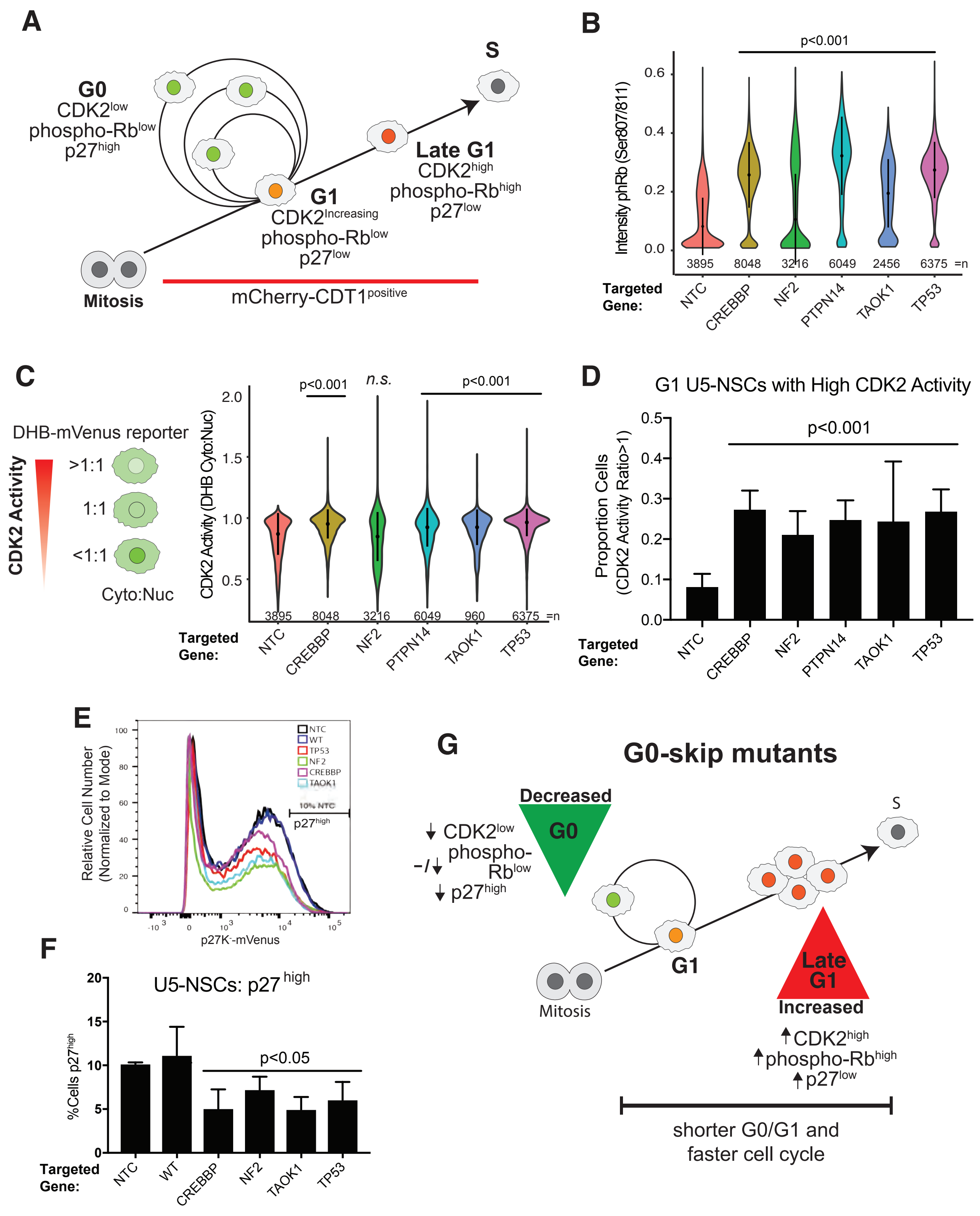

A
G0
CDK2low
phospho-Rblow
p27high
S
Late G1
CDK2high
phospho-Rbhigh
p27low
G1
CDK2Increasing
phospho-Rblow
p27low
Mitosis
mCherry-CDT1positive
B
p<0.001
Intensity phRb (Ser807/811)
3895 8048 3216 6049 2456 6375 =n
Targeted Gene:
NTC CREBBP NF2 PTPN14 TAOK1 TP53
C
DHB-mVenus reporter
CDK2 Activity
>1:1
1:1
<1:1
Cyto:Nuc
CDK2 Activity (DHB Cyto:Nuc)
p<0.001
n.s.
p<0.001
3895 8048 3216 6049 960 6375 =n
Targeted Gene:
NTC CREBBP NF2 PTPN14 TAOK1 TP53
D
G1 U5-NSCs with High CDK2 Activity
p<0.001
Proportion Cells (CDK2 Activity Ratio>1)
Targeted Gene:
NTC CREBBP NF2 PTPN14 TAOK1 TP53
E
Relative Cell Number (Normalized to Mode)
NTC
WT
TP53
NF2
CREBBP
TAOK1
10% NTC
p27high
p27K-mVenus
F
U5-NSCs: p27 high
%Cells p27high
p<0.05
Targeted Gene:
NTC WT CREBBP NF2 TAOK1 TP53
G
G0-skip mutants
Decreased
G0
CDK2low
phospho-Rblow
p27high
S
G1
Mitosis
Late G1
Increased
CDK2high
phospho-Rbhigh
p27low
shorter G0/G1 and faster cell cycle

**Figure S13: Transcriptional Target and Cell Cycle Gene Expression Following Loss of G0-skip Genes.**
**A**, Hierarchical clustering and percent overlap of differentially expressed up- and down-regulated genes between G0-skip KO and NTC G0/G1 U5-NSCs. Percent gene overlap is relative to the smaller gene set.

**B**, Heat map of high-confidence TP53 target genes (direct regulation score ≥ 10) (Fischer, 2017) that are significantly changed (white dots, FDR<0.05) in at least one G0-skip KO com- pared to NTC in G0/G1 U5-NSCs measured through RNA-sequencing. The reduction of TP53-dependent transcription is only seen in the TP53 KO.

**C**, Heat map of conserved YAP target genes (Cordenonsi et al., 2011) that are significantly changed (white dots, FDR<0.05) in at least one G0-skip KO compared to NTC in G0/G1 U5-NSCs measured through RNA-sequencing. There are unique subsets of Hippo-YAP target genes up-regulated in *NF2* and *TAOK1* KOs. Significance of overlap assessed through hyper-geometric analysis.

**D**, Venn diagram of YAP target overlap between G0/G1 U5-NSCs with G0-skip KO shows distinct gene sets up-regulated by NF2 (sgNF2-YAP) and TAOK1 (sgTAOK1-YAP) KOs as well as some shared targets. *PTPN14* KO is most similar to NF2 KO but shares some target up-regulation with *TAOK1* KO.

**E**, Gene ontology analysis of the sgNF2-YAP (yellow) and sgTAOK1-YAP (red) target subsets.

**F**, Heat map of the cell cycle genes (Biological process: G0:007049) significantly changed (FDR<0.05) in at least one G0-skip KO compared to NTC in G0/G1 U5-NSCs measured through RNA-sequencing. Subsets that are specifically up- or down-regulated in one or more KO after hierarchical clustering were designated by colored boxes. A subset of corresponding genes are highlighted below in color-corresponding boxes including DREAM complex components down in *NF2* KOs (LIN9, MYBL2, and RBL1) and some gene expression changes that were not consistent increased proliferation (i.e., up-regulation of CDK inhibitors CDKN2B/p16 and CDKN2B/p15 in NF2 and *PTPN14* KO, up-regulation of CDKN1A/p21 and TP73 in *CREBBP* KO, and others). In *NF2/PTPN14* KO boxes, N=significant in *NF2* KO only, P=significant in *PTPN14* KO only, and B = significant in both (FDR<0.05 compared to NTC).
All heat maps normalized by read counts per million (CPM) and scale depicts row-wise z-score.

### Figure S13

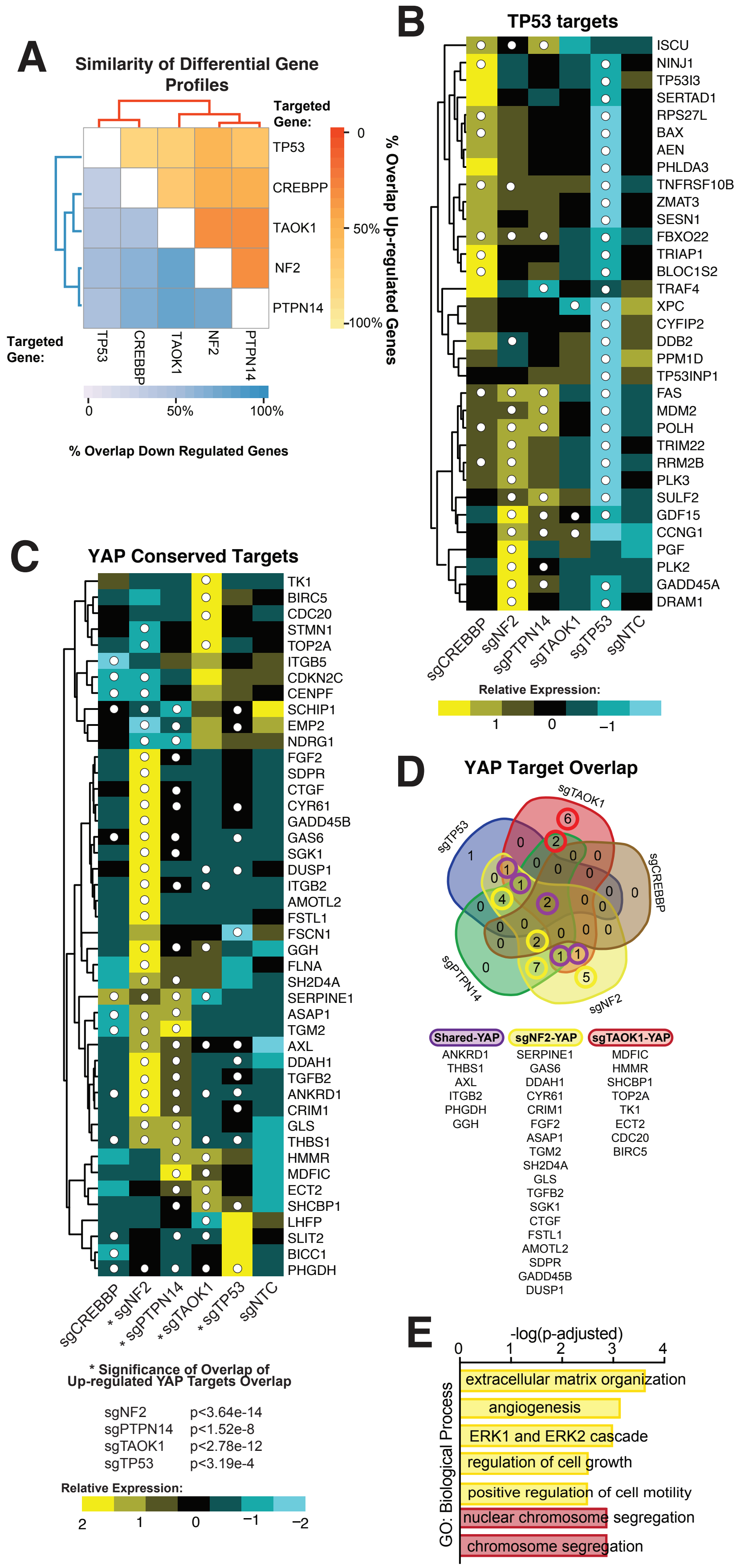

A
Similarity of Differential Gene Profiles
Targeted Gene:
TP53
CREBPP
TAOK1
NF2
PTPN14
Targeted Gene:
0
50%
100%
% Overlap Down Regulated Genes
% Overlap Up-regulated Genes
B
TP53 targets
ISCU
NINJ1
TP53I3
SERTAD1
RPS27L
BAX
AEN
PHLDA3
TNFRSF10B
ZMAT3
SESN1
FBXO22
TRIAP1
BLOC1S2
TRAF4
XPC
CYFIP2
DDB2
PPM1D
TP53INP1
FAS
MDM2
POLH
TRIM22
RRM2B
PLK3
SULF2
GDF15
CCNG1
PGF
PLK2
GADD45A
DRAM1
sgCREBBP
sgNF2
sgPTPN14
sgTAOK1
sgTP53
sgNTC
Relative Expression:
1
0
−1
C
YAP Conserved Targets
TK1
BIRC5
CDC20
STMN1
TOP2A
ITGB5
CDKN2C
CENPF
SCHIP1
EMP2
NDRG1
FGF2
SDPR
CTGF
CYR61
GADD45B
GAS6
SGK1
DUSP1
ITGB2
AMOTL2
FSTL1
FSCN1
GGH
FLNA
SH2D4A
SERPINE1
ASAP1
TGM2
AXL
DDAH1
TGFB2
ANKRD1
CRIM1
GLS
THBS1
HMMR
MDFIC
ECT2
SHCBP1
LHFP
SLIT2
BICC1
PHGDH
sgCREBBP
* sgNF2
* sgPTPN14
* sgTAOK1
* sgTP53
sgNTC
* Significance of Overlap of Up-regulated YAP Targets Overlap
sgNF2 p<3.64e-14
sgPTPN14 p<1.52e-8
sgTAOK1 p<2.78e-12
sgTP53 p<3.19e-4
Relative Expression:
2
1
0
−1
−2
D
YAP Target Overlap
sgTAOK1
sgTP53
sgCREBBP
sgPTPN14
sgNF2
Shared-YAP
ANKRD1
THBS1
AXL
ITGB2
PHGDH
GGH
sgNF2-YAP
SERPINE1
GAS6
DDAH1
CYR61
CRIM1
FGF2
ASAP1
TGM2
SH2D4A
GLS
TGFB2
SGK1
CTGF
FSTL1
AMOTL2
SDPR
GADD45B
DUSP1
sgTAOK1-YAP
MDFIC
HMMR
SHCBP1
TOP2A
TK1
ECT2
CDC20
BIRC5
E
-log(p-adjusted)
0
1
2
3
4
GO: Biological Process
extracellular matrix organization
angiogenesis
ERK1 and ERK2 cascade
regulation of cell growth
positive regulation of cell motility
nuclear chromosome segregation
chromosome segregation

**Figure S14: Gene Ontology Analysis of Genes Down Regulated in G0/G1 Following Loss of G0-skip Genes in hNSCs.**

**A**, Overlap of genes that are significantly down regulated following G0-skip KO in G0/G1 U5-NSCs compared to the Neural G0 single cell RNA-sequencing cluster.

**B,** Dot plot of selected enriched gene ontology (GO) biological processes following G0-skip KO in G0/G1 U5-NSCs for up (A) and down (B) regulated genes compared to NTC. Gene ratio com- pares the proportion of genes contained within the gene ontology group that are significantly changed in the G0-skip KOs to the total number of genes significantly changed in the G0-skip mutants.

**C,** Overlap of a selection of biological process enriched in down regulated genes in G0/G1 following KO of G0-skip genes using gene ontology analysis. Full list in Supplementary Table S7.

### Figure S14

A

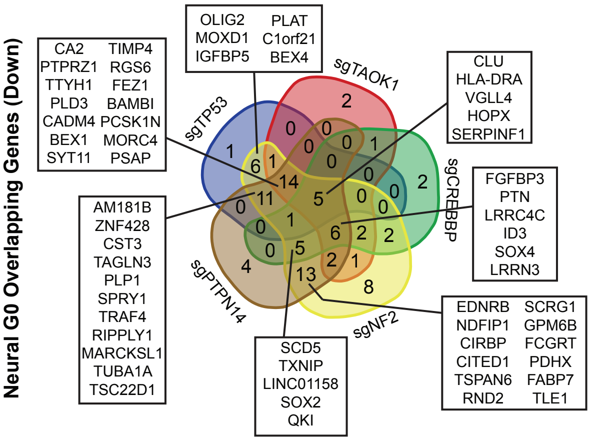


B

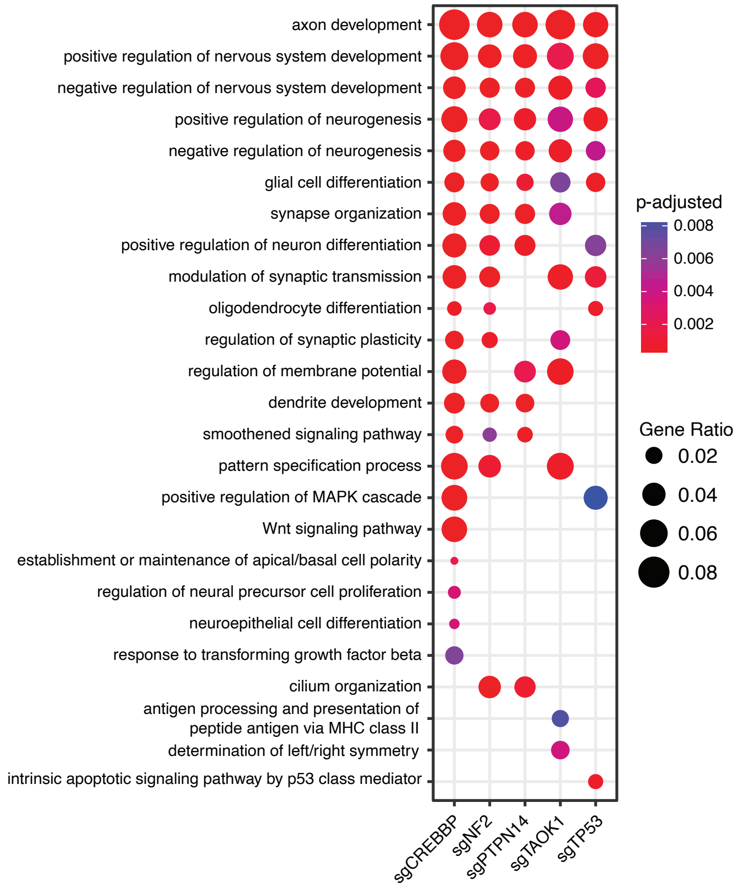


C

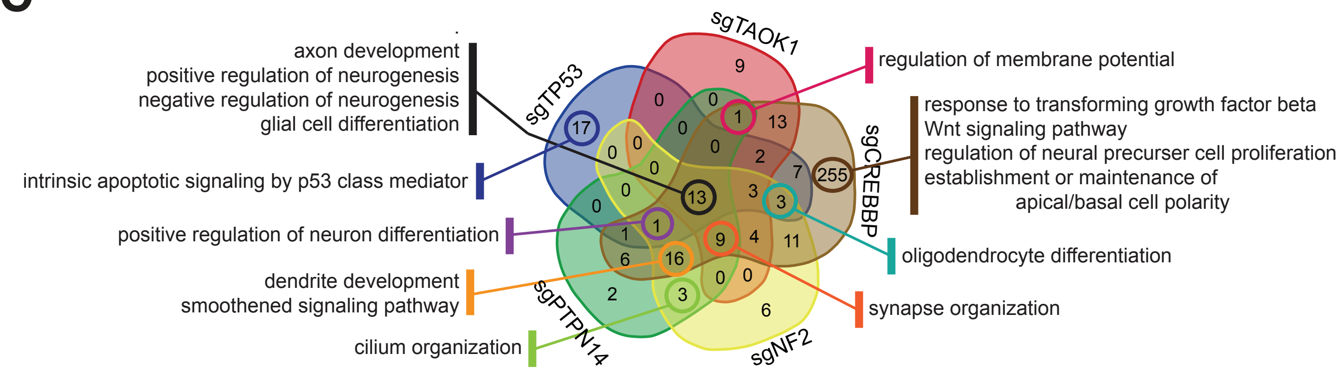

**Figure S15: Gene Ontology Analysis of Genes Up Regulated in G0/G1 Following Loss of G0-skip Genes in hNSCs.**
**A**, Overlap of genes that are significantly up following G0-skip KO in G0/G1 U5-NSCs compared to the Late G1 (B) single cell RNA-sequencing cluster.

**B,** Dot plot of selected enriched gene ontology (GO) biological processes following G0-skip KO in G0/G1 U5-NSCs for up regulated genes compared to NTC. Gene ratio compares the proportion of genes contained within the gene ontology group that are significantly changed in the G0-skip KOs to the total number of genes significantly changed in the G0-skip mutants.

**C,** Overlap of a selection of biological process enriched in up regulated genes in G0/G1 following KO of G0-skip genes using gene ontology analysis. Full list in Supplementary Table S7.

### Figure S15

**A**

**Late G1 Overlapping Genes**
**Up-regulated in G0/G1 sorted KO NSCs**

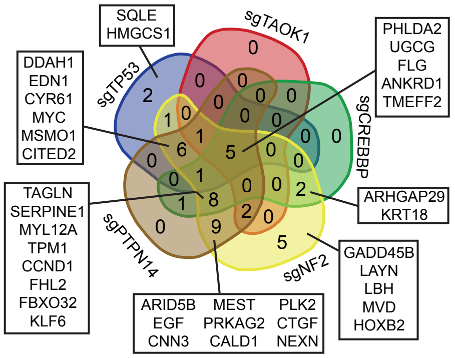


**B**

**G0/G1 Up-regulated GO Biological Processes**

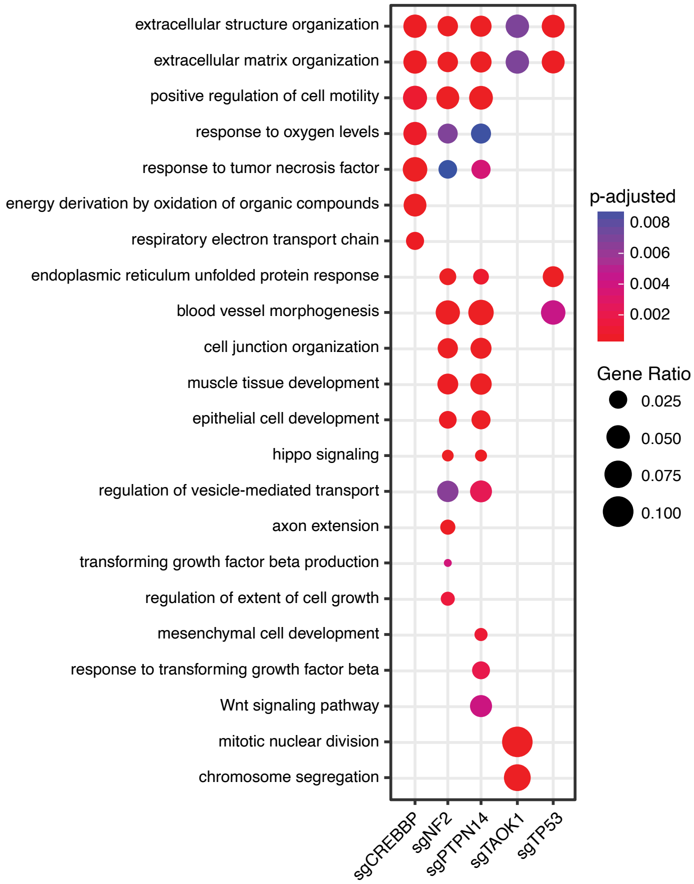


**C**

**Up-regulated Genes (G0/G1-sorted): Biological Processes**

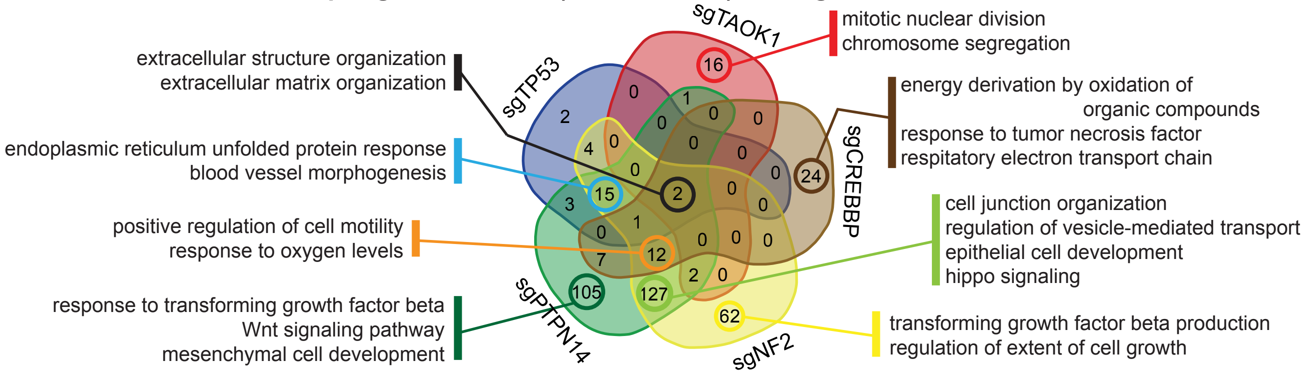

**Figure S16: hNSC cell cycle cluster genes altered in patient-derived GSCs.** Significance of overlap of the down (A) and up (B) regulated genes from bulk RNA-sequencing of G0/G1 sorted GSC-0131 and GSC-0827 cells with the single cell cluster definitions (up-regulated genes). Significance assessed though hypergeometric analysis. RF = representation factor.

### Figure S16

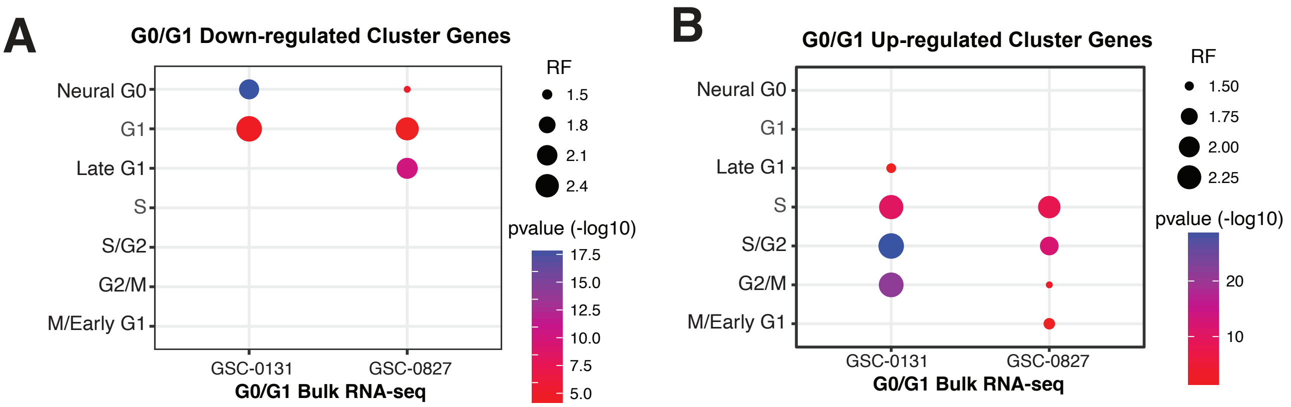
A
G0/G1 Down-regulated Cluster Genes
Neural G0
G1
Late G1
S
S/G2
G2/M
M/Early G1
GSC-0131
GSC-0827
G0/G1 Bulk RNA-seq
RF
1.5
1.8
2.1
2.4
pvalue (-log10)
17.5
15.0
12.5
10.0
7.5
5.0
B
G0/G1 Up-regulated Cluster Genes
Neural G0
G1
Late G1
S
S/G2
G2/M
M/Early G1
GSC-0131
GSC-0827
G0/G1 Bulk RNA-seq
RF
1.50
1.75
2.00
2.25
pvalue (-log10)
20
10

**Figure S17: Additional gene expression analysis of G0-skip mutants in hNSCs.**
**A-D**, Heat maps for genes (FDR<0.05) defining some of the significant biological process gene ontology groups in G0/G1 U5-NSCs (Table S7). (A) Cell cycle genes significant altered in each G0-skip mutant. (B) HIPPO signaling genes (GO:0035329), significantly enriched in *NF2* (p-adj.=3.52E-05) or *PTPN14* KO (p-adj.=9.42E-05). (C) Mitotic nuclear division genes (GO:007067), significantly enriched in *TAOK1* KO (p-adj.=3.16E-10). (D) Respiratory electron transport chain genes (GO:00229004), significantly enriched in *CREBBP* KO (p-adj.=1.62E-04).
All heat maps normalized to the regularized log (RLD). Scale applies to all heat maps and refers to the row-wise z-score. White dots indicate gene expression significantly different from NTC (FDR<0.05).

### Figure S17

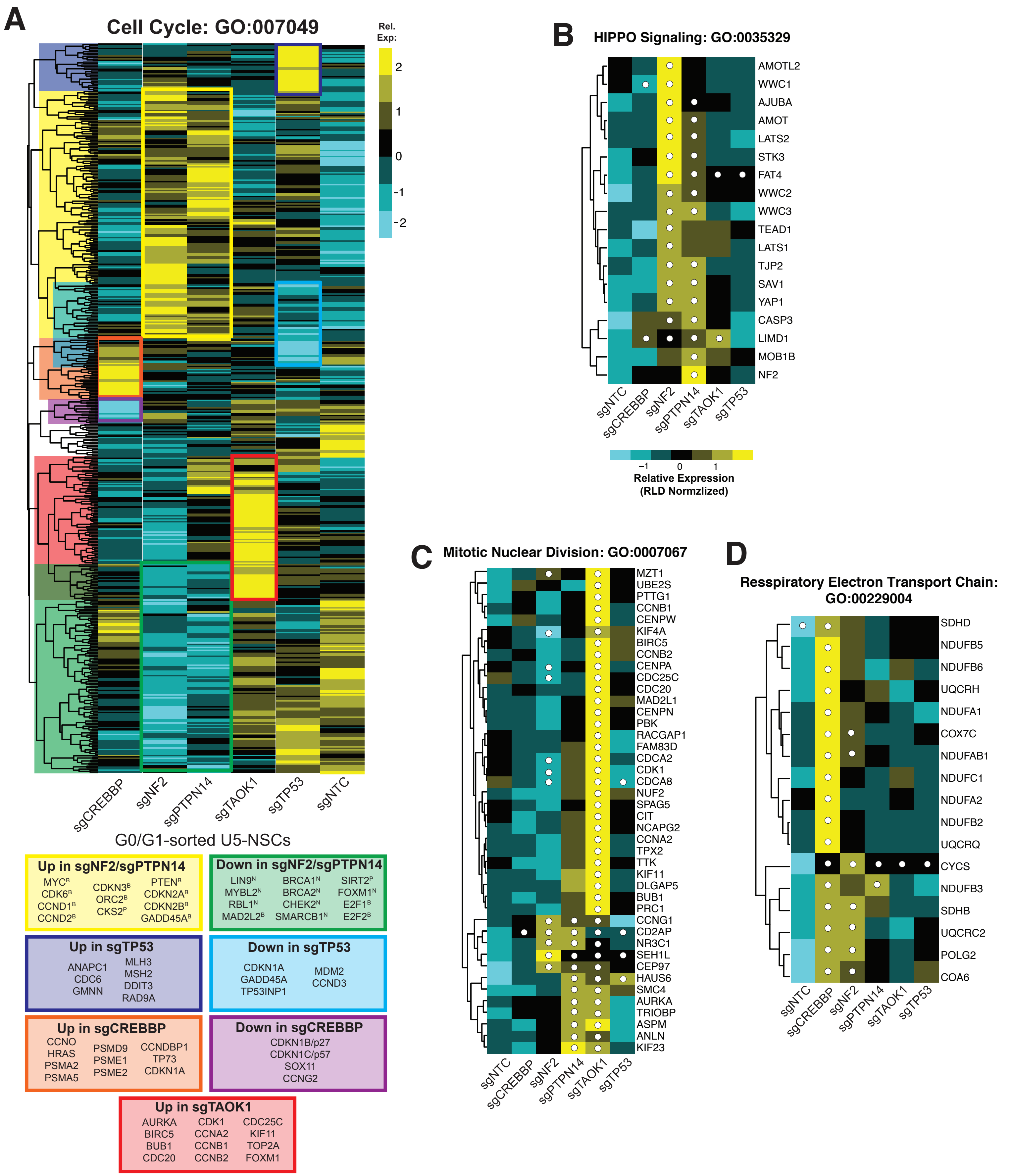
A
Cell Cycle: GO:007049
Rel. Exp:
2
1
0
-1
-2
sgCREBBP
sgNF2
sgPTPN14
sgTAOK1
sgTP53
sgNTC
G0/G1-sorted U5-NSCs
Up in sgNF2/sgPTPN14
MYC[B] CDK6[B] CCND1[B] CCND2[B] CDKN3[B] ORC2[B] CKS2[P] PTEN[B] CDKN2A[B] CDKN2B[B] GADD45A[B]
Down in sgNF2/sgPTPN14
LIN9[N] MYBL2[N] RBL1[N] MAD2L2[B] BRCA1[N] BRCA2[N] CHEK2[N] SMARCB1[N] SIRT2[P] FOXM1[N] E2F1[B] E2F2[B]
Up in sgTP53
ANAPC1 CDC6 GMNN MLH3 MSH2 DDIT3 RAD9A
Down in sgTP53
CDKN1A GADD45A TP53INP1 MDM2 CCND3
Up in sgCREBBP
CCNO HRAS PSMA2 PSMA5 PSMD9 PSME1 PSME2 CCNDBP1 TP73 CDKN1A
Down in sgCREBBP
CDKN1B/p27 CDKN1C/p57 SOX11 CCNG2
Up in sgTAOK1
AURKA BIRC5 BUB1 CDC20 CDK1 CCNA2 CCNB1 CCNB2 CDC25C KIF11 TOP2A FOXM1
B
HIPPO Signaling: GO:0035329
AMOTL2
WWC1
AJUBA
AMOT
LATS2
STK3
FAT4
WWC2
WWC3
TEAD1
LATS1
TJP2
SAV1
YAP1
CASP3
LIMD1
MOB1B
NF2
sgNTC
sgCREBBP
sgNF2
sgPTPN14
sgTAOK1
sgTP53
−1
0
1
Relative Expression
(RLD Normzlized)
C
Mitotic Nuclear Division: GO:0007067
MZT1
UBE2S
PTTG1
CCNB1
CENPW
KIF4A
BIRC5
CCNB2
CENPA
CDC25C
CDC20
MAD2L1
CENPN
PBK
RACGAP1
FAM83D
CDCA2
CDK1
CDCA8
NUF2
SPAG5
CIT
NCAPG2
CCNA2
TPX2
TTK
KIF11
DLGAP5
BUB1
PRC1
CCNG1
CD2AP
NR3C1
SEH1L
CEP97
HAUS6
SMC4
AURKA
TRIOBP
ASPM
ANLN
KIF23
sgNTC
sgCREBBP
sgNF2
sgPTPN14
sgTAOK1
sgTP53
D
Resspiratory Electron Transport Chain:
GO:00229004
SDHD
NDUFB5
NDUFB6
UQCRH
NDUFA1
COX7C
NDUFAB1
NDUFC1
NDUFA2
NDUFB2
UQCRQ
CYCS
NDUFB3
SDHB
UQCRC2
POLG2
COA6
sgNTC
sgCREBBP
sgNF2
sgPTPN14
sgTAOK1
sgTP53

**Figure S18: scRNA-seq of TAOK1 KO hNSCs.**
**A**, tSNE plots and relative cluster proportions from G0/G1 sorted (Fucci CDT1+) U5-NSCs for control and sgTAOK1 cells characterized based on cell cycle cluster definitions.

**B**,**C**, Proportion of cells in cycle clusters that are significantly different in G0/G1 sgTAOK1 vs. G0/G1 WT U5-NSCs after single cell RNA-sequencing. Significance was assessed using the proportion test.

### Figure S18

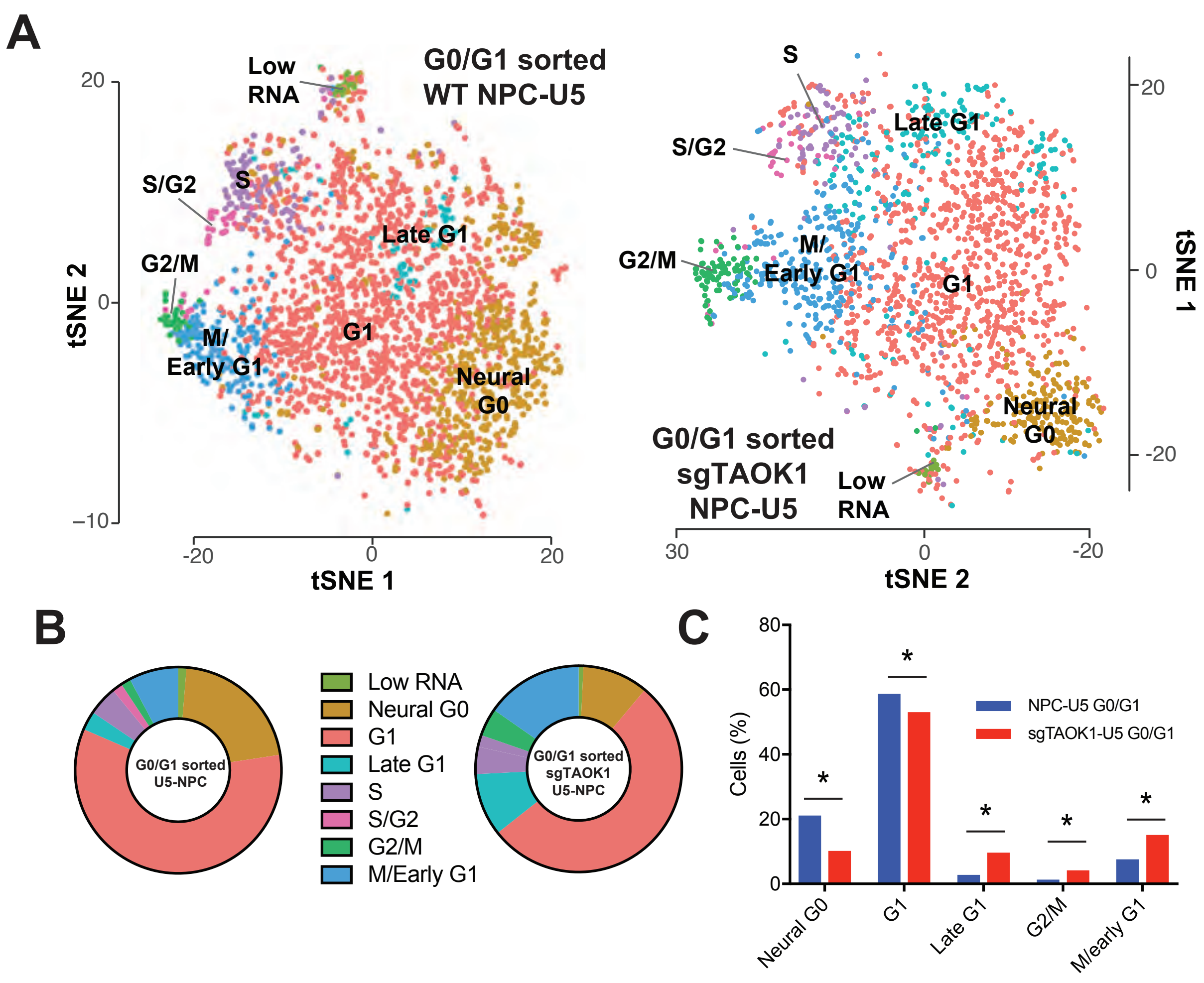

A
Low RNA
G0/G1 sorted WT NPC-U5
S/G2
S
Late G1
G2/M
M/ Early G1
G1
Neural G0
tSNE 2
tSNE 1
20
0
−10
-20
0
20
S
Late G1
S/G2
G2/M
M/ Early G1
G1
Neural G0
G0/G1 sorted sgTAOK1 NPC-U5
Low RNA
tSNE 2
tSNE 1
30
0
-20
20
0
-20
B
G0/G1 sorted U5-NPC
Low RNA
Neural G0
G1
Late G1
S
S/G2
G2/M
M/Early G1
G0/G1 sorted sgTAOK1 U5-NPC
C
Cells (%)
80
60
40
20
0
NPC-U5 G0/G1
sgTAOK1-U5 G0/G1
*
Neural G0
G1
Late G1
G2/M
M/early G1
