## Supplemental Table S8 for "Neural G0: a quiescent-like state found in neuroepithelial-derived cells and glioma"

**Table S8: List of Key Resources Including Antibodies, Validation sgDNA Sequences, and Primers**

***Antibodies***

| **Antibody Target (Use, dilution)** | **Source** | **Product Number** |
| --- | --- | --- |
| anti-CREBBP (WB, 1:500) | Cell Signaling | 7389 |
| anti-NF2 (WB, 1:200) | Santa Cruz | SC-332 |
| anti-Beta-Actin (WB,1:1000) | Cell Signaling | 3700 |
| anti-H4 (WB, 1:2,000) | Abcam | 17036-100 |
| anti-phosphorylated RB (Ser807/811) (IF, 1:1600) | Cell Signaling | 8516 |
| anti-Rabbit AF647 (2°, IF, 1:200) | Fisher | A21245 |
| anti-GFAP (IF, 1:1500) | Millipore | AB5804 |
| anti-β-tubulin III (TUJ1) (IF, 1:400) | Chemicon | MAB1637 |
| anti-Nestin (IF, 1:250) | Santa Cruz | sc-23927 |
| anti-Sox2 (IF, 1:200) | Cell Signaling | 3579S |
| anti-Mouse AF488 (2° IF, 1:200) | Fisher | A11001 |
| anti-Rabbit AF568 (2° IF, 1:200) | Fisher | A11011 |

WB: western blot; IF: immunofluorescence; 2°: secondary antibody

***Guide Sequences***

| **Guide Name** | **Sequence** |
| --- | --- |
| sgNTC | GTAGCGAACGTGTCCGGCGT |
| sgCREBBP-1 | AGCGGCTCTAGTATCAACCC |
| sgCREBBP-2 | GAATCACATGACGCATTGTC |
| sgCREBBP-3 | CCCGCGTGACCAGTCATTTG |
| sgCREBBP-4 | CAGCGAGCCTATGCTGCTCT |
| sgNF2-1 | ATTCCACGGGAAGGAGATCT |
| sgNF2-2 | AAACATCTCGTACAGTGACA |
| sgNF2-3 | CCTGGCTTCTTACGCCGTCC |
| sgNF2-4 | GTACTGCAGTCCAAAGAACC |
| sgPTPN14-1 | TGTTATCGAGTGCACGCTGT |
| sgPTPN14-2 | CTAGCCGGCCTAGCTGTGCA |
| sgPTPN14-3 | ATTACGATGTACATTGGACC |
| sgPTPN14-4 | CGTTGTAGCGCCGTGTCCGG |
| sgTAOK1-1 | ATTTACGTGAACACACAGCA |
| sgTAOK1-2 | TCTGCTTCGGATTTACTAGA |
| sgTAOK1-3 | CCCAACAGTATAGAATACAA |
| sgTAOK1-4 | GGAATAACATGTATTGAACT |
| sgTP53-1 | CCGGTTCATGCCGCCCATGC |
| sgTP53-2 | ACTTCCTGAAAACAACGTTC |
| sgTP53-3 | CATGTGTAACAGTTCCTGCA |
| sgTP53-4 | CGCTATCTGAGCAGCGCTCA |
| sgGNAS-1 (Control) | CCCCGAGAACCAGTTCAGAG |
| sgGNAS-2 (Control) | CAACCAGACCAACCGCCTGC |
| sgGNAS-3 (Control) | TGTGGCCGCCATGAGCAACC |
| sgGNAS-4 (Control) | TGTGACTGCCATCATCTTCG |
| sgKIF11-1 | TTCTTCTTCGCAGACGAATT |
| sgKIF11-2 | TTCGTCTGCGAAGAAGAAAG |
| sgKIF11-3 | TGCGAAGAAGAAAGAGGAGA |

***Primers***

| **Primer Name** | **Purpose** | **Sequence** |
| --- | --- | --- |
| PLC_Seq_R1_F | First Round Guide Amplification (Screens) | GAGGGCCTATTTCCCATGATTCCTTCA |
| PLC_Seq_R1_R | First Round Guide Amplification (Screens) | AACTTCTCGGGGACTGTGG |
| PLC_seq_R2_RS | Second Round Guide Amplification (Screens) | CAAGCAGAAGACGGCATACGAGATGTGACTGGAGTTCAGACGTGTGCTCTTCCGATCTTGCCACTTTTTCAAGTTGATAACGGACT |
| PLC-Seq-R2-barcode* | Second Round Guide Amplification (Screens) | AATGATACGGCGACCACCGAGATCTACACTCTTTCCCTACACGACGCTCTTCCGATCT**(BARCODE)**TCTTGTGGAAAGGACGAAACACCG |
| ArrayF | Validation Guide Assembly | TAACTTGAAAGTATTTCGATTTCTTGGCTTTATATATCTTGTGGAAAGGACGAAACACCG |
| ArrayR | Validation Guide Assembly | ACTTTTTCAAGTTGATAACGGACTAGCCTTATTTTAACTTGCTATTTCTAGCTCTAAAAC |

*Each sample receives its own barcode using a specific PLC-Seq-R2-barcode primer.
